# Distinguishing among modes of convergent adaptation using population genomic data

**DOI:** 10.1101/119578

**Authors:** Kristin M. Lee, Graham Coop

## Abstract

Geographically separated populations can convergently adapt to the same selection pressure. Convergent evolution at the level of a gene may arise via three distinct modes. The selected alleles can (1) have multiple independent mutational origins, (2) be shared due to shared ancestral standing variation, or (3) spread throughout subpopulations via gene flow. We present a model-based, statistical approach that utilizes genomic data to detect cases of convergent adaptation at the genetic level, identify the loci involved and distinguish among these modes. To understand the impact of convergent positive selection on neutral diversity at linked loci, we make use of the fact that hitchhiking can be modeled as an increase in the variance in neutral allele frequencies around a selected site within a population. We build on coalescent theory to show how shared hitchhiking events between subpopulations act to increase covariance in allele frequencies between subpopulations at loci near the selected site, and extend this theory under different models of migration and selection on the same standing variation. We incorporate this hitchhiking effect into a multivariate normal model of allele frequencies that also accounts for population structure. Based on this theory, we present a composite-likelihood-based approach that utilizes genomic data to identify loci involved in convergence, and distinguishes among alternate modes of convergent adaptation. We illustrate our method on genome-wide polymorphism data from two distinct cases of convergent adaptation. First, we investigate the adaptation for copper toxicity tolerance in two populations of the common yellow monkey flower, *Mimulus guttatus*. We show that selection has occurred on an allele that has been standing in these populations prior to the onset of copper mining in this region. Lastly, we apply our method to data from four populations of the killifish, *Fundulus heteroclitus*, that show very rapid convergent adaptation for tolerance to industrial pollutants. Here, we identify a single locus at which both independent mutation events and selection on an allele shared via gene flow, either slightly before or during selection, play a role in adaptation across the species’ range.

## 1 Introduction

Convergent adaptive evolution, where selection independently drives the evolution of the same trait, demonstrates the impressive ability of natural selection to repeatedly shape phenotypic diversity (Losos, 2011). Many studies have revealed cases of repeated adaptation resulting from changes in the same molecular mechanisms across distinct lineages (Stern, 2013; Wood et al., 2005). Here, we use the term convergence to define all cases of repeated evolution of similar traits across independent lineages, and do not distinguish between convergent and parallel evolution (Arendt and Reznick, 2008). In some cases, these convergent adaptive changes are identical at the level of the same orthologous gene or nucleotide (Martin and Orgogozo, 2013), suggesting adaptation may be more predictable and constrained than previously appreciated. Studying repeated evolution has long played a key role in evolutionary biology as a set of replicated natural experiments to help build comparative arguments for traits as adaptations, and to identify and understand the ecological and molecular basis of adaptive traits (Harvey and Pagel, 1991).

While we often think of convergent evolution among long-separated species, populations of the same (or closely-related) species often repeatedly evolve similar traits in response to similar selective pressures (Arendt and Reznick, 2008). Convergent adaptation at the genetic level among closely related populations may arise via multiple, distinct modes (see Stern, 2013, for a recent review). Selected alleles present at the same loci in multiple populations can have multiple independent mutational origins (e.g. Pearce et al., 2009; Chan et al., 2010; Tishkoff et al., 2007). Alternatively, adaptation in different populations could proceed by means of selection on the standing variation present in their ancestor (e.g. Colosimo et al., 2005; Roesti et al., 2014), or a single allele spread throughout the populations via gene flow (e.g. Heliconius Genome Consortium, 2012; Song et al., 2011). Understanding the source of convergent adaptation can aid in our understanding of fundamental questions about adaptation. Distinguishing among these modes may provide evidence for how restricted the paths adaptation can take are to pleiotropic constraints and if adaptation is limited by mutational input (Orr 2005, for review). Additionally, we can improve our understanding of the role of standing variation and gene flow in adaptation (Barrett and Schluter, 2008; Hedrick, 2013; Welch and Jiggins, 2014).

With the advent of population genomic data, it is now possible to detect genomic regions putatively underlying recent convergent adaptations. A growing number of studies are sequencing population genomic data from closely related populations, in which some have potentially converged on an adaptive phenotype (e.g. Turner et al., 2010; Jones et al., 2012). Population genomic studies of convergent evolution often take a paired population design, sampling multiple pairs of populations that independently differ in the key phenotype or environment. These studies are usually predicated on finding large effect loci which have rapidly increased from low frequency to identify the population genomic signal of selective sweeps shared across populations that independently share a selective pressure. Regions underlying convergent adaptations can potentially be identified by looking for genomic regions where multiple pairs of populations are strongly differentiated (e.g. using *F*_*ST*_) compared to the genomic background. Another broad set of approaches identify convergent loci by looking for genomic regions where the populations that share an environment cluster together phylogenetically in a way unpredicted by genome-wide patterns or geography (e.g. Pease et al., 2016; Jones et al., 2012). While these methods have proven useful in identifying loci involved in convergent adaptation, currently there are few model-based ways to identify the signal of convergence in population genomic data or to distinguish the different modes of convergent adaptation. In the case where an allele is shared due to adaptation from standing variation or migration, chunks of the haplotype on which the selected allele arose and swept on will also be shared among the populations (Slatkin and Wiehe, 1998; Bierne, 2010; Kim and Maruki, 2011; Roesti et al., 2014), providing a useful heuristic for these modes to be distinguished from convergent sweeps from independent mutations. We also note there are a variety of approaches to detect introgression (see Hedrick, 2013; Racimo et al., 2015; Rosenzweig et al., 2016, for recent reviews). However, these methods are not usually focused on detecting sweeps in both populations, but rather look for signatures of unusual amounts of shared ancestry between populations. Here, we present coalescent theory that leverages these signatures selection has on linked neutral variation in a model-based approach. We extend this to a statistical method that utilizes genomic data to identify loci involved in and distinguish between modes of genotypic convergence.

Positive selection impacts neutral diversity at linked loci due to hitchhiking (Maynard Smith and Haigh, 1974; Kaplan et al., 1989) and can be modeled as an increase in the variance in neutral allele frequencies around their ancestral frequencies. We develop coalescent theory to show how shared hitchhiking events between subpopulations act to increase covariance in allele frequencies around their ancestral frequencies at loci near the selected site, and extend this theory under different models of migration and selection on the same standing variation. We incorporate this hitchhiking effect into a multivariate normal model of allele frequencies that also accounts for population structure, allowing for the application to data from many populations with arbitrary relationships. Based on this theory, we present a composite-likelihood-based approach (Kim and Stephan, 2002; Nielsen et al., 2005; Chen et al., 2010; Racimo, 2016) that utilizes genomic single-nucleotide polymorphism (SNP) data to identify loci involved in convergence, and distinguish among alternate modes of convergent adaptation. As these models are also specified by relevant parameters, it is possible to obtain estimates for parameters of interest such as the strength of selection, the minimum age and frequency of a standing variant, and the source population of the beneficial allele in cases of migration. We also present a parametric-bootstrapping approach to help with model choice and construct confidence intervals for our parameters as standard likelihood approaches are not applicable to composite likelihoods.

This method should be of wide use with the increase in population genomic samples from across the geographic range of a species. Here, we illustrate the utility of our inference method by applying it to genome-wide polymorphism data from two distinct cases of convergent adaptation. First, we investigate the basis of the convergent adaptation observed across populations of the annual wildflower *Mimulus guttatus* to copper contaminated soils from two populations sampled near Copperopolis, California (Wright et al., 2015). We find selection has been acting on standing variation shared between these populations for a tolerance allele present prior to the onset of copper mining in this region. To further exemplify the flexibility of our method, we study a more complex population scenario: the rapid adaptation of four populations of killifish (*Fundulus heteroclitus*) to high levels of pollution, sampled across the Eastern seaboard of the United States (Reid et al., 2016). We find that even at the level of a single gene, both convergent mutation and selection on an allele shared via gene flow, either slightly before or during selection, have played a role in adaptation in this species.

## 2 Models

In the following section, we present models for the three modes of genotypic convergent adaptation: (1) multiple independent mutations at the same locus, (2) selection on shared ancestral standing variation, and (3) migration between populations spreading a beneficial allele (Figure 2). Throughout this section, we compare our derived expectations to coalescent simulations using mssel, a modified version of ms (Hudson, 2002) that allows for the incorporation of selection at a single site. This simulation program takes as input the frequency trajectory of the selected allele for each population. We simulate stochastic trajectories of the selected allele in populations following our three modes of convergence (see Appendix A.2 for simulation details). We focus on a set of four populations as shown in Figure 1 where populations 2 and 3 are adapted to a shared novel selection pressure and populations 1 and 4 are in the ancestral environment. The average coancestry coefficient values across simulations, estimated as described in Appendix A.1, are plotted for 100 bins of recombination distance away from the selected site, which occurs at distance 0. The results for all three models are shown in dashed lines in Figure 3.

**Figure 1:**
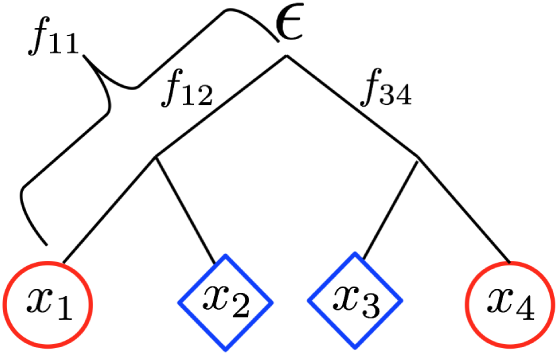
Present day population allele frequencies at a given neutral locus (*x*_1_–*x*_4_ for populations 1–4, respectively) are derived from ancestral allele frequency ϵ. Each population has a coancestry coefficient proportional to the amount of drift experienced since the split from the ancestral population. *f*_11_ is shown for population 1. Here, populations 1 and 2, and 3 and 4 share drift relative to the ancestral population and have nonzero coancestry coefficients *f*_12_ and *f*_34_, respectively. Blue diamonds represent the novel selective environment and red circles the ancestral environment. Note that branch lengths are not proportional to time in generations (unless there is no migration and the amount of drift is small).

**Figure 2:**
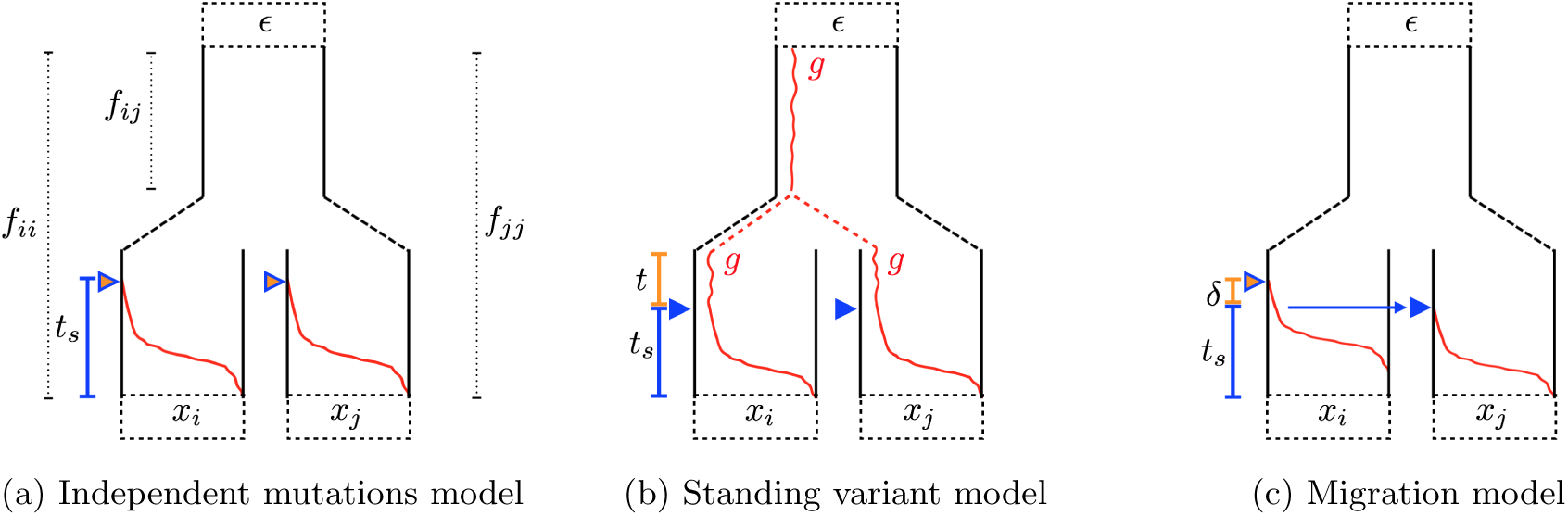
Trajectories of the beneficial allele (red) for the three modes of convergent adaptation. Populations *i* and *j* are under selection with present-day allele frequencies *x*_*i*_ and *x*_*j*_ at a neutral locus, derived from an ancestral population with allele frequency ϵ. The populations share some amount of drift proportional to *f*_*ij*_ before reaching the ancestral population. (2a) Beneficial mutations, indicated by the orange triangles, occur independently in the selected populations after they have become isolated. Selection begins, indicated by the blue triangles, once the beneficial allele is present in the population. The beneficial allele sweep to fixation in *t*_*s*_ generations. (2b) The beneficial allele is standing at frequency *g* in the ancestral population. After the selected populations split, it is still standing at frequency *g* for *t* generations prior to the onset of selection. (2c) The beneficial allele arises in population *i* and begins sweeping in population *i*. Meanwhile, there is a continuous low level of migration from population *i* into population *j*. The beneficial allele establishes in *j* after *δ* generations, where it is swept to fixation in *t*_*s*_ generations.

**Figure 3:**
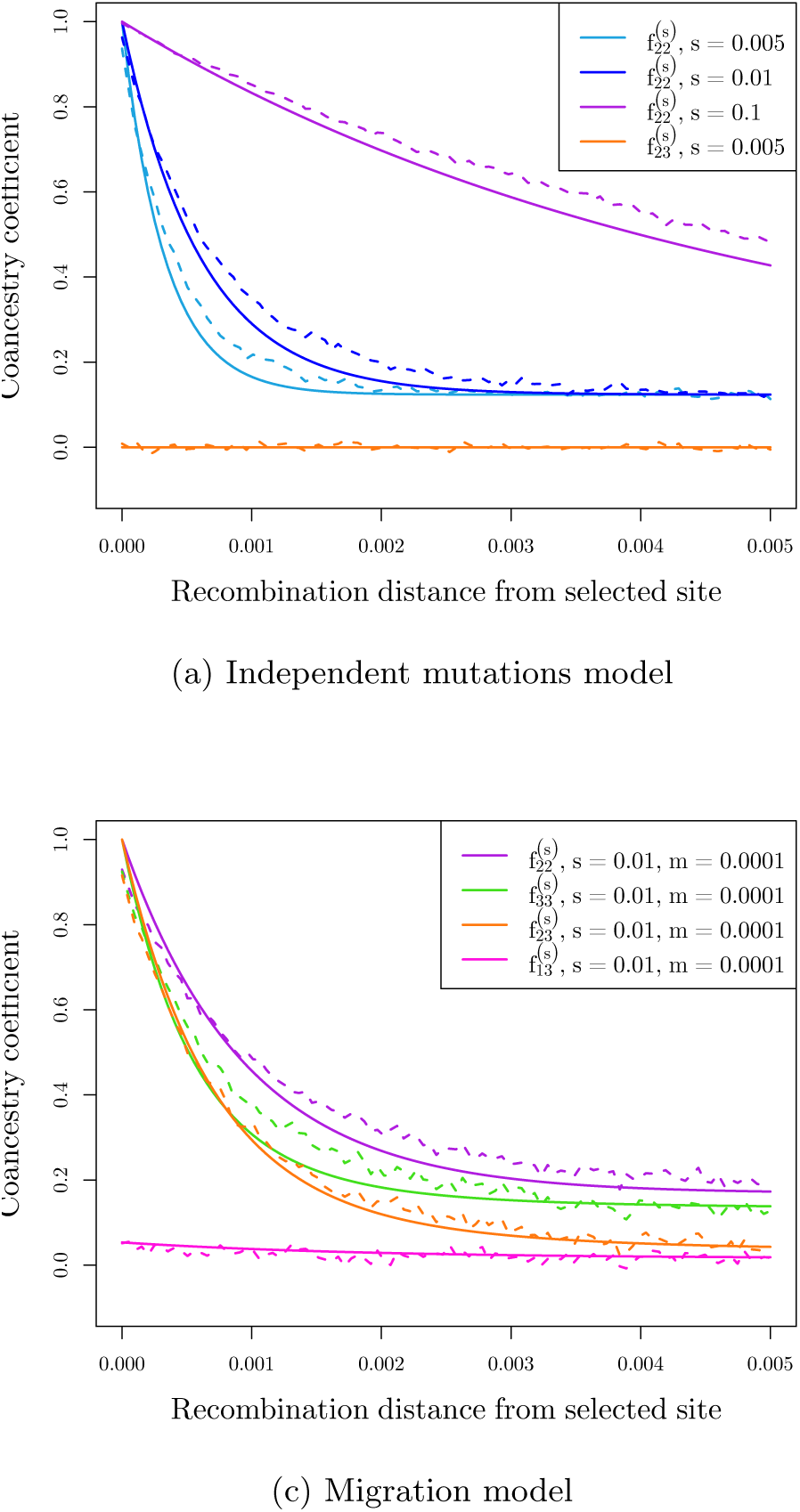
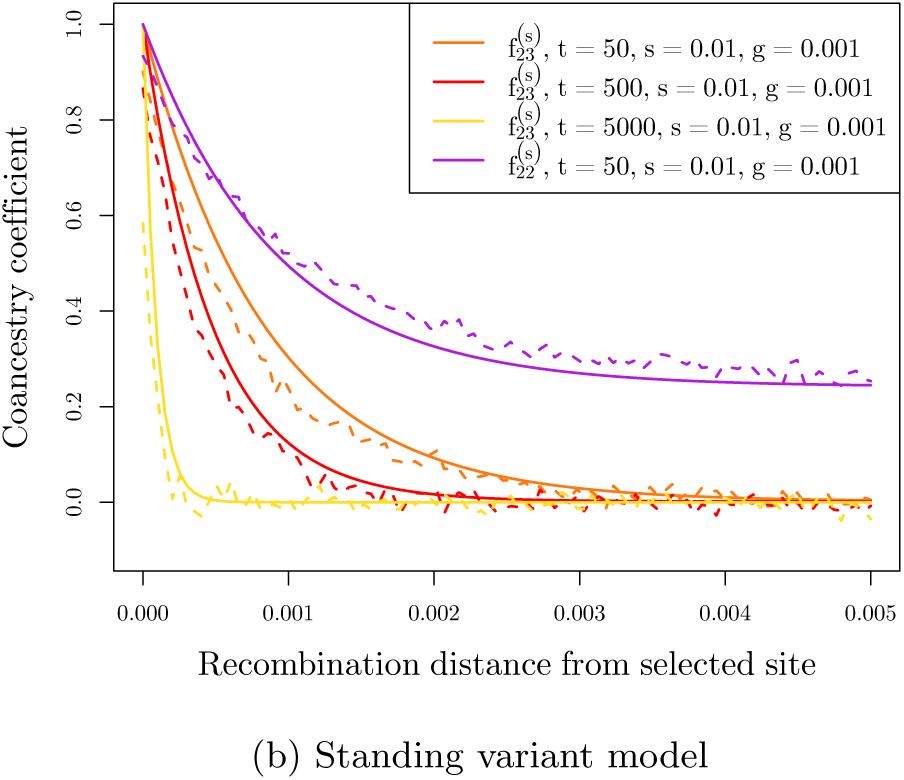
We calculated the average coancestry coefficient values across 1000 runs of simulations for each of 100 bins of distance away from the selected site to compare our simulation results (dashed lines) to our theoretical expectations (solid lines). (3a) Average coancestry coefficients under the independent mutations model (*N*_*e*_ = 100, 000) within a selected population (population 2) with varying *s*. Also shown is the coancestry coefficient between selected populations which in this case is 0, the neutral expectation. (3b.) Coancestry coefficients under the standing variation model between selected populations with varying amount of time beneficial allele has been independently standing in populations (*t*). The coancestry coefficient within a single population is also shown for *t* = 50. For all, *N*_*e*_ = 10, 000, *g* = 0.001*, s* = 0.01. (3c) Coancestry coefficients under the migration model, within both selected populations (source population 2 and recipient population 3) as well as between source and recipient (2,3) and between recipient and a non-selected population (1,3). Here we are showing one set of parameters (*s* = 0.01, *m* = 0.001, *N*_*e*_ = 10, 000) as estimates do not vary dramatically with changing *m* (see Figure S2).

### 2.1 Null Model

We aim to model the variances and covariances of the neutral allele frequencies within and between populations due to convergent sweeps. First, we must specify a null model that accounts for population structure. Populations will have some level of shared deviations away from an ancestral allele frequency, ϵ, due to shared genetic drift. Let *x*_*i*_ represent the present day allele frequency in population *i* (Figure 1). We denote the deviation of this frequency from the ancestral frequency by Δ*x*_*i*_ = *x*_*i*_-ϵ. Genetic drift, in expectation across loci, does not change the population allele frequencies (i.e. 𝔼[Δ*x*_*i*_] = 0) as an allele increases or decreases in frequency with equal probability. Drift however does act to increase the variance in this deviation across loci, with this variance increasing as more time is allowed for drift. The variance in the change of neutral allele frequencies in population *i* is

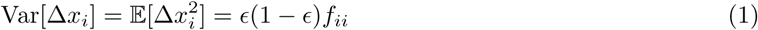

where *f*_*ii*_ can be thought of as the genetic drift branch length leading from the ancestral population to population *i* (Nicholson et al., 2002), specifying how much allele frequencies in population *i* deviate from their ancestral values (Figure 1). By rearranging Equation 1, *f*_*ii*_ can be interpreted as the population-specific *F*_*ST*_ for population *i* relative to the total population, here represented by the ancestral population (Wright, 1943, 1951; Weir and Hill, 2002; Nicholson et al., 2002).

Populations covary in their deviations from ϵas some populations are more closely related due to shared genetic drift resulting from shared population history or gene flow. The covariance in this deviation between populations *i* and *j* is

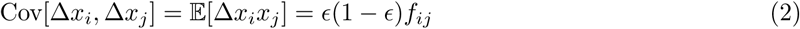

where *f*_*ij*_ is interpreted as the coancestry coefficient between populations *i* and *j*, and can be thought of as the shared branch length connecting *i* and *j* to the ancestral population (Figure 1).

Other natural interpretations of *f*_*ii*_ and *f*_*ij*_ follow from these definitions. Specifically, these values are probabilities of a pair of lineages being identical by descent relative to the ancestral population, i.e. the probability two sampled lineages coalesce before reaching the ancestral population (see Thompson, 2013, for a recent review). We briefly review this coalescent interpretation in Appendix A.1. For *f*_*ii*_ these two lineages are sampled both from population *i*. For *f*_*ij*_, one lineage is sampled from population *i* and the other from population *j*. We note that in practice we do not get to observe the ancestral frequency, nor may the history of our populations be well represented by a tree-like structure (for instance the history of our populations may be reticulated). However, for the sake of clarity, we proceed with these assumptions and deal with these complications in the implementation of the method.

We define a matrix, **F**, for *K* populations as a *K* × *K* matrix of coancestry coefficients. For example, for the four populations shown in Figure 1, this matrix takes the following form:

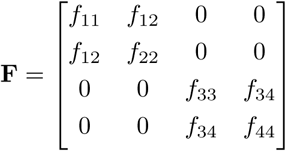

Populations *i* and *j* that split after the ancestral population and share no additional drift (e.g. populations 1 and 3) have *f*_*ij*_ = 0 by definition.

### 2.2 Incorporating selection

Positive selection impacts neutral diversity at linked loci due to hitchhiking. As the beneficial allele increases rapidly in frequency, so does the haplotype on which it arose. Neutral alleles further from the selected site may recombine off the selected background during the sweep, whose duration depends on the strength of selection (*s*) and weakly on the effective population size (*N*_*e*_). The effect of hitchhiking on the changes of linked neutral allele frequencies is similar to that of genetic drift. Hitchhiking does not alter the expected frequency change of linked neutral alleles across loci (i.e. 𝔼[Δ*x*_*i*_] = 0) because the selected mutation arises on a random haplotypic background. Moreover, hitchhiking increases the variance in the deviation in neutral allele frequencies away from their ancestral values (Var[Δ*x*_*i*_]) at linked sites (Gillespie, 2000). Shared hitchhiking events between subpopulations will act to increase covariance in allele frequency deviations between subpopulations (Cov[Δ*x*_*i*_, Δ*x*_*j*_]) at loci near the selected site. This effect of hitchhiking on linked diversity, within and among populations gives us a way to distinguish among alternate modes of convergent adaptation.

We define new matrices of coancestry coefficients that incorporate selection in addition to drift as **F**^(*S*)^. In the following section, we use a coalescent approach to derive coancestry coefficients within and between populations, 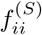 and 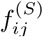, for the three modes of genotypic convergent adaptation (Figure 2). In Supplement S2 we derive some of the same results forwards in time to help guide the reader’s intuition. For all models, we assume the beneficial allele has gone to fixation in all selected populations recently. Note that all our models of selection are phrased in terms of distortions to the neutral matrix **F**; therefore, the precise source of the neutral population structure (e.g. whether its due to shared population history or migration) is relatively unimportant to our approach. A deeper knowledge of the basis of this structure does add to the interpretation of the results, as we explain in the discussion.

#### 2.2.1 Independent mutation model

We first consider the case when a beneficial allele arises independently via *de novo* mutations at the same locus, or tightly linked loci, in both of the selected populations. We expect hitchhiking to increase the variance in neutral allele frequency deviations around the selected site in both populations. However, as the sweeps are independent and there is no gene flow between populations during or after the sweep, we expect no covariance in the neutral allele frequency deviations between these populations, beyond that expected under neutrality due to shared population history prior to the introduction of the beneficial allele.

Moving backward in time, sampled neutral lineages linked to the selected site will be forced to coalesce if both lineages do not recombine off the sweep. We define the probability that a single neutral allele fails to recombine off the background of the beneficial allele during the sweep phase as *y*, which we can approximate as

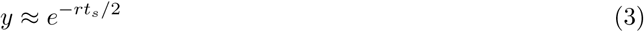

(Kim and Stephan, 2002; Durrett and Schweinsberg, 2004; Nielsen et al., 2005) where *r* is the recombination rate between the neutral locus and selected site, and *t*_*s*_ is the amount of time the sweep phase takes (Figure 2a). When the beneficial allele arises from a new mutation and selection is additive, *t*_*s*_ *≈* 2*log*(4*N*_*e*_*s*)/*s*, where *s* is the selection coefficient for the heterozygote, such that heterozygotes experience a selective advantage of *s* and homozygotes 2*s* (Gillespie, 2000; Barton, 1998). The factor of 4*N*_*e*_*s* is due to the fact that our new mutation, if it is to establish in the population, rapidly reaches frequency 1/(4*N*_*e*_*s*) in the population and then increases deterministically from that frequency (Maynard Smith, 1971; Barton, 1998; Kim and Stephan, 2002; Kim and Nielsen, 2004).

The coancestry coefficient in population *i* that experiences a sweep, 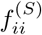, is defined as the probability that two lineages sampled from population *i* coalesce either due to the sweep phase or neutrally before reaching the ancestral population. With probability *y*^2^, both lineages fail to recombine off the beneficial background during the sweep, and they will be forced to coalesce. If one or both lineages recombines off the sweep (with probability 1 − *y*^2^), they can coalesce before reaching the ancestral population with probability *f*_*ii*_. Combining these we find

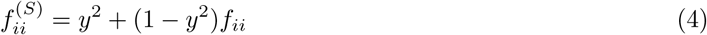

For convenience, in our inference procedure, we assume the same strength of selection between our selected populations and thus duration of the sweep is the same. So, 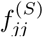 takes the same form as Equation 4, with its own neutral probability (*f*_*jj*_) of coalescing. Given that we assume the sweeps complete recently and have the same duration, the mutational events occur at approximately the same time in each selected population. If we assume there is no neutral migration amongst populations, Equation 4 will hold regardless of where the sweep occurs on the branch leading to *i* (but when migration occurs we need the sweep to be recent so that lineages sampled from population *i* are found in population *i* when the sweep occurs).

For the coancestry coefficient between two selected populations *i* and *j*, we can calculate the probability two lineages, one sampled from population *i* and the other from population *j*, coalesce. When the sweeps are independent, the lineages can only coalesce with probability *f*_*ij*_ before reaching the ancestral population, as they have no probability of coalescing during the sweep phases which have independent origins. Thus,

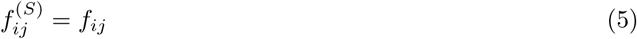

**Comparison to simulated data** In Figure 3a we show the case of convergence due to independent origins of the beneficial allele. As we predicted, there is no additional coancestry between the selected populations. Additionally, we show how the coancestry within a selected population decays with distance from the selected site for a range of values for the strength of selection. These coancestry values decay to the neutral expectation at other regions of the genome. With larger *s*, this decay is slower as the sweep occurs more rapidly and there are fewer chances for recombination to occur during this time.

#### 2.2.2 Standing variant model

We turn now to the case of a sweep shared between populations *i* and *j* due to selection acting on shared ancestral variation (Figure 2b). Our model is appropriate for cases where the standing variation from which the sweep arises was previously neutral or was maintained in the population at some low frequency by balancing selection. Let the beneficial allele be standing at frequency *g* in the ancestral population. We assume that the beneficial allele frequency does not deviate much from that of the ancestral population such that it is still *g* in the daughter populations prior to selection. Selection favoring the beneficial alleles begins *t* generations after the populations split and the beneficial allele reaches fixation in both populations after *t*_*s*_ generations (see Figure 2b). We assume *t*, *g*, and *s* are the same for all of our selected populations. More work is needed to allow population-specific parameters to relax these assumptions. We acknowledge all selected populations starting from the same beneficial allele frequency may be unrealistic in many cases, particularly if *t* is long or if the populations experience bottlenecks at the time of the split.

We first consider the coalescent process of two lineages within a single selected population. Again, *y* is the probability that a neutral lineage fails to recombine off the background of the beneficial allele during the sweep phase. Given that the beneficial allele is increasing from frequency *g*, *y* takes the same form as Equation 3, where now *t*_*s*_ ≈ 2 log(1/*g*)/*s*. If both lineages fail to recombine off the beneficial background during the sweep, there is a probability of coalescing during the standing phase that is higher than the probability of two neutral lineages randomly sampled from the population coalescing. Following from our assumptions during the standing phase, the rate at which two lineages coalesce within a population is 1/(2*N*_*e*_*g*) per generation. Alternatively, a lineage can recombine off in the standing phase onto the other background with probability *r*(1 − *g*) ≈ *r* per generation. As these are two competing exponential processes, the probability two lineages coalesce before either recombines off the beneficial background can be simplified to

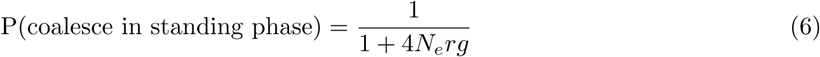

as described by Berg and Coop (2015). If either neutral lineage recombines off the beneficial background before they coalesce, the probability of coalescing with the other lineage before reaching the ancestral population can be treated as the coancestry coefficient associated with that particular portion of the population tree.

Taking these approximations into account, we derive a coancestry coefficient for a neutral allele in population *i* that experiences selection from standing variation as

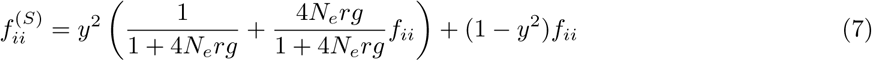

The first term corresponds to both lineages failing to recombine off the beneficial background during the sweep phase, which puts them both on the same background as the beneficial allele in the standing phase. Now, the two lineages can either coalesce in the standing phase or recombine off of the background of the beneficial allele where they can coalesce neutrally before they reach the ancestral population. Alternatively, one or both lineages can recombine off during the sweep phase and again they can coalesce neutrally.

Populations that share a sweep due to shared standing ancestral variation will have increased covariance in the deviations of neutral allele frequencies around their ancestral means around the selected site since they will have a shared segment of the swept haplotype. From a coalescent perspective, this occurs because two lineages sampled from each population have a higher probability of coalescing if they stay on the beneficial background during the sweep and standing phases than two lineages sampled randomly between the populations.

The probability that a single lineage does not recombine off onto the non-beneficial background during the standing phase for *t* generations can be approximated as

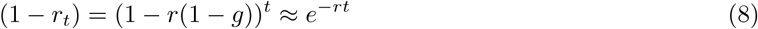

The coancestry coefficient between populations *i* and *j* is now

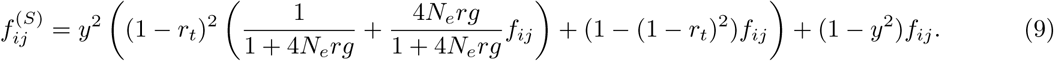

This derivation follows from that of 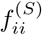in Equation 7, but now incorporates the additional probability (1−*r_t_*)^2^ of both lineages failing to recombine off the beneficial background during their independent standing phases for time *t*.

This standing variation case represents a simple model of selection on standing variation However, we expect in many cases that the beneficial allele has not been standing since the ancestral population of the convergent population, but rather has been moved among populations by migration before becoming adaptive at some later time point. In these cases we invoke a model where the standing allele spreading by migration from some source population to recipient populations *t* generations in the past before the allele became favored. See Appendix A.4 for details. This model differs from the migration model presented in the next section in which we assume a continuous rate of migration throughout the duration of the sweep and that the variants sweep as soon as they are established in the population. In this standing case with a source of the standing variant, moving backwards in time we assume that the allele is standing for *t* generations in a population after the sweep and before the beneficial lineage migrates back instantly into a specified source population (see Figure 11). Biologically, it naturally captures the case where the allele is shared between the populations due to migration but is standing for sometime before it sweeps. For data analysis, we default to using this more complex model, where sampled selected populations are evaluated as possible sources of the standing variant.

**Figure 11:**
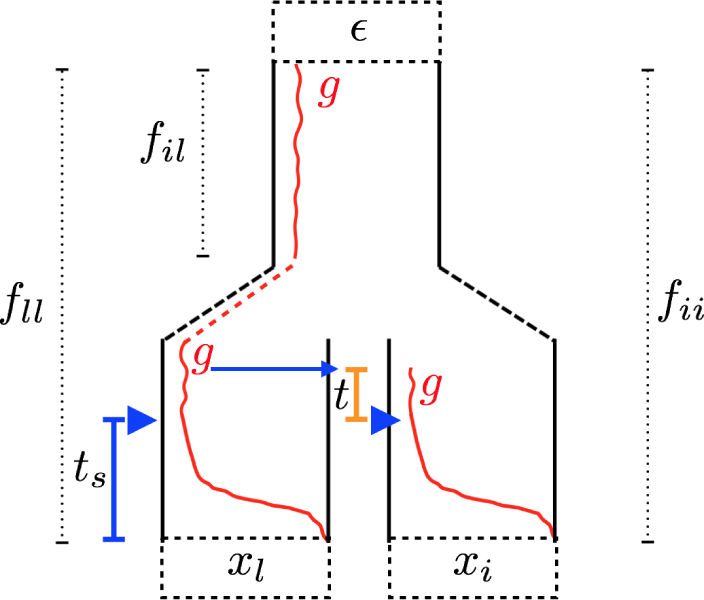
Trajectories of the beneficial allele (red) for the standing variant model with a source population. Populations *l* and *i* are under selection with present-day allele frequencies *x*_*l*_ and *x*_*i*_ at a neutral locus, derived from an ancestral population with allele frequency ∊. The populations share some amount of drift proportional to *f*_*il*_ before reaching the ancestral population. The beneficial allele is standing at frequency *g* in the source population, *l*. It migrates into population *i* from *l*,where it is standing at frequency *g* for *t* generations prior to the onset of selection, indicated by the blue triangles.

Extending this models to allow for the source to be a non-sampled population would be useful in studying the so-called “the transporter hypothesis” (Schluter and Conte, 2009; Bierne et al., 2013; Welch and Jiggins, 2014) where adaptive gene flow is acting to introduction variation standing in another population. Here, more work is needed to address issues related to estimating coancestry coefficients for unsampled populations (see Appendix A.4 for more information).

**Comparison to simulated data** In Figure 3b we show comparisons of simulations to show the fit of our predictions to simulations with adaptation from standing variation in the classic sense. As the duration of the independent standing phases, *t*, increases, the coancestry at linked neutral alleles between selected populations decreases. Forward in time, this has the interpretation that the longer the beneficial allele is standing in the populations, the shorter the shared haplotype between the populations will be due to independent recombination events before selection begins. In the case that the beneficial allele has been standing for a very long time (*t* → *∞*) before selection occurs, this additional covariance will reduce to zero as in the independent sweeps case (Equation 5). We acknowledge this scenario is biologically unrealistic. For large values of *t* at small *g*, we expect it is likely that the allele would get either be lost or there may be allelic turnover due to recurrent mutations of the beneficial allele. However, it is useful here to gain intuition about when our models overlap. Conversely, if the standing variant is very young (*t* → 0), the decay in covariance between populations takes the form of the variance within populations (Equation 7) which, as we will see in the next section, looks similar to the pattern generated under the migration model.

#### 2.2.3 Migration model

We now consider the case where the selected allele is spread across sub-populations by migration. This scenario has been studied by a number of authors (Slatkin and Wiehe, 1998; Santiago and Caballero, 2005; Kim and Maruki, 2011, note these all assume that the allele sweeps in all of the populations), and our approach here follows similar lines to that of Kim and Maruki (2011). Let there be a single origin of the beneficial allele, which occurs in population *i*. We assume a low, continuous level of migration during the sweep, with a proportion *m* of individuals in population *j* coming from population *i* each generation. Here we are considering only unidirectional migration from population *i* into population *j*. We say the sweep began in population *j* at time *t*_*s*_ generations in the past and at time *t*_*s*_ + *δ* for population *i* (Figure 2c). Kim and Maruki (2011) found that the mean delay time, *δ*, between the two sweeps can be approximated by

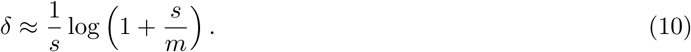

The coancestry coefficient of the source population, 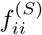, follows that of a population experiencing an independent sweep from new mutation (Equation 4). To derive the coancestry coefficient of the recipient population, 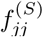, we first need to consider the fate of two lineages sampled in population *j* at the selected site. Two events can occur if we trace the lineages of two beneficial alleles back in time: either the two lineages coalesce in population *j* and a single lineage migrates back into population *i* or the two lineages independently migrate back into the source population and coalesce there. We define the probability of these two events as *Q* and 1 − *Q*, respectively. We use the approximation

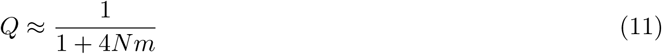

(see Pennings and Hermisson, 2006). Assuming *m* is small, such that a beneficial allele sampled at present day in population *j* migrates back into population *i* approximately *t*_*s*_ generations in the past, the probability of a linked neutral allele recombining off during the sweep phase in population *j* can be approximated by *y*. If the lineage migrates back into population *i* before it recombines off the beneficial background, there is an additional time *δ* in population *i* for recombination to happen. So, there is an additional probability, *e*^−*rδ*^, of recombination of our linked neutral allele off the beneficial background.

Thus, the coancestry coefficient for the recipient population is now

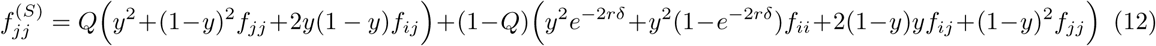

The terms in this approximation correspond to the following coalescent scenarios: First, if two lineages sampled in population *j* coalesce before migrating (with probability *Q*), then linked neutral alleles can coalesce either during the sweep if neither lineage recombines off the beneficial background, neutrally in population *j* if both lineages recombine off, or neutrally shared drift phase of populations *i* and *j* if just one lineage recombines off. Alternatively, if the two lineages fail to coalesce before one or both migrates (w.p. 1 − *Q*), there are four ways linked neutral alleles can coalesce:

1. Both lineages fail to recombine off the beneficial background during the sweep and are forced to coalesce during the sweep in population *i*. The factor *e*^-*2*^*rδ* represents the additional opportunity for recombination when both lineages have migrated back into population *i*.
2. Both lineages stay on the beneficial background in population *j* (w.p. *y*^2^) but one or both lineages recombines off in population *i* (w.p. 1 − *e*^−*2*^*rδ*) and they coalesce neutrally in the source population with probability *f*_*ii*_ before reaching the ancestral population.
3. Either lineage recombines off the beneficial background while it is still in population *j* and the two lineages coalesce neutrally in the shared drift phase of populations *i* and *j*, with probability *f*_*ij*_ before reaching the ancestral population.
4. Both lineages recombine off during the sweep phase while they are still in population *j* and they coalesce neutrally with probability *f*_*jj*_.

When a beneficial allele is shared between populations *i* and *j* via migration, there will be additional covariance in the deviations of linked neutral allele frequencies from their ancestral means. In this case, there are three ways a lineage sampled from population *i* and a lineage sampled from population *j* can coalesce. They are forced to coalesce during the sweep if both lineages fail to recombine off the background of the sweep, which occurs with probability *y*^2^*e*^−*rδ*^. Alternatively, the lineage sampled in population *j* can recombine off the beneficial background before it migrates back to source population *i*, in which case the lineages can coalesce neutrally before reaching the ancestral population in their shared drift phase, with probability *f*_*ij*_. Lastly, if the lineage sampled in population *j* migrates back into population *i* then the two sampled neutral lineages can coalesce neutrally in population *i* with probability *f*_*ii*_ if the lineages don’t coalesce due to the sweep (i.e. either recombines off in time *t*_*s*_ or *δ*). Thus, in the case of continuous migration the coancestry coefficient between the source and recipient population is

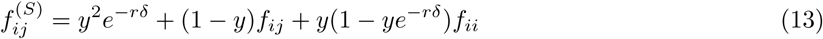

To fully specify the coancestry matrix with selection, we need to take into account the effect migration has on non-selected populations. Specifically, the coancestry coefficients between recipient and non-selected populations are impacted since there is some probability linked neutral lineages will migrate from the recipient population into the source population backwards in time. Let population *k* be a non-selected population. Now, the coancestry coefficient between populations *j* and *k* can be expressed as

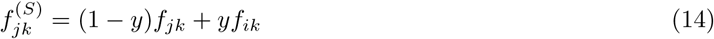

This is informative about the direction of migration. First, there is no impact of selection on the relationship between the source and non-selected populations. Additionally, the sweep shared via migration will induce additional coancestry between *j* and *k* if *k* is more closely related to our source population (e.g. population 1 in Figure 1 if population 2 is the source). The opposite is true if *k* is more closely related to our recipient population (e.g. population 4). Now, there is a deficit in the background level of coancestry between populations *j* and *k* near the selected site.

**Comparison to simulated data** In Figure 3c we show our results above compared to simulations with continuous migration during the sweep phase, for a single set of parameters (*s* = 0.01, *m* = 0.001). Here, we have migration occurring from population 2 into population 3. We show the four relevant coancestries as a function of distance from the selected site: the covariance within source 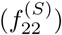, within recipient 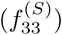, between source and recipient 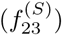, within recipient and a non-selected population 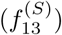. We see the coancestry within the recipient population decays more rapidly than coancestry within the source population. This fits our expectations as there is some probability a lineage will, backwards in time, migrate back to the source population, decreasing the probability of coalescing before reaching the ancestral population when *m* is small. As *m* increases, this relationship changes (Figure S2). We also see increased coancestry near the selected site between the selected populations. The pattern of decay varies from that observed in our standing variation model, except for when *t* is small. Additionally, we see increased coancestry between the recipient population and a non-selected population that decays with recombinational distance to their neutral expectation. Note, the reverse, coancestry recovering to the neutral expectation with recombinational distance is observed for populations that initially are more related to the recipient population (i.e. population 4), is also seen (Figure S3a). The coancestries between the source population and non-selected populations are unaffected (Figure S3b). Together, these observations using information from non-selected populations help distinguish possible source populations.

## 3 Inference

We have described how selection at linked loci affects the matrix of coancestry coefficients, allowing us to parameterize the variance and covariance in neutral allele frequency deviations within and between populations. To estimate the likelihood of our data under convergent adaptation models, we need a probability model for how allele frequencies depend on these variances and covariances. Neutral allele frequencies across *K* populations can approximately be modeled jointly as a multivariate normal distribution around the ancestral allele frequency, ϵ, with covariance proportional to the coancestry coefficients (Nicholson et al., 2002; Weir and Hill, 2002; Coop et al., 2010; Samanta et al., 2009). Specifically,

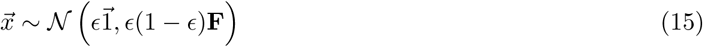

where 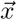 is a vector of population frequencies and **F** is the *K* by *K* matrix of coancestry coefficients without selection.

Above we demonstrated that we can generate coancestry matrices **F**^(*S*)^ to explain the coancestry between multiple populations due to neutral processes and various modes of convergent adaptation. **F**^(*S*)^ is a function of the neutral coancestry, (**F**) the model of convergence (*M*) and its parameters (Θ_*M*_), and the recombination distance a neutral site is away from a selected site (*r*_*l*_). Thus, modeling neutral allele frequencies as multivariate normal with covariance proportional to this new coancestry matrix, we can calculate the likelihood of observed data a given distance away from the selected site under a specific model of convergence as

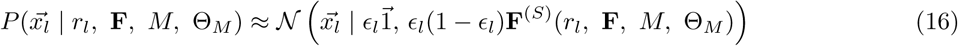

In practice, we do not know the true ancestral mean at a given locus, ϵ_*l*_, so we use the mean of the present day population allele frequencies and calculate likelihoods of mean-centered allele frequencies and coancestry matrices (we account for this mean centering in appendix A.2.6). We also do not know the true neutral coancestry matrix, **F**, but estimate it from deviations of allele frequencies from sample means across the entire genome. We also incorporate the effects of sampling into this variance-covariance matrix. See Appendix A.1 for details.

### 3.1 Composite-likelihood framework

We calculate the likelihood of all data (*D*_*𝓁*_) in a large window around the selected site (*𝓁*) under a given model of convergent adaptation (*M*), with its associated parameters (Θ_*M*_), as the product of the marginal likelihoods for sites all distances away from the selected site. This composite likelihood is used as an approximation to the total likelihood of all sites, but is not a proper likelihood as neighboring sites are correlated due to shared histories. Moving L_left_ sites to the left of the proposed selected site and L_right_ sites to the right,

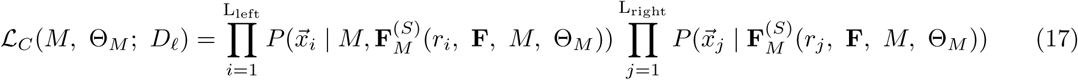

where *r*_*i*_ is the genetic distance from site *i* to *𝓁*, and similarly for *r*_*j*_. We can also obtain a composite likelihood of our data under a neutral model (*N*), ℒ_*C*_ (*N*; *D*_*𝓁*_), which is only parameterized by **F**. This framework enables us to:

1. Identify the maximum likelihood location of the selected locus in a region by varying the location of the proposed selected site. For a given region and model of convergent adaptation we vary the location of the selected site, taking the maximum composite likelihood over a grid of parameters. We take as our best estimate of the location under a given model of convergence, the maximum composite-likelihood location of the selected site 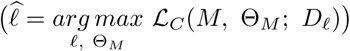.
2. Determine the parameter(s) which maximize our composite-likelihood estimates under a given model at a given location of the selected site (*𝓁*). We obtain these maximum composite-likelihood estimate (MCLE) parameters by evaluating the composite likelihood across a grid of parameters for a given location of the selected site 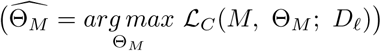
3. Distinguish between modes of convergence, and neutrality, in a genomic region by comparing the maximum likelihood under various models of convergent evolution. At a given location of the selected site (*𝓁*) we compare the maximum composite likelihood of each model to the neutral model 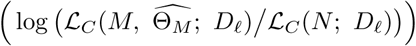

This composite likelihood ignores the correlation in allele frequencies (linkage disequilibrium) between neutral sites so the composite-likelihood surface will be too peaked. A number of authors have taken composite-likelihood approaches to inferring a range of population genetic parameters (e.g. Hudson (2001); see Larribe and Fearnhead (2011); Varin et al. (2011) for a broader statistical views on composite likelihood). In the setting of inferring genome-wide parameters, e.g. parameters of neutral demographic models, the maximum composite-likelihood parameter estimates are known to be consistent in the limit of many unlinked genomic regions (Wiuf, 2006). While in general composite-likelihood methods perform well, in all of these settings typical measures of uncertainty of parameters (confidence intervals) and model choice methods (e.g. AIC) are undermined due to the over peakiness of the likelihood.

Composite-likelihood approaches have also been used in the context of selective sweeps, starting with Kim and Stephan (2002) who take a composite likelihood formed like Equation 17 of the product of marginal probabilities of allele frequencies within a single population moving away from a proposed selected site (an approach expanded on by Kim and Nielsen, 2004; Nielsen et al., 2005; Chen et al., 2010; DeGiorgio et al., 2014; Racimo, 2016). Our method is most closely related to that of Chen et al. (2010) and Racimo (2016) who look at allele frequencies across two or three populations respectively, and look for the signal of a sweep in one of the populations (or in the case of Racimo, 2016, in the ancestor of a pair of populations). We note that we have a further layer of abstraction over these previous composite-likelihood methods. Extending Kim and Stephan (2002), previous methods have calculated the likelihood of the sample frequency considering a binomial draw from some underlying population frequency, which is naturally modeled as being bounded between 0 and 1. We, however, use a multivariate normal likelihood to model our sample frequencies, which does not bound allele frequencies between 0 and 1. This further abstraction is justified by the fact that by using the multivariate normal approach we are able to handle arbitrarily large number of populations with arbitrary population structure and to flexibly model different forms of selection into an easily extendable form to the covariance matrix. Future work could potentially concentrate on hybrid approaches, combining the flexibility of our approach with the realism of previous approaches.

### 3.2 Inference method on simulated data

To test our method, we utilized the datasets generated using mssel (as discussed above with details in Appendix A.2) to see if we could recover the parameters and convergent mode used for simulation. The neutral coancestry matrix **F** was estimated using data from 1000 runs with no selection (as described in Appendix A.1). We assume that the model parameters *N*_*e*_ and *r* are known and we set these at the values used to generate the simulations. We calculated the composite log-likelihoods for each of the simulated datasets under the following four models: neutral (no selection), independent sweep model, standing variation model, and migration model with the beneficial allele originating in population 2. We calculate the likelihoods under a dense grid of selection coefficients (*s*), migration rates (*m*), and standing times (*t*). In the standing variation model, the standing frequency (*g*) is held at 0.001. See Appendices A.2.4 and A.2.5 for details. We repeat this procedure for each of 100 runs of all simulated datasets. To compare between models, we calculate the composite log-likelihood differences between the true model and all other models including the neutral model, at the maximum composite-likelihood parameter estimate (MCLE) obtained under each model.

#### 3.2.1 Parameter estimation

**Location of selected site** To explore our method’s ability to localize the selected site, we vary the true location of the selected site simulating under the independent mutation model. We estimate the maximum composite-likelihood location under the independent sweep model over a fine grid of locations and selection coefficients. The method is able to correctly identify the location of selection (Figure 4a), with higher accuracy when the true location of the site is in the middle of the window. The method does show an edge effect when the true location of the selected site is at the edge of the region of interest perhaps because we do not get to see the decay of coancestry on both sides of the selected site. Additionally, we are able to correctly estimate the strength of selection while allowing the location of the selected site to vary (Figure S1a) and there is no correlation between these joint parameter MLCEs (Figure S1b).

**Figure 4:**
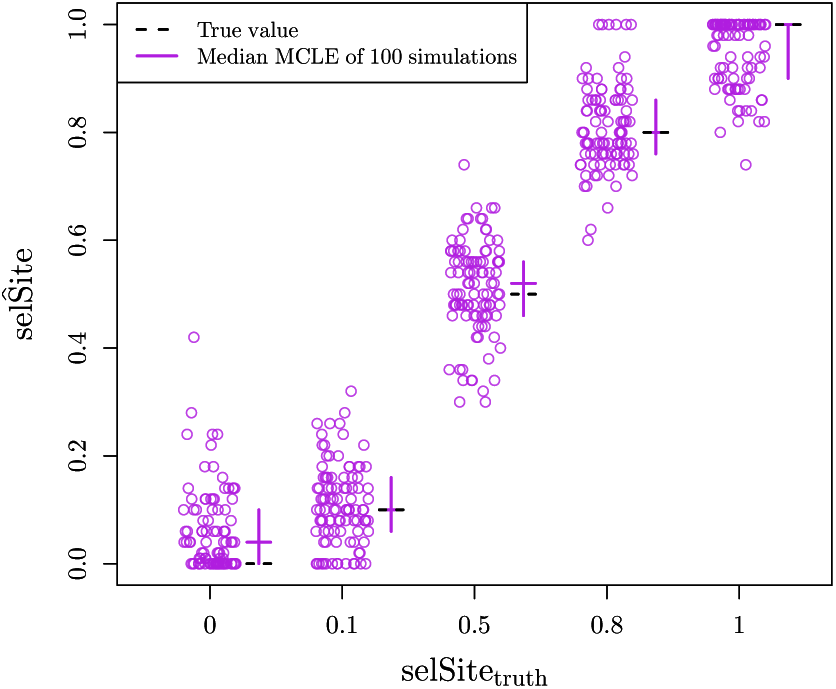
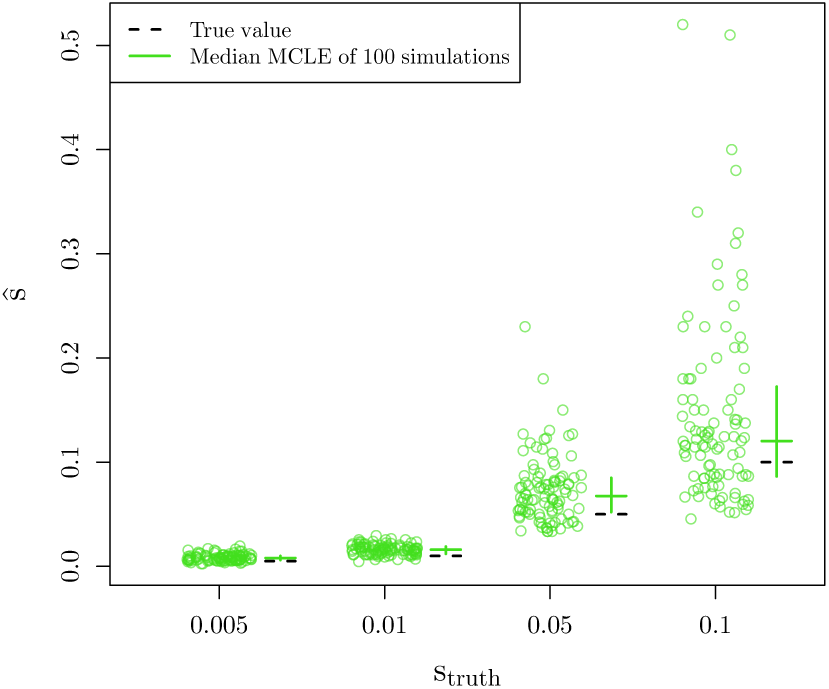
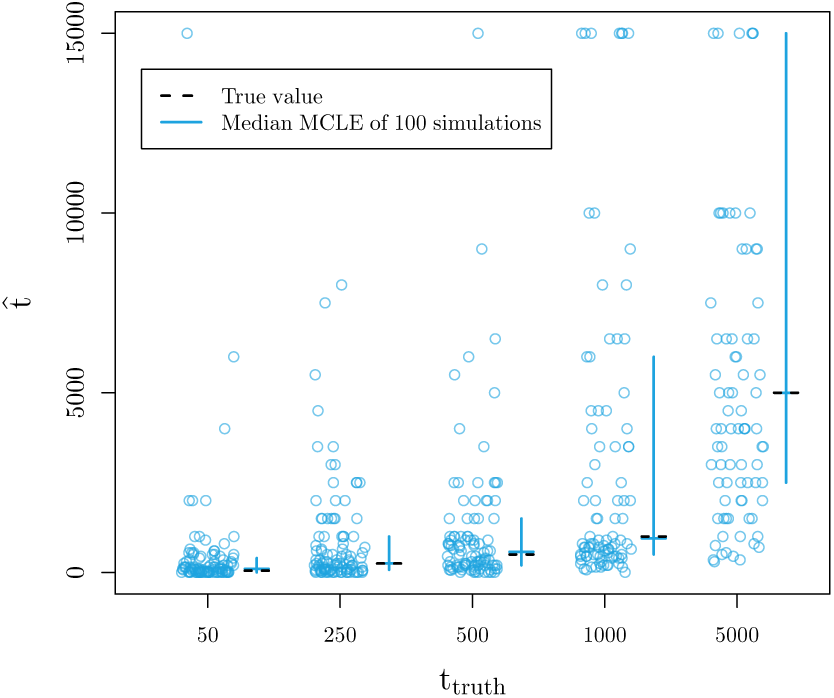
Maximum composite likelihood **parameter estimates** calculated under **model used for simulation**. We vary the true value of the parameter used for simulations along the x-axis and show the MCLE for each of 100 simulations (points). Crossbars indicate first and third quartiles with second quartiles (medians) as the horizontal line. The true values of the parameters are marked with dashed, black lines. (a) MCLE of the **location of selected site** for 100 simulations under the **independent mutation model** (10 chromosomes per population, *Ne* = 100,000, *s* =0.05) (b) MCLE of the **strength of selection** (*s*) for 100 simulations under the **independent mutation model** (10 chromosomes per population, *Ne* = 100,000) (c) MCLE of the **standing time** (*t*) for 100 simulations under the **standing variant model** (10 chromosomes per population, *Ne* = 10,000, *s* = 0.01, *g* = 0.001). For scale, we left out estimates of *t* > 15,000 (2, 9, and 21 data points when *t*_truth_ = 500, 1000, and 5000, respectively.)

**Independent mutations model** To verify our ability to recover the selection coefficient, we simulated under the independent mutation model for a range of values for *s*, holding the location of the selected site at its true value. We are able to recover the parameters used for simulation (Figure 4b). The ability to correctly estimate *s* breaks down for large enough *s*, given a fixed window-size around the selected site and *r*_*BP*_, since we will not observe the full decay in coancestry.

**Standing variant model** To explore our inference using the standing variant model, we hold the location of the selected site at its true location and take as our estimate of *s* and *t* their values at the joint maximum composite likelihood. Under the standing variant model, we are again able to accurately estimate *s* (Figure *S6). The inference of s* and *g* simultaneously is somewhat more confounded (Figure 5). How the signal of the sweep within populations decays, as we move away from the selected site, is primarily determined by *s* and *g* (see Equation 7). While a higher frequency of the standing variant (*g*) can lead to a quicker decay, this can be partially compensated for the strength of the sweep being stronger (higher *s*, lower *t*_*s*_). This explains the *J*-shaped ridge in the likelihood surfaces for *s* and *g*, seen in Figure 5. Therefore, in practice we can often infer a lower bound *s* and an upper bound for *g*, but not find the precise values of each when inference is performed under the standing variation model. We are able to accurately estimate the time the beneficial allele has been standing in the independent populations prior to selection, *t*, as shown in Figure 4c. Our inference of *t* is relatively free of confounding with *s* and *g*, as *t* primarily governs the decays in coancestry between populations, making it separable from the scale of the sweep within populations.

**Figure 5:**
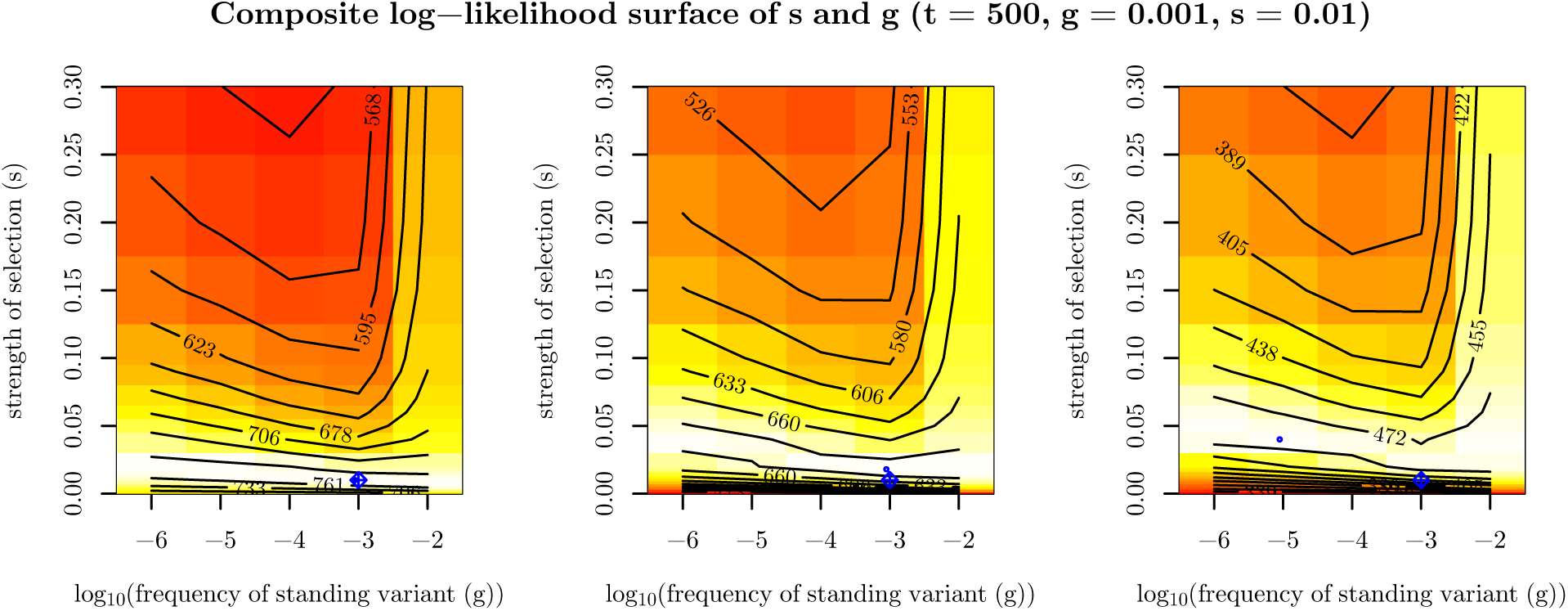
Composite log-likelihood surface of the **strength of selection** (*s*) and the **frequency of standing variant** (*g*) for three simulations (with *N*_*e*_ = 10,000, *t* = 500, *g* = 0.001, *s* = 0.01) to exemplify confounding of *s* and *g* under the **standing variant model**. Blue diamond pluses represent the true location of the parameters used for simulation. Blue circles represent MCLE.

**Migration model** We explored our inference under the migration model of parameters *m* and *s*, again fixing the location of the selected site and taking the joint maximum composite-likelihood estimate. We are able to correctly estimate *s* (Figure S4b). However, we obtain poor estimates of the rate of migration*, m* (Figure S4a). This is perhaps unsurprising as the coancestry coefficients under the migration model depend only weakly on *m*. We obtain fairly bimodal estimates of *m* that are usually either very low (10*^-5^* to 10*^-3^*) or high (1). As the true value of *m* increases, we see fewer estimates of small *m* and more estimates of *m* = 1. These estimates of *m* seem to be a true reflection of the patterns in the simulated datasets. Specifically, this effect is mostly observed in the variance within the recipient population as Equation 12 depends on *m* in both *Q* and *δ*. High *m* estimates correspond to datasets with lower empirical levels of coancestry within the recipient than datasets where low estimates of *m* were obtained (Figure S5). We believe that the bimodality results from stochasticity in how many lineages ancestral to the sample migrate before they recombine off the sweep in the recipient population. While our estimates of *m* are noisy, the migration model does capture key features of the spread of adaptive alleles by migration, allowing it potentially to be distinguished from other modes of convergence. We now turn to the performance of the method in distinguishing modes of convergence.

#### 3.2.2 Model comparison

To test the ability of our method to distinguish between modes of convergence, we calculated the maximum composite log-likelihood of 100 simulations for each dataset generated under both the true model and all other models with a fixed, fine grid of parameter values. The location of the selected site is fixed at its true location. The results are summarized in Figure 6, which shows histograms of the difference in maximum composite log-likelihoods calculated under a given model relative to the true model used for simulation. For example, in evaluating the independent mutations model, we present the difference in the composite loglikelihoods calculated for data simulated under the independent mutations model for all other models and the composite log-likelihood calculated for the true independent mutations model. Thus, values less than zero indicate that the correct model has a higher maximum composite log-likelihood than the true model. Conversely, values greater than zero indicate the incorrect model of convergence has a higher composite log-likelihood than the true model. For inference under the migration model, we fix the source to be the true source of the selected allele when simulating under the migration model, and to an arbitrary one of the two selected populations when performing inference on simulations under other models.

**Figure 6:**
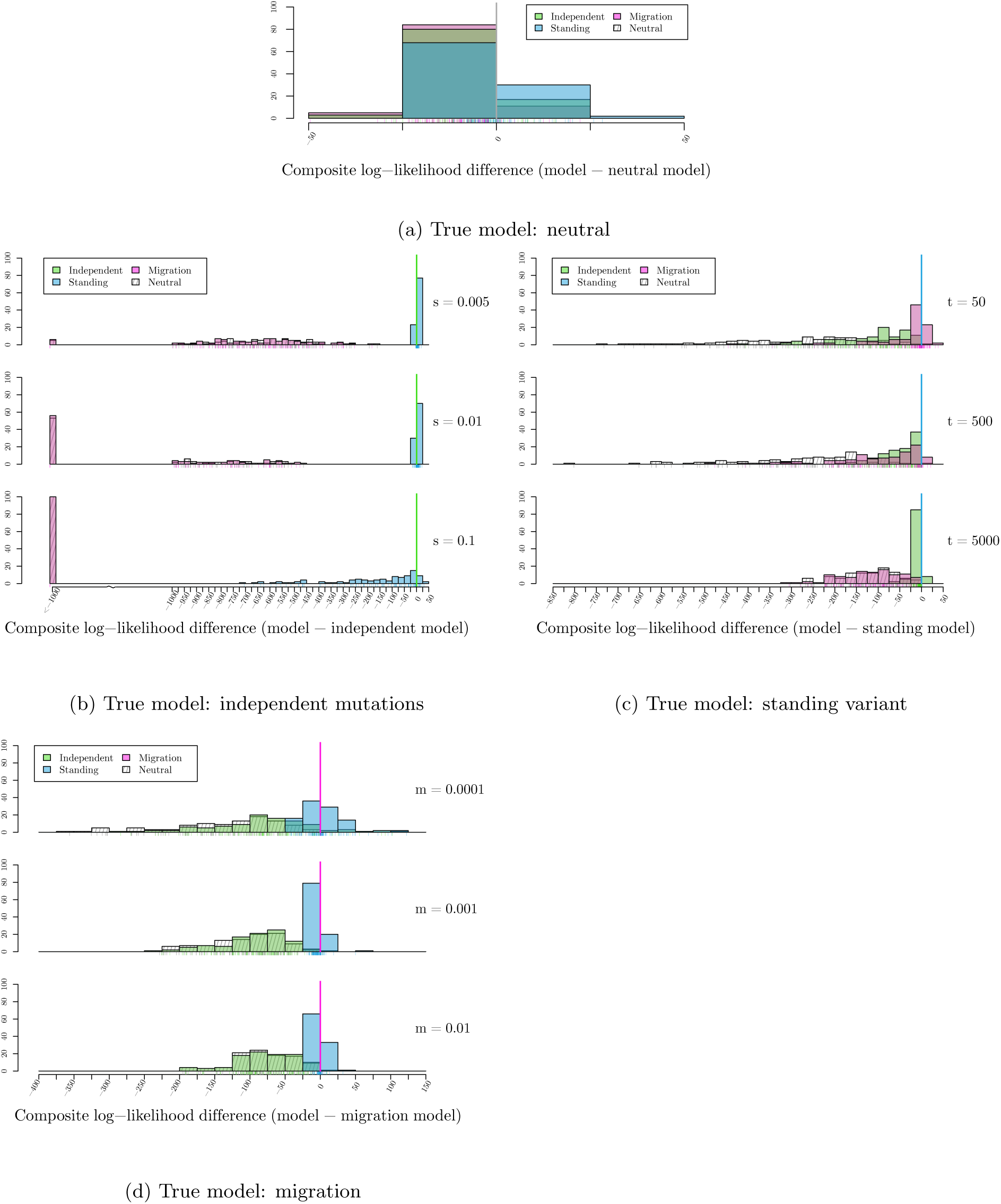
Histograms of the **differences in maximum composite log-likelihoods** calculated under a given model relative to the true model used for 100 simulations. Parameter values used to simulate are noted, varying along the vertical dimension. Values less than zero, marked with solid line, indicate the true model has a higher maximum composite likelihood than alternative model. Conversely, values greater than zero indicate the alternative, incorrect model of convergence has a higher composite log-likelihood than the true model. For (6b) *N*_*e*_ = 100, 000, (6c) *N*_*e*_ = 10, 000, *s* = 0.01, *g* = 0.001, (6d) *N*_*e*_ = 10, 000, *s* = 0.01. (a) True model: neutral (b) True model: independent mutations (c) True model: standing variant (d) True model: migration

**Neutral model** We first compare the composite likelihoods calculated for data generated with no selection. For the selection models, we fix the location of the selected site. The distributions of the resulting composite log-likelihood ratios are shown in Figure 6a. As expected for a composite likelihood, the composite loglikelihood ratio between a convergent selection model and the neutral model with no selection are inflated compared to those expected under the usual asymptotic *χ* ^2^ distribution. However, these likelihood ratio differences are relatively small compared to those we observed when simulating under alternative models. This is because when *s*→ 0 in all models with selection, the coancestries converge to our neutral expectations. Indeed when we look at the MCLE for the strength of selection (*ŝ*) under the incorrect models with selection, we see that for all nearly simulations *ŝ* is close to zero 0 (Figure 7a). Overall, this suggests that our null model is reasonably well calibrated, given the limitations of composite-likelihood schemes.

**Figure 7:**
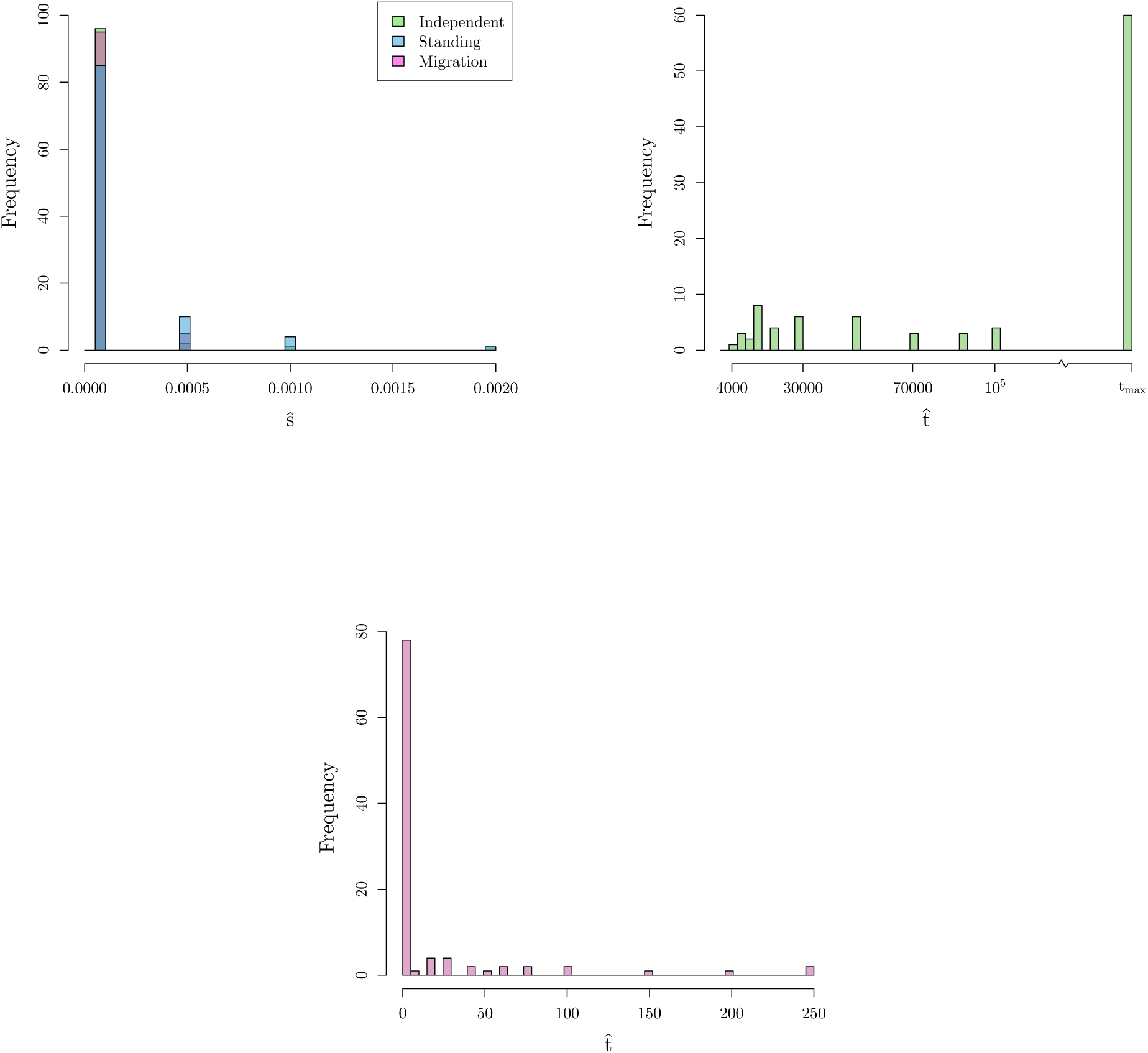
Histograms of MCLE for **parameters** estimated under **incorrect models**. (a) Histogram of MCLE of the **strength of selection** (*s*) **under all convergent models** where the **neutral model** is true model used for simulations. (b) Histogram of MCLE of the **standing time** (*t*) **un-neutral der standing variant model** where the **independent mutation model** is true model used for simulations (*s*= 0.01, *N*_*e*_ = 100,000). (c) Histogram of MCLE of the **standing time** (*t*) **under standing variant model** where the **migration model is true model** used for simulations (*m* = 0.001, *s* = 0.01, *N*_*e*_ = 10,000).

**Independent mutations model** As shown in Figure 6b, we are able to correctly distinguish between a neutral model of no selection and the true independent mutation model by at least 160 composite loglikelihood units even for relatively weak selection (*s* = 0.005). This difference increases as the true value of *s* increases. This same relationship is true when comparing the migration model to the true independent mutation model. Therefore, we have good ability to distinguish the independent sweeps model from neutral and migration model over a range of selection coefficients.

Our ability to distinguish between the standing variation model and the true independent mutation model is less clear. When the true *s* is small, the two models have comparable composite log-likelihoods, with differences ranging from −3 to 20. This difference decreases, with higher likelihood for the true independent mutation model more frequently, as *s* increases. This result makes sense when we look into the maximum likelihood estimate of the parameter *t* (Figure 7b). We obtain estimates of *t* approaching our highest value on the grid (10^6^). Thus, we may not be able to distinguish between the cases where the origins of the beneficial allele are truly independent or whether selection has been on a single variant that has been standing independently for a long time as these two models converge for large *t*.

**Standing variant model** Simulating under the standing variation model, the picture is more complicated. Like the other models, we can exclude the neutral model, although note that this would become challenging when the allele has been standing at high frequencies, *g*≫ 0 (Berg and Coop, 2015). When the independent standing time, *t*, is small, we see little difference in the composite log-likelihoods between the true standing model and the migration model. As *t* increases, we see a larger difference between these two models. However, as *t* increases, the composite log-likelihood difference between the independent mutation model and standing variation model tightens around 0. These results fit our expectations as we know the models look similar in the extreme values of *t*, the migration model when the standing time is small and independent mutation model when the standing time is large, respectively.

**Migration model** We are able to distinguish the migration model from the neutral and independent sweeps model. However, the standing variation and true migration model are again somewhat confounded. The values of the composite log-likelihood differences range from −44 to 123 when *m* = 10*^-4^* and this range narrows closer to 0 as *m* increases. These results fit our understanding when we again look at the MCLEs of *t* in the standing model. Now, the estimates are at *t* = 0 (Figure 7c) indicating it is hard to distinguish between convergence that is due to migration or selection on a shared standing variant that has only been standing for a very short time, as they result in similar patterns in decay of coancestries.

**Summary** We can clearly distinguish the outcomes of the migration and independent sweeps models from each other. Both models are hard to distinguish from the standing variation case, but in very different regimes of the standing variation model. The estimated time the variant has been standing (*t*) for is a helpful indicator of the mode of convergence, with very low estimates meaning that the standing model is indistinguishable from the migration model, while very high estimates mean that the standing model is indistinguishable from the independent sweeps model. When data is simulated under the standing model with intermediate values of *t*, we can distinguish this from both independent sweeps and recent migration models. This is because an intermediate value of *t* generates a covariance pattern not well explained by either other model. Therefore, while comparing the maximum composite likelihoods between models is useful, the estimated value of *t* is useful in judging the different models.

#### 3.2.3 Evaluating properties of the estimators and models for real datasets

Our use of a composite likelihood means that we cannot rely on standard asymptotic properties of likelihood estimators to construct confidence intervals or help with model choice (e.g. AIC). Therefore, we take a parametric-bootstrapping approach, simulating datasets under the MCLEs of various models matched for sample sizes and number of segregating sites and other qualities (recombination rate and size of the region, *N*_*e*_, neutral **F** matrix) as the original data. See Appendix A.3 for more details. From these simulations, we generate a distribution of composite-likelihood ratios. Specifically, we wish to understand if we have support for a model (*j*) as compared to a seemingly less likely model (*i*); this could be a model with selection to one without, or a model with standing variation compared to one with independent mutations. We simulate datasets under one model (*i*), using the MCLE of that model applied to the real data, we then estimate the maximum composite log-likelihood of dataset *k* under this model (*L*_*ki*_), and the maximum composite log-likelihood under a second model *j* (*L*_*kj*_) and form the distribution over our simulations of the difference *L*_*kj*_ − *L*_*ki*_. We can then compare the value of the composite log-likelihood ratio (*L*_*Dj*_ − *L*_*Di*_) obtained for our true dataset *D* to this distribution to obtain the parametric-bootstrap p-value for the comparison the alternative model (*j*) compared to the null model (*i*). Additionally, we generate parametric-bootstrap confidence interval for parameters of interest, particularly *t*, the minimum age of the standing variant, as this parameter is informative about the overlap of models as shown above.

## 4 Applications

### 4.1 Copper tolerance in *Mimulus guttatus*

The study of adaptation to toxic mine tailings is a classic case of rapid local adaptation to human altered environments (MacNair et al., 1993). We apply our inference method to investigate the basis of the convergent adaptation seen between populations of the annual wildflower *Mimulus guttatus* to copper contaminated soils near Copperopolis, CA. Wright et al. (2015) sequenced pooled samples from 20-31 individuals from two mine and two off-mine populations from two distinct copper mines in close geographic proximity (all populations within 15 km of each other) to 34-72X genome-wide coverage for each population. They observed elevated genome-wide estimates of genetic differentiation between mine and off-mine populations (*F*_*ST*_ M/OM= 0.07 and 0.14), with similar levels of differentiation between the mine populations (*F*_*ST*_ MM= 0.13). Only a small number of regions had high levels of differentiation. Here, we focus on the region with the strongest signature of differentiation between the two mine/off-mine pairs found on Scaffold8 by Wright et al. (2015). They observed low genetic diversity within each mine population in this region compared to off-mine populations. When the mine populations are compared to each other, they have elevated differentiation in this region, except for in the center where they share a nearly identical core haplotype. This pattern suggests the sweeps may not have been independent within each mine population, and that the sweep is possibly shared either due to migration or selection of shared standing variation.

We estimate the **F** matrix using SNPs from twelve scaffolds that showed no strong signals of selection (shown in Table S6). Using all SNPs in the 169.3 kb Scaffold8, we apply our inference framework to both identify the locus under selection and distinguish between modes of convergence between the two mine populations. We move the proposed selected site along this scaffold and calculate the composite likelihood under our three modes of convergent adaptation: (1) both mine populations have had independent mutations at the same locus, (2) the beneficial allele was standing in one of the mine populations and was spread via migration into the other mine population where it is still standing prior to the onset of selection (as detailed in Appendix A.4), and (3) the beneficial allele arose in one of the mine populations and spread to the other via migration. We estimate the maximum composite likelihood over a dense grid of parameters used to specify these models (Table S7). For the migration model, we allow both adapted populations to be possible sources. We use an *N*_*e*_ = 7.5×10^5^, calculated from the observed pairwise diversity *π* = 4*N*_*e*_*μ* using a mutation rate of *μ* = 1.5 × 10^-8^ and *r*_*BP*_ = 4.72×10^-8^ (Lee, 2009).

In Figure 8a, we summarize the results, showing the difference in maximum composite log-likelihoods between a given model of convergence and the neutral model of no selection as a function of the proposed selected sites along the scaffold. We see the three likelihoods peaking when the selected site is approximately at position 303-308 Kbp and that the model with the highest likelihood is selection on shared ancestral standing variation.

**Figure 8:**
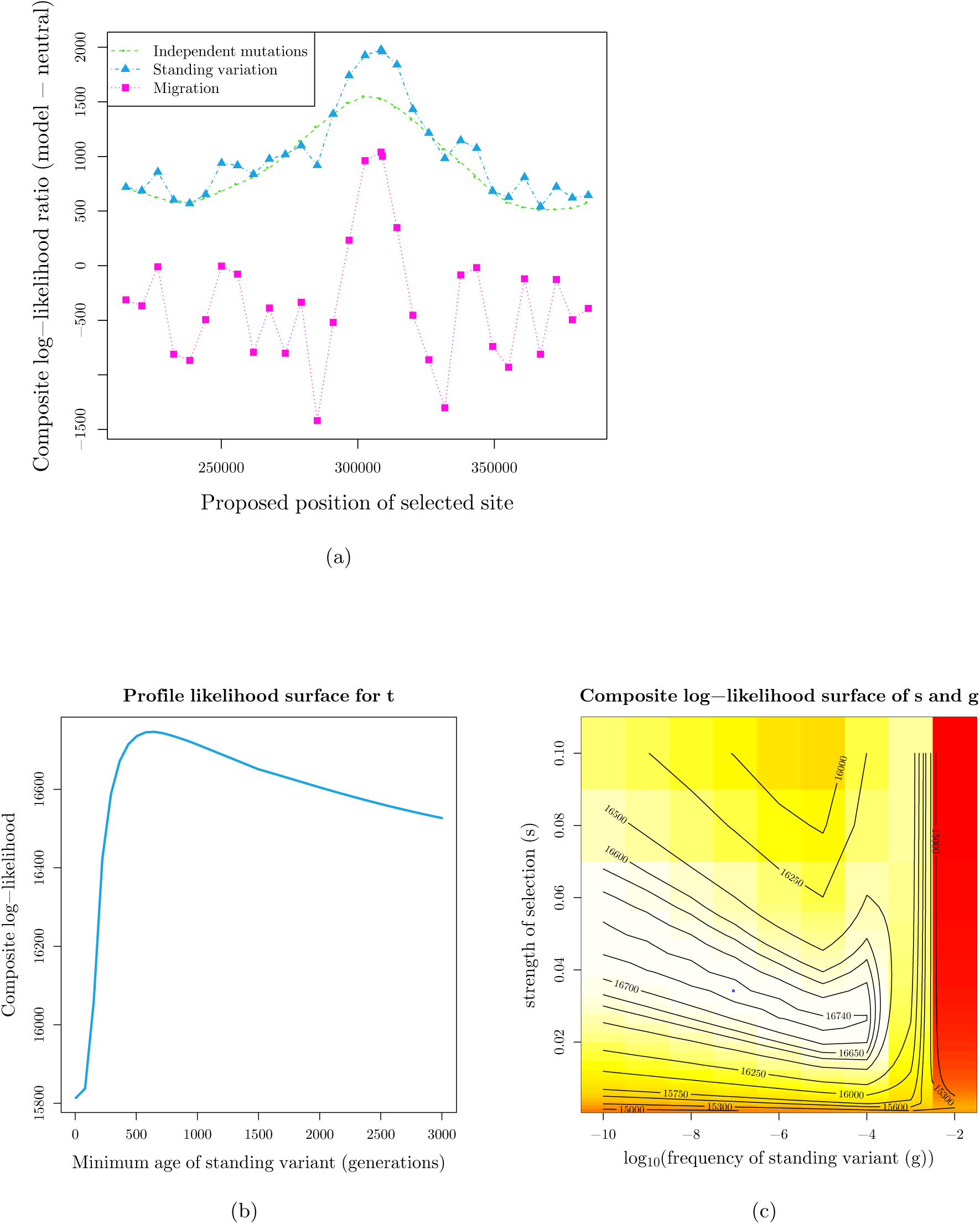
Inference results for *Mimulus guttatus* copper tolerance adaptation on Scaffold8. (a) Composite loglikelihood ratio of given model relative to neutral model of no selection as a function of the proposed selected site. We show likelihoods for the standingsource model maximizing over possible sources, but all results can be seen in Figure S7a.(b, c) MCLE of parameters in standing variation model with position 308503 as selected site. (b) Profile composite loglikelihood surface for minimum age of standing variant, maximizing over other parameters, with peak at 646 generations (c) Composite log-likelihood surface for strength of selection versus frequency of standing variant. Blue circles represents point estimate of joint MCLE (*ŝ* = 0.034, *ĝ* = 10*^-7^*). *t* is held constant at MCLE of 646 generations.

To judge the significance of differences in the composite log-likelihood between the standing-source model and the other models we used our parametric-bootstrap procedure. We simulated 100 datasets under the independent and migration modes of convergent adaptation at their MCLE as well as a neutral model with no selection (see Appendix A.3 for details). For each simulated dataset, we calculate the composite log-likelihood ratio comparing the standing source model to the likelihood of each of the other models (for their respective simulations), under the same parameter grid as the original data (Table S7) but holding the location of the selected site and, where relevant, the source population constant at their respective MCLEs used for simulation. Our observed composite log-likelihood ratio, comparing the standing source model to each of the others, was well outside the range those obtained by simulation (implying a parametric-bootstrap p-value of < 1/100). The smallest difference is under the migration model where the range of out 100 composite log-likelihood ratios is [4.12, 749.45], while the observed ratio is 945.95 (see Table S8 for all results). These results suggest that the non-standing source models offer a significantly worse fit to the data.

Focusing on the standing-source model at the most likely selected site, we can obtain parameter estimates for the strength of selection (*s*), standing frequency of the beneficial allele (*g*), and the amount of time that the beneficial allele has been standing in both mine populations after they have been isolated but prior to selection (*t*). The strength of selection and starting frequency of the allele are confounded (Figure 8c) as expected. Our maximum composite log-likelihood parameter estimates suggest selection was relatively strong (>0.02) and the allele was not standing at very high frequencies (< 10*^-4^*) when selection began. We see the maximum composite log-likelihood is obtained when the standing time (*t*) is approximately 646 generations (Figure 8b). As the Copperopolis *Mimulus* are annual, this corresponds to 646 years. We obtained 95% parametric-bootstrap confidence interval of [364, 9525] generations (years), by simulating under the standing-source at our MCLE (see Appendix A.3). This time also has the interpretation of the minimum age of the standing variant as it has been standing for at least this amount of time and potentially longer in the source population. As copper mining started in 1861 in this region (Aubury, 1902), this suggests the tolerance allele was present prior to the onset of mining again consistent with the variant being a standing variant when selection began.

There is little information about the source population of the standing variant (we obtain identical likelihood surfaces for either copper population as the source, see Figure S7a). This is perhaps unsurprising as there is relatively little hierarchical structure among the populations. Additionally, we tested the standing variant model with no source and saw no difference in the likelihood surfaces over the proposed selected sites (Figure S7a). The maximum composite-likelihood estimate of *t* is higher for the models of standing variation with a source than the simple model of standing variation (see Figure S7b). This is likely because making one of the populations a source of the standing variant increases the covariance around the selected site among the selected populations, as described in Appendix A.4, and so the model compensates by increasing the rate of decay of this covariance.

### 4.2 Industrial pollutant tolerance in *Fundulus heteroclitus*

We demonstrate how our method can be extended to more complex population scenarios. Populations of the Atlantic killifish, *Fundulus heteroclitus*, have repeatedly adapted to typically lethal levels of industrial pollutants (Nacci et al., 1999, 2010). Reid et al. (2016) have sequenced 43-50 individuals from four pairs of pollutant-tolerant and sensitive populations along the U.S. Atlantic coast (see Figure 9a), sequencing each individual to 0.6-7X depth. The southern pair of populations form a distinct clade relative to the northern populations, consistent with a phylogeographic break centered on New Jersey (Duvernell et al., 2008).

**Figure 9:**
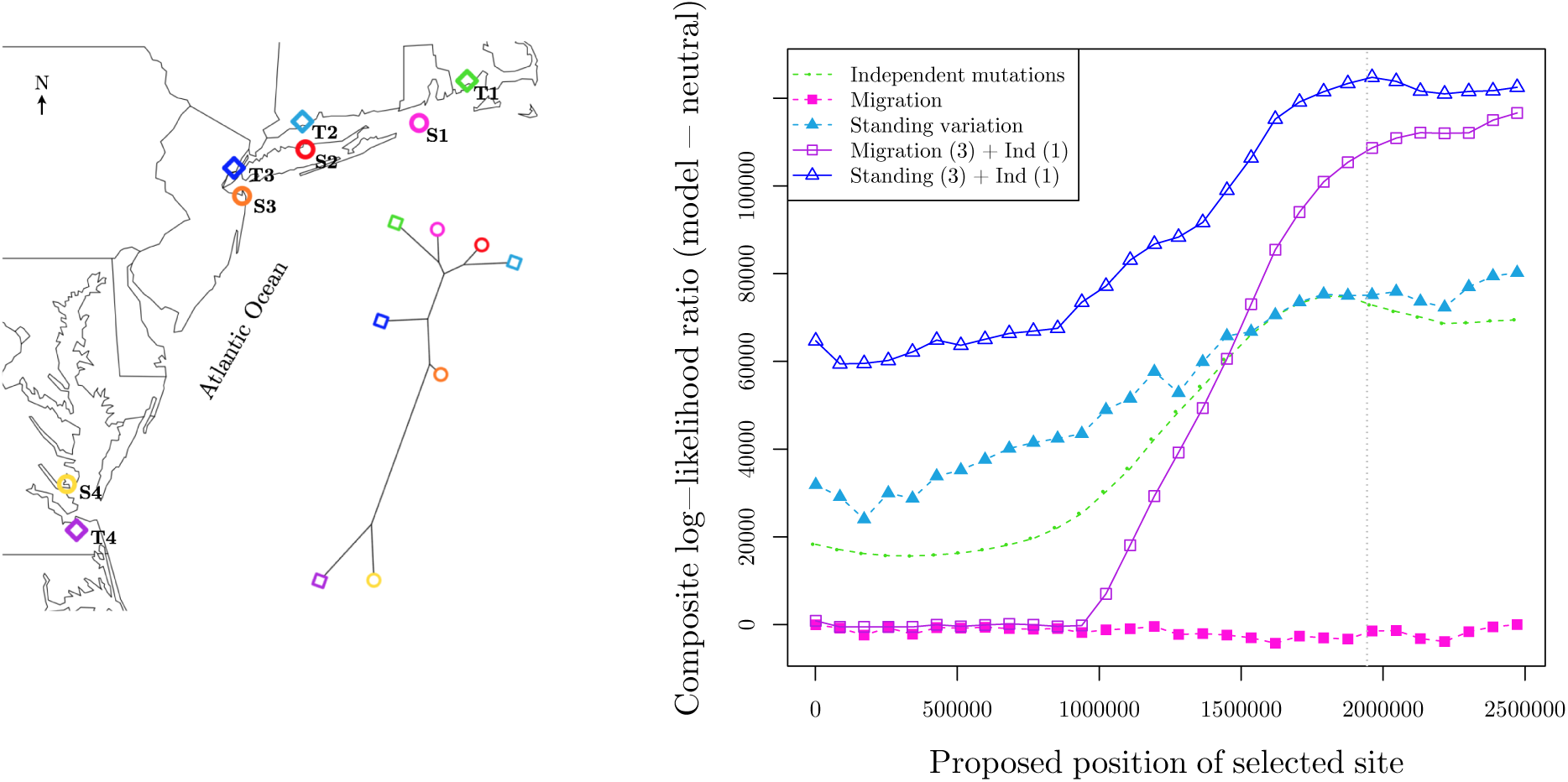
Inference results for *Fundulus heteroclitus* pollutant tolerance adaptation on Scaffold9893 (a)Map of sampled killifish populations with phylogenetic tree, showing that the southern pair (T4, S4) are more distant than other populations. Tree is estimated from genome-wide biallelic SNP frequencies using Phylogeny Infer-ence Package (PHYLIP) relative to neutral Gene Frequencies and Continuous Characters Maximum Likelihood (CONTML) module (see Reid et al. (2016) for more information). (b) Composite log-likelihood ratio of given model model of no selection as a function of the proposed selected site. Closed points represent models where all four populations have same convergent mode while open points represent Southern population (T4) having an independent mutation at the proposed selected site. We show likelihoods maximizing over possible sources, but all results can be seen in Figure S9. The AIP locus position is marked by the vertical,dashed gray lines.

Reid et al. (2016) found that a number of the strongest signals of recent selection are shared between all tolerant populations, suggesting genotypic convergent adaptation. We focus our method on their strongest signal of selection, Scaffold9893 (the scaffold containing the aryl hydrocarbon receptor interacting protein (AIP) gene), where all four pairs of tolerant/sensitive populations sampled show high levels of differentiation. Here, we test the hypotheses that all four tolerant populations show convergent adaptation due to our three previous modes of independent mutation, migration, or selection on shared ancestral variation. For our standing variation model, we specified the source of the standing variant (as described in Appendix A.4). We also test the hypotheses that there is an independent mutation in the southern tolerant population while the three northern populations are sharing a sweep at this locus, either due to migration between populations or selection on variation present in the ancestor of the Northern populations. This latter set of hypotheses is consistent with the fact that Reid et al. (2016) detect a shared haplotype in the three northern tolerant populations while a different haplotype appears to have swept in the southern tolerant population. We estimated the **F** matrix from four scaffolds that show no strong signal of selection, and it is shown in Table S9. We use *N*_*e*_ = 8.3 × 10^6^ and *r*_*BP*_ = 2.17 × 10^-8^ (N. Reid personal communication).

The results are summarized in 9b. For all models with migration or selection on standing variation, we plot the maximum composite log-likelihood for the most likely source at each location of the selected site (to reduce the number of lines plotted, see Figure S9 for the full figure). We see the model with the highest composite log-likelihood is when convergence is due to selection on shared standing variation in the North and an independent mutation in the southern tolerant population. This occurs when the selected site is at approximately position 1.96 Mbp on the scaffold.

To assess the significance in the composite log-likelihoods of this model and the other models tested, we simulate 100 datasets under each model at their MCLE (see Appendix A.3 for details). We calculate the composite log-likelihood ratio for each simulated dataset to compare the standing variation in the North with an independent mutation in the South model to the others models used for simulation. We calculate the composite likelihoods under the same parameter space as used for the original data (Table S10), holding the location of the selected site and the source population constant at their MCLEs used for simulation. For the neutral model and the three models where all four tolerant populations have the same mode of convergence, the observed composite log-likelihood ratio was far outside the range of values obtained from the simulations (see Table S11 for all results), suggesting these models offer a significantly worse fit to the data (parametric-bootstrap p-value < 1/100). However, this is not true for the model where migration is occurring in the three Northern selected populations while there is an independent mutation at the same locus in the Southern tolerant population. Here, the range of the difference in maximum composite loglikelihood for 100 simulations is [-24675, 38997], while the observed difference is 8121 (parametric-bootstrap p-value = 0.58; Figure S10). Thus we are unable to discern these models at their MCLEs.

Under the highest likelihood model of standing variation in the North and an independent mutation at the same locus in the South, we obtain the maximum composite log-likelihood estimate of the minimum age of the standing variant, *t*, of eight generations (Figure 10a). From simulating under this model at the MCLE, we obtain a 95% parametric-bootstrap confidence interval for *t* of [5, 310] generations. Thus under the standing-source model, the allele has only been standing for a very short time independently in the northern populations prior to selection. This is consistent with our observed overlap for the standing variant model and migration model. The confidence interval for *t* does not include 0, but that is also consistent with simulations under the migration model where inferred standing times are often slightly above zero (Figure 7c and Figure Figure S12). Together these results again suggest we are unable to differentiate between the models where the southernmost tolerant population has an independent mutation and the three northern populations are sharing the beneficial allele, either via migration or selection on the same young standing variant.

**Figure 10:**
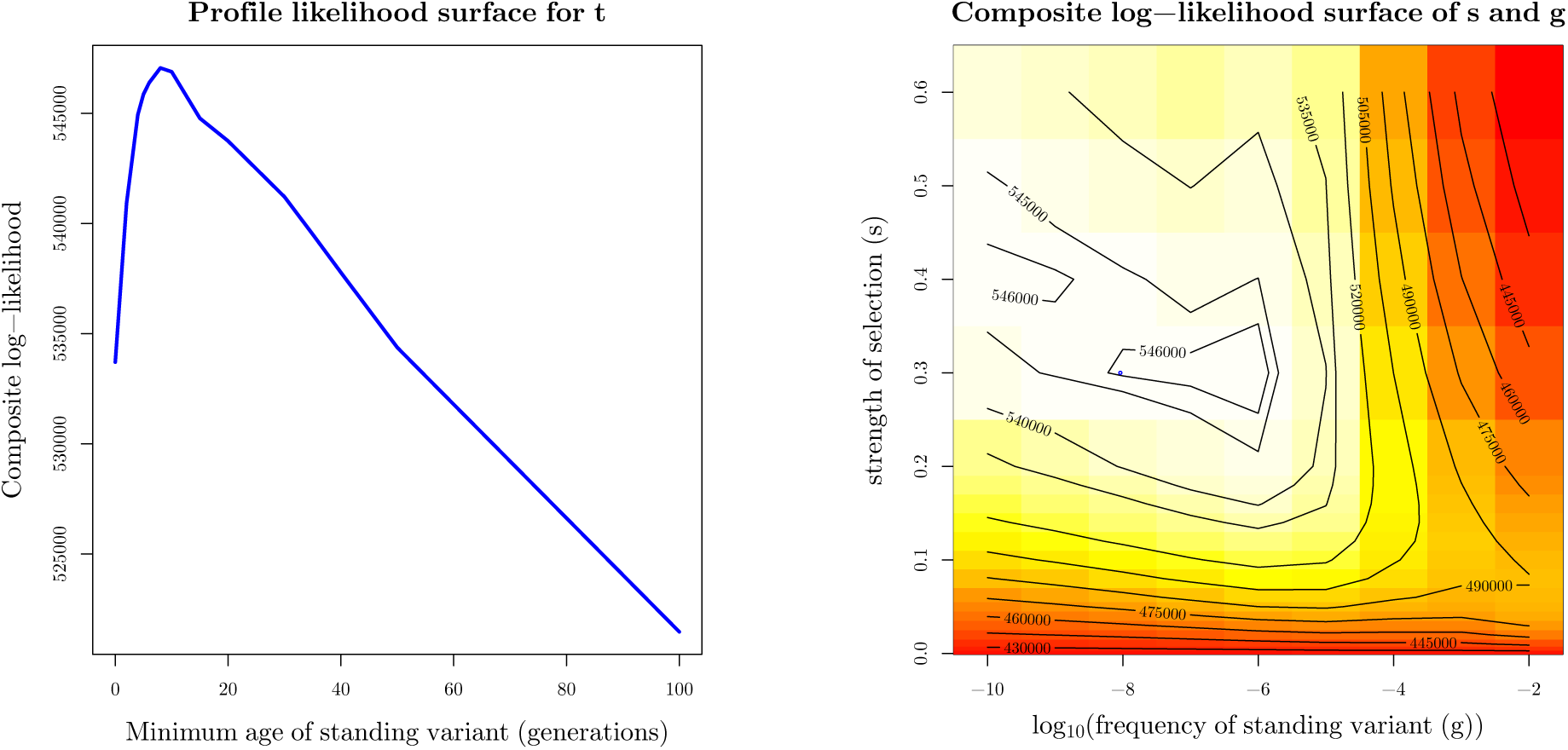
The composite log-likelihood surfaces for the parameters for *Fundulus heteroclitus* convergent data in combined standing variation and independent sweep model with position 1961198 on Scaffold9893 as selected site and population T3 as source. (a) Profile composite log-likelihood surface for minimum age of standing variant, maximizing over other parameters, showing the beneficial allele has been standing for a very short amount of time in our three northern populations (8 generations). (b) Composite log-likelihood surface for strength of selection versus frequency of standing variant. Blue circle represents point estimate of joint MCLE (ŝ = 0.3, ĝ = 10^-8^). t is held at MCLE of 8 generations.

We see partial confounding of the strength of selection and the frequency of the standing variant (Figure 10b) but our results indicate selection has been very strong (>0.3) and the allele was initially at a very low frequency (< 10*^-6^*). For the migration in the North model, we obtain similar MCLE of *s* of 0.4. Lastly, both the standing variation or migration in the North models has the highest composite log-likelihood when the source population of the standing variant is T3, the southernmost population sampled in the North (standing variation composite log-likelihood = 547060, migration composite log-likelihood = 537744), but this model may not be distinguishable from that where the source is T2 (standing variation composite log-likelihood = 545580, migration composite log-likelihood = 533426).

## 5 Discussion

In this paper we have presented a novel approach to identify the loci involved in convergent adaptation and to distinguish among the three ways genotypic convergence can arise: selection on (1) independent mutations, (2) a variant standing independently in the selected populations, and (3) beneficial alleles introduced via migration. We leverage the effects selection has on linked neutral sites via a coalescent-based model approach that captures many of the heuristics that have been used in previous studies. This approach also allow us to potentially distinguish between more subtle models, such as the origin and the direction of gene flow of a beneficial allele, since they are explicitly modeled in our framework. Our approach takes advantage of information among all of the population samples simultaneously while accounting for population structure. Therefore, it naturally accommodates information from across multiple samples, rather than just pairs of populations, and thus offers a number of advantages in identifying the mode of convergence over other approaches. We provide the relevant R code for our approach in https://github.com/kristinmlee/dmc.

**Distinguishing among models** We have demonstrated that our method is able to accurately distinguish among modes of convergent adaptation, across a relatively wide parameter space, in simulated data. However, we do see some confounding of models in particular regions of parameter space. In particular, we see the patterns generated from a model of selection on ancestral standing variation can look like our expectations for the other two modes of convergent adaptation for extreme values of the parameter *t*, the time the beneficial allele has been standing time independent in the selected populations.

When *t* is small, we see confounding between the standing model and a model of convergence due to gene flow. The two models are very similar since in our standing variation model, as *t*→0, the covariance in the deviations of a neutral allele between selected populations approaches the variance within a selected population. The strong overlap in models is especially true when we have a source for the standing variant. Intuitively, this indicates that the beneficial allele is on a haplotype that is mostly shared among the selected populations. This can be due to a very young standing variant shared amongst very closely-related populations from an ancestral population, a standing variant that was shared by gene flow before selection, or by the selected haplotype quickly moving across populations by gene flow after selection began (which are all closely related models, see Welch and Jiggins, 2014, for additional discussion).

To illustrate distinguishing between these possibilities we now briefly revisit our applications. The Northern tolerant killifish populations, under a standing variation model with gene flow prior to selection, have a very low estimate of the standing time *t* (8 generations with 95% CI [5, 310] generations). However, given this very low estimate of *t*, the allele cannot have been standing since the common ancestral population of T1, T2, T3 (which we estimate to coalesce more than 800 thousand generations ago, assuming no migration, using the estimation procedure outline in Appendix A.3.1). Therefore, the allele must be shared by gene flow among the three populations and it seems likely that the migration of the allele occurred either after selection began in one of the populations or very shortly before, with our parametric-bootstrapping approach suggesting we are not able to discern these two models. Interestingly, Reid et al. find no clear signals of admixture from migration elsewhere in the genome between Northern tolerant populations, suggesting that the migration of this allele might be a rare event, although we note that this may reflect a lack of power to detect gene flow.

The case for adaptation from ancestral standing variation is more clear for the *Mimulus* copper tolerance example. Here, the estimate of *t* is much greater than zero (646 generations with 95% CI [364, 9525] generations) and indeed older than the putative selection pressure (approximately 150 generations ago). Additionally, the standing variant model considerably outperforms the other models and the results of our parametric-bootstrapping approach support this. In this case, we again favor the model that incorporates gene flow prior to selection on standing variation. The level of neutral differentiation of the mine populations very likely reflects much more than 646 generations of drift (see Appendix A.3.1), thus it seems likely that this allele is shared between the mine populations by gene flow but that the allele was standing in both populations for some time before selection began. Together these applications show distinguishing among models of convergence is possible in some cases, but may require extra knowledge of population history to aid our inference and understanding.

Conversely, when *t* is large, we see a collapse of our standing model onto a model of convergence due to independent mutations in our selected populations. This intuition holds forwards in time since as *t→ ∞* generations, recombination in our isolated populations independently breaks down the similarity of the haplotypes carrying the beneficial mutation. Thus, when selection for the standing variant begins, even tightly-linked, hitchhiking neutral alleles will not be shared between populations more than expected by chance. This is also the case when beneficial alleles arise multiple times independently. For example, in the case of the killifish, it is formally possible that the signal of independent selection in the Southern tolerant population is actually due to a very old standing variant shared with the Northern populations where there is almost no overlap between the Southern and Northern tolerant populations in the haplotype the selected allele is present on, even close to the selected site. As the precise functional variant(s) in this swept region are currently unknown (Reid et al., 2016) it is hard to totally rule out this very old standing variant hypothesis. In other cases it may be possible to rule out the standing variant hypothesis with very large parameter estimates of *t* if we know more about the population histories (i.e. our selected populations split more recently than the standing time). Additionally, it may be possible to totally rule out the standing variant hypothesis in cases where if the functional variants can be tracked down to clearly independent genetic changes (e.g. Tishkoff et al., 2007). However that degree of certainty may be difficult to achieve in many cases.

**Extendibility and flexibility of our approach** We show the applicability of our method on two empirical examples of convergent adaptation: the evolution of copper tolerance in *Mimulus guttatus* and of pollutant tolerance in *Fundulus heteroclitus*. The latter exemplifies the extendibility and flexibility of our approach. As the number of selected populations increase, our potential number of hypotheses grows since any grouping of two or more populations could share selection due to migration or standing variation. Additionally, with more populations, we have more potential sources of the beneficial allele in the migration model. Our model could also be extended to have selection occurring in some of the adapted populations and the neutral model in others, to identify genomic regions that are not experiencing convergent adaptation among all populations sharing the selected environment. These models are all relatively easy to implement into our framework; however, the sheer number of possible hypotheses as the number of populations grows will likely call for some more systematic way of implementing these models and exploring their relationships.

**Caveats and possible extensions** Studying repeated evolution has long played a key role in evolutionary biology as a tool to help identify the ecological and molecular basis of adaptation. It is worth noting with this approach, we are able to identify sweeps in the same region and whether they appear to be shared or independent. However, in the scale of an entire genome, it may be possible for two, functionally unrelated sweeps to overlap. In the case of adaptation via independent mutations across multiple populations, it is especially hard to determine whether selection at the same site was acting on the same phenotype. It is potentially more plausible to claim that the phenotype and selection pressure are shared among populations in cases where the swept haplotype is shared. Ultimately, in demonstrating convergence, we will have to rely on a range of evidence. Shared sweeps can offer one substantial piece of evidence, particularly when we are studying recent adaptation to a strong selective pressure that is distinct to the adapted populations.

In addition to assuming that the same locus is under selection in all adapted populations, we assume a single selected change underlies the sweep within a population and that recombination is free to break down associations between neutral alleles and this selected variant. If, for instance, selection acts on an epistatic, haplotypic combination of allele that sweeps, a long haplotype could be shared between populations not due to recent migration but because selection acts against recombinants breaking up the haplotype (Kelly and Wade, 2000). Convergent adaptations due to shared inversions also violate the assumptions of our method. Inversions can repress recombination across the entire inversion (see Kirkpatrick, 2010, for a recent review). Inversions significantly alter both neutral and selective model expectations (e.g. Guerrero et al., 2012) and could lead to long shared haplotypes among populations even if the shared inversion is old. It may be possible to use our approach to model the decay in coancestries outside of the inverted region, but this requires knowledge of the inversion and its break points *a priori* and a detailed knowledge of recombination rates surrounding the inversion.

Throughout this paper we assume that the sweeps have fixed recently, and it will be important to relax this assumption. In these cases, models of migration that include selection against maladaptive migrants (Barton and Bengtsson, 1986; Charlesworth et al., 1997; Roesti et al., 2014) will be important to consider. Long-term selection against migrant alleles (i.e. due to local adaptation) lowers the effective migration rate at linked neutral sites and so will distort the covariance relationships among populations (and may in some cases confound the signal of the mode of convergence). These deviations could be incorporated into our models, allowing us to perform inference under these models. However, in practice we would likely be underpowered, as we only model segregating sites we cannot (in the current framework) fully account for selection that deepens the absolute divergence among particular populations.

Additionally, our framework could be extended in various ways to both leverage more information and model more biologically relevant or interesting scenarios. There is more information to be gained from haplotypes and associations between sites that we fail to include in our composite likelihood when we sum across information from individual sites. Here we use this approach to analyze genomic regions that we *a priori* assume to be under convergent selection. In part this is due to the phylogenetic relationships among the populations (with convergent populations not being sister to each other). Additionally, we could then model ancestral sweeps to address whether sister populations sharing an adapted phenotype is truly convergent or simply due to selection in their ancestor Racimo (2016). We are currently working on ways to efficiently extend this approach to the application of genome-wide data to scan for genomic regions exhibiting convergence.

## 6 Acknowledgements

We wish to thank members of the Coop lab for helpful discussion and feedback on earlier drafts. We’d also like to gratefully acknowledge Noah Reid, Andrew Whitehead, John Willis, and Kevin Wright sharing their data and thoughtful comments. We thank Nicolas Bierne, Joachim Hermisson, and an anonymous reviewer for valuable suggestions on an earlier draft. This work was supported by the National Science Foundation Graduate Research Fellowship awarded to K. Lee (1148897) and by grants from the National Science Foundation under Grant No. 1353380 to John Willis and G. Coop and the National Institute of General Medical Sciences of the National Institutes of Health under award numbers NIH R01 GM108779 awarded to G. Coop.

## A Appendix

### A.1 Coalescent interpretation of covariances and F-matrix estimation

Let *x*_*il*_ be the allele frequency of allele 1 in population *i* at locus *l*, and that the frequency of this allele in the ancestral population is ϵ_*l*_. Consider the covariance Cov(Δ*x*_*il*_, Δ*x*_*jl*_) over replicates of the drift processes at locus *l*. We can write

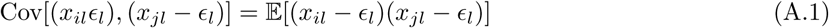

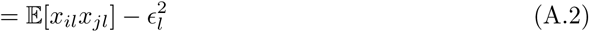

which follows from the fact that 𝔼[*x*_*il*_] = 𝔼[*x*_*jl*_] = ϵ_*l*_. We can interpret 𝔼[*x*_*il*_*x*_*jl*_] as the probability that we sample a single allele in *i* and an allele in *j* and that they both are of type 1. Taking that interpretation, assuming that there is no mutation, 𝔼[*x*_*il*_*x*_*jl*_] is the probability that, tracing back a coalescent lineage from *i* and a lineage from *j*, both lineages trace back to type 1 alleles in the ancestral population. Let our pair of lineages drawn from *i* and *j* coalesce with probability *f*_*ij*_. If our lineages coalesce before reaching the ancestral population then they will be identical by descent, and share the ancestral choice of allele. Therefore, we can write

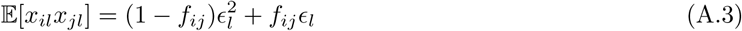

Then we can rewrite the covariance

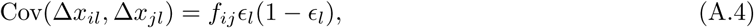

and for the variance we set *i* = *j*. Thus, under a model of genetic drift alone, we can interpret the entries of our covariance matrix as expressions of the underlying coalescent probabilities.

**Estimating F** In the main text we assume that we have estimates of our neutral coancestry matrix **F**. We now describe how we obtain these. From above, Equation A.3, the expectation of *x*_*il*_*x*_*jl*_ across loci is

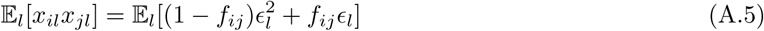

Therefore we can write estimate *f*_*ij*_ as

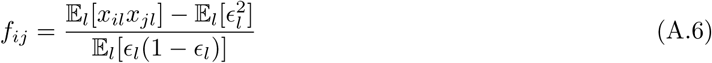

We can obtain an unbiased estimate of 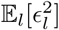 and 𝔼_*l*_[ϵ_*l*_(1- ϵ_*l*_)] using the sample allele frequencies from two populations on either side of the root of the population phylogeny (see Supplement of Lipson et al., 2013). Let *i′* and *j′* be a pair of populations that span the root of the population tree, then we can use the estimate

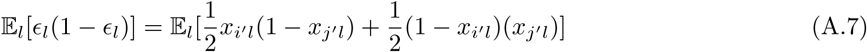

Likewise, we use the estimate

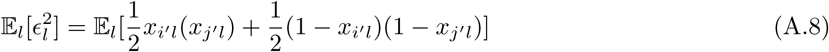

An estimate of the term 𝔼_*l*_[*x*_*il*_*x*_*jl*_] can be obtained by using the sample frequency of allele 1 in populations *i* and *j*. However, as we only have a sample from the population frequency we need to account for the finite sampling bias within populations (*i* = *j*). Let *n* be the sample size in population *i*, then

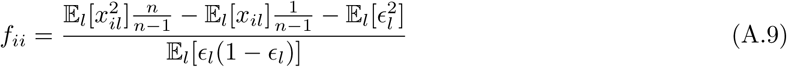

where our *x* are now sample frequencies. There is no finite-sample size correction for *f*_*ij*_, *i* ≠ *j* and Equation A.6 can be used directly.

In our simulations to show the effect of selection on the coancestry coefficients (Figure 3), we estimate *f*_*ij*_ in bins of fixed genetic size moving away from the selected site. We do this by approximating the expectations in the numerator and denominators in Equations A.6 and A.9 by the average of the expression over all of the SNPs that fall in a given genetic distance bin over all of the relevant simulations. To account for biases induced by defining the allele of interest, we randomize the reference allele at each SNP.

### A.2 Simulation implementation details

We perform coalescent simulations using mssel, a modified version of ms (Hudson, 2002) that allows for the incorporation of selection at single site (the code for this is provided in https://github.com/kristinmlee/dmc). The program allows the user to specify the frequency trajectory of the selected allele through time across populations, this trajectory is then used to simulate genetic data under the coalescent model conditioning on this trajectory (using the sub-divided coalescent model Hudson and Kaplan (1988); Kaplan et al. (1991)). We generate stochastic trajectories for the selected allele across populations and describe the simulation process below. We simulate multiple instances of the stochastic trajectories and average our results across datasets generated for these trajectories. We focus on a set of four populations with relationships as shown in Figure 1. Populations 2 and 3 are adapted to a shared novel selection pressure and populations 1 and 4 are in the ancestral environment.

The original implementation of mssel assumes only a single origin of the selected allele, which occurs moving backward in time when the frequency of the derived allele goes to zero in the final population it segregates in. We modified the mssel source code directly to accommodate multiple origins of the selected allele as is necessary in the independent sweep model. We do so by allowing an independent origin of the selected allele in any population where the frequency of the derived selected allele goes to zero, if that population currently has a migration rate of zero to any other population containing the selected allele.

#### A.2.1 Generating stochastic trajectories for the selected allele

We generate stochastic trajectories for the selected allele to be used as input for mssel to generate sequence data for given convergent adaptation scenarios. We simulate the allele frequency trajectory for the selected allele forward in time using a normal deviate approximation to the simulation the Wright-Fisher diffusion. Specifically, given the frequency of the beneficial allele at time *t*, *X*(*t*), we simulate its frequency at time *t* + Δ*t* according to

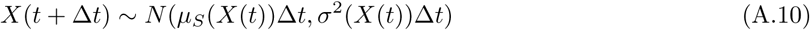

where *μ_S_* ( ) and *σ*^2^( ) are the infinitesimal mean and variance of the Wright-Fisher diffusion. We set Δ*t* = 1/(2*N*), representing one Wright-Fisher generation on the diffusion time-scale (2*N* generations). We set *X*(0) = *g*, the initial frequency of the beneficial allele. When selection starts from a new mutation, *g* = 1/(2*N*).

For all our models, the infinitesimal variance is

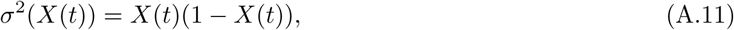

representing the effect of genetic drift.

For populations not impacted by migration, we condition our trajectory on the beneficial allele going to fixation forward in time. To do this we use the conditional infinitesimal mean

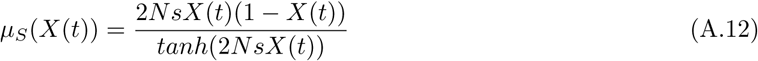

(see Przeworski et al., 2005; Berg and Coop, 2015, for previous applications). We simulate this process forward in time till fixation is reached. Given that we are assuming the sweeps completely recently, we have fixation occur at time zero so that the time of a new mutation is determined by the time of the sweep.

**Migration model** In the case of our migration model, there is one way migration from population *i* into *j*. The trajectory of *x*_*i*_ is simulated first forwards in time, conditioning on fixation, using the above approach. We then simulate the frequency in population *j* starting from *X*_*j*_(0) = 0, with the infinitesimal mean

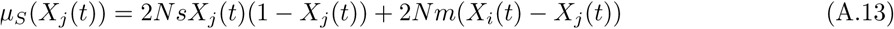

(expanded from Ewens, 2004). We simulate the process forward in time until the selected allele reaches fixation in both populations. The first population to reach fixation is held at frequency 1 until the other population fixes for the beneficial allele.

**Standing variation model.** We define the standing variation trajectory as having three phases, the neutral phase, the standing phase, and the selected phase. To specify a trajectory in which the beneficial allele has been standing at frequency *g* for time *t*, we simply hold the allele frequency constant for this amount of time. We simulate a stochastic neutral trajectory of our beneficial allele from frequency *g* to 0 backwards in time according to

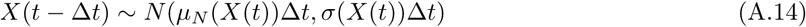

using the infinitesimal mean conditional of the neutral allele going to loss

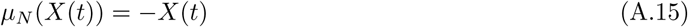

(see Przeworski et al., 2005; Berg and Coop, 2015, for previous applications). We simulate the selection phase forward in time for 2 log(1/*g*)/*s* generations. If the beneficial allele has reached fixation before this time, it is held constant at frequency 1 for the remaining time. If not, the trajectory is simply stopped at this time. This allows for the interpretation of the standing time and the time of the onset of selection to be the same throughout simulations. For the whole trajectory of a beneficial allele, we paste together these three components: neutral increase of allele from frequency 0 to *g*, the standing phase at frequency *g* for time *t* generations, and the selective phase. For populations not experiencing selection, the beneficial allele is kept at frequency *g* for the entire length of the trajectory. We acknowledge this is an untested approximation but think it has little impact on our results. The frequency of the standing variant matters mostly for estimating the duration of the sweep within populations, so its frequency during this standing phase is not as important as the frequency at the onset of selection. Additionally, we assume that *g* is small such that the probability of recombining off onto the other background during this phase is simply *r*. The frequency of the variant during the standing phase does impact the probability of coalescing before recombination (or vice versa) during this phase, but only weakly.

#### A.2.2. Details of coalescent simulations

In this section we give the details of the coalescent simulations. The mssel command lines can be found in Supplement S3. The mssel input can be interpreted as follows,

~~~
./mssel nsam_tot nreps nsam_anc nsam_der trajFile locSelSite -t *Θ* -r *ρ* nsites -I npops nAnc_pop1 nDerv_pop1 … nAnc_popi nDerv_popi
~~~

For all of the simulations we generate neutral allele frequency data for 10 samples from each of 4 populations. The populations are related to each other as shown in Figure 1. Note, we did 1000 replications of the simulations for parameters used to generate comparisons of average simulations coancestry coefficients compared to theoretical expectations. 100 replications were done for simulations used for parameter estimates and model comparisons. For simulations used for both, the first 100 runs were used.

**Independent sweep model.** We generated beneficial allele frequency trajectories under four different selection coefficients: *s* = [0.005, 0.01, 0.05, 0.1] under the independent sweep model with *N*_*e*_ = 100, 000. We set *r*, the per generation probability of cross-over between ends of the simulated locus, to 0.005. The neutral mutation rate, *μ*, for the entire locus is the same as *r*. We also simulate, with ms the same population structure with no selection to generate data to estimate the neutral coancestry matrix, **F**.

**Standing variation model.** With *s* = 0.01 and *g* = 0.001, we generated beneficial allele frequency trajectories for standing times *t* = [50, 250, 500, 1000, 5000] generations under the standing variation model with *N*_*e*_ = 10, 000. Our *t* references the time that the populations have been independent. Therefore, we adjusted the split times to ensure that the *t* of interest corresponded to the duration of time that the selected populations had the standing variant prior the populations joining in the ancestral population. The population split times were determined to ensure selection started after the populations were completely isolated and to maintain a similar ratio of time for 4 independent populations to 2 ancestral populations. We again set *r* = *μ* = 0.005. Again, neutral regions were simulated in ms using the same population structure (i.e. each parameter set had its own neutral data generated).

**Migration model.** Lastly, we simulated under the migration model with *m* = [0.0001, 0.001, 0.01, 0.1], holding *s* = 0.01 for *N*_*e*_ = 10, 000. Again, we simulated 10 samples from 4 populations related to each other as specified in Figure 1. Now, in mssel, we specify migration to start just prior to origin of the beneficial allele in the source population and to continue until the sweep has reached fixation (time zero in the past since we fix sweeps to complete at the end). We set population 2 to be the source and have 4*N*_*e*_*m* migrants from population 2 into population 3 each generation. We again set *r* = *μ* = 0.005. Neutral regions were again simulated using ms. Each set of parameters has its own neutral data generated as the migration rate impacts neutral coancestry as well.

#### A.2.3 Interpretating mssel output

The output from mssel and ms is in the form of haplotypes for each of the sampled chromosomes at polymorphic sites in addition to their positions on a scale of (0, 1). We use this to calculate sample allele frequencies at each site for each population. Prior to performing further estimations or analyses with these neutral allele frequencies, we randomize the reference allele so that there is no bias resulting from which allele was called ancestral or derived. We exclude sites where the average allele frequencies across populations are less than 5% or greater than 95%.

#### A.2.4 Composite likelihoods of simulated data under all models details

We calculated the composite log-likelihoods of each the simulated datasets under all models, including the neutral model, with the same parameter space shown in Table S1.

#### A.2.5 Maximum likelihood estimate of parameters from simulated data under correct model

We also calculated the composite log-likelihoods of each the simulated datasets under the correct model used to generate the data now with a more dense grid of parameters to obtain better estimates of the MCLE of each parameter. We allowed *g* to vary in the calculations of the MCLEs under the standing variation model. See Table S2, Table S4, Table S5.

#### A.2.6 Inference details: mean-centering allele frequencies and covariances, sample size correction, and speed-ups

Given that we do not know the true ancestral mean at locus *l*, ϵ_*l*_, we use the mean of the present-day sample allele frequencies at this locus, 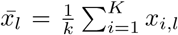. When mean-centering, we lose a degree of freedom so in calculating the likelihood it is necessary to drop information from one population. Since the information from the dropped population is incorporated in the mean, the choice of the dropped population is arbitrary. In matrix form, the mean-centered allele frequencies with one dropped population can be expressed as

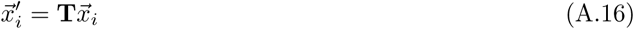

where **T** is an *K* − 1 by *K* matrix with 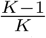on the main diagonal and 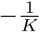 elsewhere. Prior to mean-centering, we randomize the reference allele at each SNP to account for biases induced by defining the allele of interest.

Now, we model the mean-centered allele frequencies as multivariate normal around mean zero with covariance proportional to a mean-centered parameterized covariance matrix (**F**^(*S*)')^ as

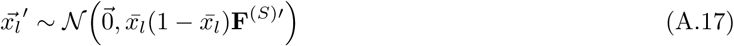

where we use the average present day allele frequency across populations at the locus, 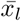, as an estimate of ϵ _*l*_ in the site-specific term in the covariance. We note that 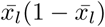 is a slightly downwardly biased estimate of ϵ (1 -ϵ), but for our purposes it seems sufficient to include this term as a locus-specific adjustment to the expected covariance.

To obtain the corresponding mean-centered covariance matrix, dropping the same population, we can apply the following matrix operations,

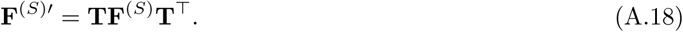

this new matrix is *K*-1 by *K*-1 and full rank.

Before mean-centering, **F**^(*S*)^, we apply a sample size correction to correct for the finite sampling bias. We add 1/*n*_*i*_ to the diagonal where *n*_*i*_ is the sample size in population *i*. We take twice the number of diploid individuals sampled in population *i* as *n*_*i*_ for data applications. In simulations, we use the number of chromosomes sampled in population *i* as *n*_*i*_. Note that both this mean-centering and sample size correction is also preformed on the neutral matrix, **F** before likelihood calculations under a neutral model with no selection.

To decrease some of the computational time involved in our likelihood calculations, we precompute the mean-centered covariance matrices with selection, **F**^(S)′^, for given bins of distance away from a putative selected site. We first divide our distances in our window into 1000 bins and take the midpoint of the distances in these bins to calculate **F**^(S)′^ as this matrix is a function of distance. To avoid the costly step of recomputing the corresponding inverses and determinants needed for likelihood calculations, we do this step first and use these values for all SNPs in a given bin, and store them and reuse them over all locations of the selected site.

Thus, we calculate the likelihood of mean-centered allele frequencies, 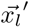, given our model *M* and its parameters Θ_*M*_, a given locus *l* as

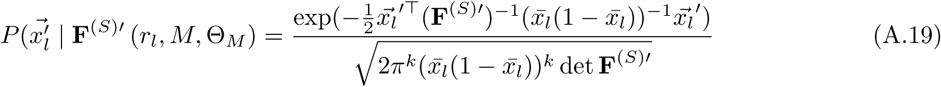

where *k* = *K* − 1, the rank of matrix **F**^(S)′^.

### A.3 Parametric bootstrapping approach details

To carry out the parametric-bootstrapping approach, we again perform coalescent simulations using mssel for simulations with selection and ms for neutral simulations. We specify the number of populations and the sample size for each populations (twice the number of individuals sampled). Now, instead of specifying *Θ*, we specify the number of segregating sites as the number of SNPs in our window of interest. We also simulate with the same population-scaled recombination rate and number of sites between which recombination can occur as the number of base pairs in our analysis window. To match the population-scaled recombination rate, we take the genetic map of our region *r* and scale it to be 4*N*_*e*_*r*, assuming that recombination is uniformly distributed over our region.We down-scaled the effective population size for computational efficiency in the generation of the simulations, which impacts both *ρ* and the times in the trajectories of the beneficial allele by a linear rescaling. Additionally, we specify the location of the selected site (*𝓁*) to be at the MCLE of the model used for simulation.

While in the rest of the paper we make use of stochastic trajectories, for the parametric-bootstrap simulations we generated deterministic trajectories of the selected allele to be used as input for mssel. This is because we need to set our simulations up to accommodate both the MCLE selection coefficient and the coalescent times within and between populations, which is somewhat fiddly to automate with fully stochastic trajectories across all the models. Now, we fix the time of the sweep to be

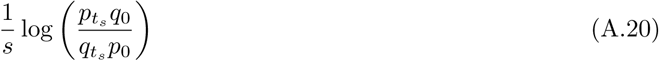

where *p*_0_, the frequency of the beneficial allele at time 0, is 1/2*N* for a new mutation or *g* for the standing variant model. While *p*_*t*_*s*, the frequency of the beneficial allele at fixation, is set to 0.999. For the migration model, we start this trajectory (from 1/2*N*) after the delay time (Equation 10) for recipient population(s). We simulate with migration after *δ* for a few generations. For the standing variant model with a source population, we start the selected allele trajectory (from frequency *g*) in the recipient population(s) after *t* generations. We simulate with a brief burst of migration at time *t* until the frequency of the beneficial allele goes to 0 in the recipient population(s), at a very low rate. This forces an instantaneous coalescent event back into our source population. The parameters (*s*, *t*, *g*, *m*, and the source population) are all set to the MCLE of the corresponding model.

We simulate each convergent and neutral model 100 times and interpret the output and calculate the likelihood of our simulated data (as detailed in Appendix A.2) under the model used for simulations and the model with the largest composite likelihood for the original data. The mssel command lines can be found in Supplement S4.

#### A.3.1 Approximating demography given a neutral F matrix

For the parametric bootstrap we need to simulate under a model of population structure that approximately matches that in our data. To do so we assume that our sampled populations are related through a bifurcating population phylogeny (with no neutral migration). While this is a crude approximation it allows us a good match to the observed *F* matrix of the data. and considerably simplifies the task of setting up the simulations. In practice since our method works with these covariances, and inferring the details of population structure is not our primary concern here, we view this as an acceptable compromise.

For simulating under the approximate population structure in our data, we need to estimate join times for population pairs. We use

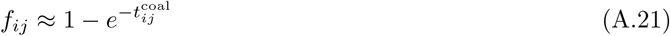

where 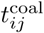 is in coalescent time units to approximate the shared branch length between populations *i* and *j*, assuming no migration. Migration will impact the coancestry coefficients and thus our interpretations of the coalescent times. For example, migration between two populations will increase their relatedness and can make their shared branch length appear longer. We also use this approximation to compare the split time between populations to the standing time for our adaptive alleles *t*, to judge whether they could have been standing for a given time between two populations, or if migration must be invoked.

To generate join times, we first solve for all 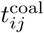 using A.21 from an estimated neutral **F** matrix. We find populations *i* and *j* with the largest 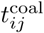. We approximate the join time as the average of the differences between the total time associated with each population (i.e. 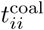 and 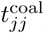) and the time between them 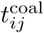. This follows from assuming drift is acting additively such that *f*_*ii*_*≈ f_ij_* +*f*_*i*_ where *f*_*i*_ is the coancestry coefficient associate with population *i* in isolation (see Supplement S2 for more). We then effectively join these two populations, updating all 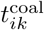 and 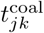 where *k* is any unjoined population to be the average of *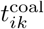* where *k* and 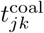 where *k*. We repeat this procedure, joining the two remaining populations with the largest 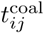 until all populations are joined. From this, we are able to specify join times for simulations that capture the general population structure of a given **F** matrix.

The population structure used for simulation is now represented in a bifurcating tree, which may fail to capture of the complexity represented in a given **F** matrix. Thus, when performing the composite-likelihood calculations we use a modified **F** matrix estimated using the procedure detailed in A.1 with neutral data simulated with these join times, to parameterize our models.

Additionally, these estimates for the between population coalescent times, assuming no migration and a bifurcating tree, can give us insight it is possible for the beneficial allele to have been standing for a given *t* since the ancestral population or whether it is necessary to invoke the model where migration has a role in spreading the beneficial allele prior it standing. For example, in our *Mimulus* analysis, we estimate our join time to be 0.050 in coalescent units. Our MCLE for t under the classic standing model is 434 generations or 0.00029 coalescent units, which is much shorter than the time in which our selected populations coalesce. We caution against assigning too much value to these inferences, given the assumptions, but do find these approximations to be broadly useful.

### A.4 Standing variant model with a source population

When there are multiple selected populations and they do not follow a bifurcating tree structure, it is necessary to incorporate a model that has a source population for the standing variant to have self-consistent mean-centered covariance matrices.

Let population *l* be a selected population and the source of the beneficial allele. In all other populations, the beneficial allele is standing for time *t* generations at frequency *g* before the lineage returns to the source population where it still standing at frequency *g* (see Figure 11). We can define pairwise coancestry coefficients for all pairs of populations under this model. Let populations *i* and *j* represent populations that experience selection and population *k* be any unselected population.

Since population *l* is the source, its variance follows the same form as Equation 7.

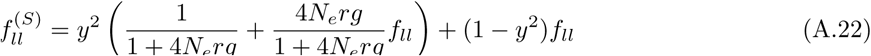

All other selected populations have a modified variance since lineages that fail to recombine off the beneficial background during the sweep and fail to coalesce or recombine during the standing phase return to the source population. Thus,

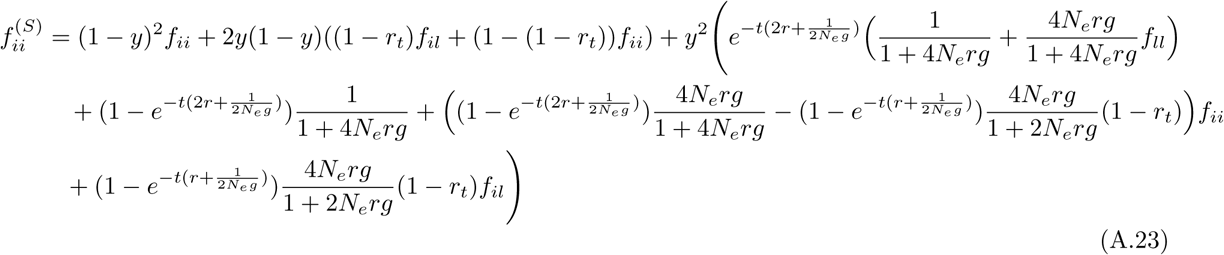

There is additional coancestry between pairs of selected populations. This takes a different form than Equation 9 as there since if either lineage fails to recombines off the beneficial background during the sweep or standing phase, the lineage will be in population *l*. For selected populations *i* and *j*, now

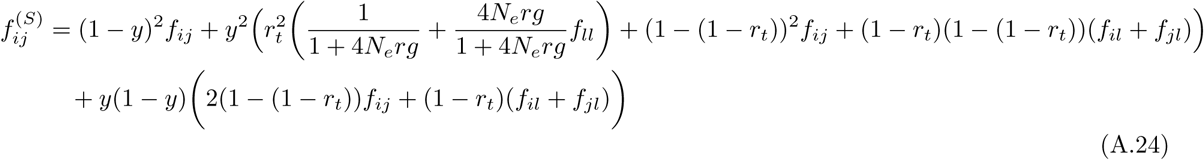

If either population is the source, *l* this reduces to

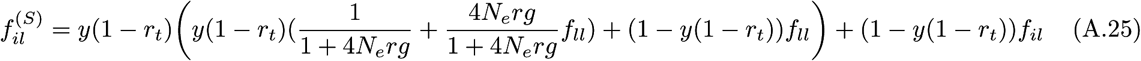

since if the lineage fails to recombines off the beneficial background in population *i*, it is back in population *l*. If the lineage in *l* is still on the beneficial background after the sweep and the initial *t* generations of standing, they can coalesce during the standing phase in population *l*. Else, the lineages will coalesce neutrally in population *l*. However, if the lineage sampled in population *i* does not return to the source population (i.e. it recombines during the sweep or standing phase of *t* generations), the lineages can coalesce with neutral probability *f*_*il*_.

Lastly, we must incorporate the impact linked selection has on the coancestry between lineages sampled from any pair of non-source selected population *i* and non-selected population *k*.

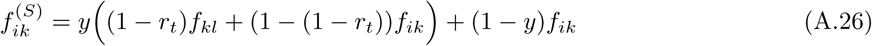

Since lineages that do not recombine off the beneficial background in population *i* go back into the source population *l*, non-selected populations may now have more or less coancestry with population *i* depending on whether *l* is neutrally has more or less coancestry with population *l*, respectively.

It may be possible to extend these models to allow the source population to be an unsampled population, *u*. In this case, we need information about how our unsampled source is related to our sampled populations. Specifically, we have *f*_*iu*_ and *f*_*uu*_ terms in the coancestry coefficients of any selected population *i* as well as *f*_*iu*_, *f*_*ju*_, and *f*_*uu*_ for coancestry between any selected population pairs *i* and *j* and *f*_*kl*_ for unselected populations *k*. More work is needed to address this problem. It is possible to use all sampled populations, including non-selected populations, as proxies for the unsampled source to give us information about which sampled population our unsampled source is more closely related to. Additionally, if we assume the unsampled population is distantly related to our sampled populations, such that they span the root, the coancestry between *u* and any other sampled population will be 0.

### A.5 Migration model: more than two non-source selected populations

In the main text, we consider two selected populations *i* and *j* where population *i* is the source of the beneficial allele. We need to extend this model when we have more than two non-source selected populations. Specifically, we need to define coancestry coefficients between selected non-source pairs. Now, let population *l* be a selected population and the source of the beneficial allele.

The coancestry between non-source selected populations is affected by migration as there is some probability or either or both lineage failing to recombine off the beneficial background of the sweep and to migrate back into population *l*. Thus, for selected populations *i* and *j*,

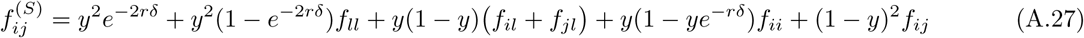

If *l* is either population *i* or *j*, this reduces to Equation 13, up to a factor of 2*δ* as now only one population experiences the delay, *δ*, as the other is the source. Thus, Equation 13 is more accurate for defining the coancestry coefficient between the source and selected populations. Equation 12 holds for the coancestry within all non-source selected population and Equation 14 for all non-selected and non-source selected population pairs. Lastly, again, we assume the source coancestry within the source population *l* follows that of an independent sweep from new mutation (Equation 4).

Similar to the standing variant model with a source population above, we can think about extending this migration model to allow the source population to be unsampled. More work is needed to address the same issues related to estimating coancestry coefficients for unsampled populations.

## S1 Single pulse of migration models

We also considered models of a single pulse of migration. We solve for 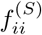 and 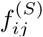 for the bounds on the time during which the beneficial allele could migrate: (1) “instantly” after the beneficial allele arises in population *i* and (2) after the beneficial allele reaches fixation in the population *i*.

### S1.1 Beneficial allele migrates instantly after it arises in population *i*

In this case, we are specifying the pulse of migration from population *i* into population *j* occurs sufficiently soon enough after the sweep began such that the entire haplotype the beneficial mutation arises on in population *i* migrates to population *j* (i.e. there is no time for recombination to occur). This case gives us results for an extreme of a single pulse of migration may not be particularly relevant as the spread of the beneficial allele into population *j* will likely only occur after it has reached a sufficiently high frequency in population *i* as it may be lost due to drift. However, these results aid in our intuition of this model.

As the beneficial allele originates in population *i*, again,

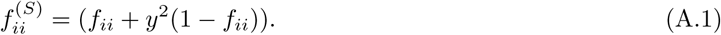

The probability of two lineages in the recipient population, *j*, coalescing before reaching the ancestral population is now

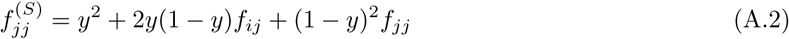

Here, both lineages can fail to recombine off the sweep (w.p. *y*^2^) and therefore coalesce with probability 1. Exactly one lineage can recombine off the sweep (w.p. 2*y*(1 − *y*)) and therefore the two lineages can only coalesce in the shared drift phase (w.p. *f*_*ij*_) as the lineage that does not recombine off the sweep migrates into population *i*. Both lineages can recombine off the sweep (w.p. (1 − *y*)^2^) and then can coalesce in population *j* before they reach the ancestral population.

The probability of two lineages drawn from each population coalescing before reaching the ancestral population is

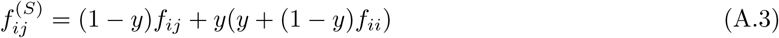

In this case, if the lineage in population *j* recombines off the sweep (w.p. 1 − *y*), the two lineages can only coalesce in the shared drift phase (w.p. *f*_*ij*_) before reaching the ancestral population. If the lineage in population *j* fails to recombine off the sweep (w.p. *y*), it migrates back to population *i* and will be forced to coalesce with the lineage in population *i* if it also failed to recombine, else they will coalesce neutrally in population *i*.

### S1.2 Beneficial allele migrates after it reaches fixation in population *i*

For the coancestry coefficient for population *j*, the logic follows from that of when the pulse of migration happens instantly. However in deriving the coancestry coefficient between populations *i* and *j*, in the case where the lineage sampled from population *j* fails to recombine off the sweep and migrates back to population *i*, which happens with probability y, it is like we have two lineages sampled in population *i*. Now, both could either fail to recombine off the sweep and coalesce with probability 1 or one or both could recombine off the sweep and coalesce neutrally in population *i*. This can be written as

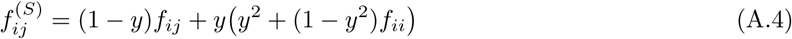

Together, these results characterize the other end point of a single pulse of migration spreading the beneficial allele to the recipient population.

## S2 Forward in time derivation examples

For the forward in time results we utilize Gillespie’s (2000) psuedohittchiking approximation with the incorporation of recombination to model the variance in the change in neutral allele frequencies due to a selective sweep (Δ*_S_x_i_* for population *i*). A new beneficial mutation will arise on the same background as a neutral allele with probability equal to its frequency in the population, *x*. In the case no crossing over occurs and the new mutation sweeps to fixation, the neutral allele frequency after the hitchhiking event, *x*′, will either be 1 with probability *x* or 0 with probability 1 − *x*. Therefore,

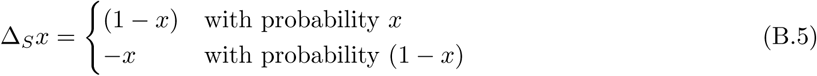

thus 𝔼[Δ*_S_x*] = 0 and Var[Δ*_S_x*] = *x*(1 − *x*).

Recombination can be incorporated into this model, allowing the neutral allele to stop hitchhiking before it reaches fixation. The frequency of the haplotype on which the favorable mutation arises will increase to *y* and all other alleles will have their frequencies reduced by 1 − *y*. So, if the favorable allele appears on the same background of our neutral allele, which happens with probability *x*, *x*′ = (1 − *y*)*x* + *y*. Else, with probability 1 − *x*, *x*′ = (1 − *y*)*x*. Therefore,

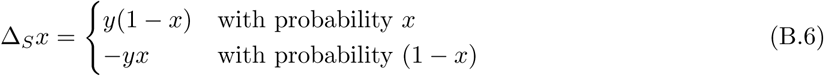

thus with recombination, 𝔼[Δ*_S_x*] = 0 and Var[Δ*_S_x*] = *y*^2^*x*(1 − *x*).

We can break down the changes in allele frequencies in the two populations from the ancestral allele frequency *E* into three components if we assume the independent drift in each population after the sweep is negligible: the change due to (1) shared drift between populations *i* and *j* before they split (Δ*_N_ x_ij_*), (2) independent drift in each population before the sweep (Δ*_N_ x_i_* and Δ*_N_ x_j_*), and (3) the selective sweep occurring in each population (Δ*_S_x_i_* and Δ*_S_x_j_*).

Define 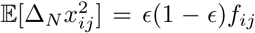 and 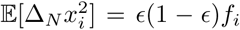 for population *i*. The total amount of variance in a neutral allele frequency for the *i*th population is defined as ϵ (1 − ϵ)*f*_*ii*_ which we approximate as *E*(1 − *E*)(*f*_*ij*_ + *f*_*i*_). This only holds if we assume the time intervals are short relative to drift so that these terms act additively. If this is not the case, the 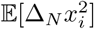 is no longer the probability that two alleles drawn from population *i* before the sweep begins are identical by descent with reference to the ancestral population with neutral allele frequency *E*, but rather with reference to the population before the split into populations *i* and *j* with neutral allele frequency *x*_*ij*_. A more careful treatment of these parameters could be done to relax this assumption, and follows naturally in a coalescent setting.

From a forward in time perspective, we can solve for Var[Δ*x*_*i*_], Var[Δ*x*_*j*_], and Cov[Δ*x*_*i*_, Δ*x*_*j*_] with Δ*x*_*i*_ =Δ*_N_ x_ij_* + Δ*_N_ x_i_* + Δ*_S_x_i_*. Assuming drift terms are independent of each other, we are left with the following expressions

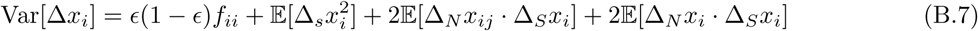

and

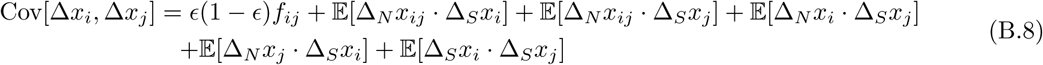

### S2.1 Independent sweep model

In the case of independent sweeps where there is no gene flow between populations, many terms in Equations B.7 and B.8 equal zero since the sweeps are independent. For the variances, we are left with

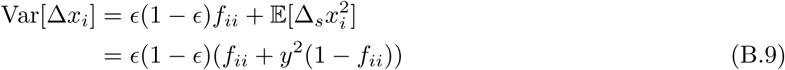

The covariance in allele frequencies between populations *i* and *j*, is simply what we would expect under neutrality.

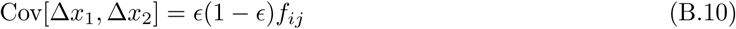

### S2.2 Shared sweeps via migration

The migration models better exemplifies these forward in time calculations. We demonstrate the calculations of Var[Δ*x*_*j*_] and Cov[Δ*x*_*i*_, Δ*x*_*j*_] for pulse of migration models specified in Supplement S1.

#### S2.2.1 Beneficial allele migrates instantly after it arises in population *i*

The background on which the beneficial mutation arises depends on the neutral allele frequency in population *i* before the sweep, *x*_*i*_. We are specifying the pulse of migration from population *i* into population *j* occurs sufficiently soon enough after the sweep began such that the entire haplotype the beneficial mutation arises on in population *i* migrates to population *j* (i.e. there is no time for recombination to occur). Now Δ*_S_x_j_* depends on the neutral allele frequency in population *i* before the sweep.

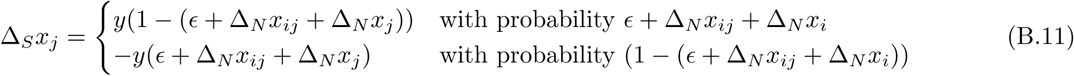

As the beneficial allele originates in population *i*, again,

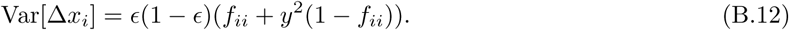

Now Δ*_S_x_j_* depends on *x*_*i*_, 𝔼[Δ*_N_ x_i_* Δ*_S_x_j_*], 𝔼[Δ*_S_x_i_* Δ*_S_x_j_*], and 𝔼[Δ*_N_ x_ij_* Δ*_S_x_j_*] are no longer zero. So,

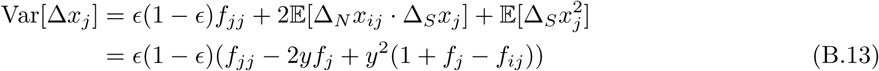

and

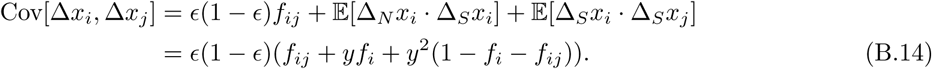

This result is the same as Equation A.3 if the assumption about drift being additive holds such that *f*_*ii*_ = *f*_*i*_ + *f*_*ij*_.

#### S2.2.2 Beneficial allele migrates after it reaches fixation in population *i*

Now, the frequency of a neutral allele in population *i* after the sweep has occurred is

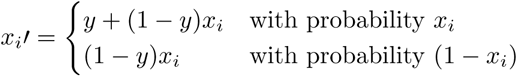

Fixing that the migration from population *i* into *j* occurs after the sweep has finished in population *i*,

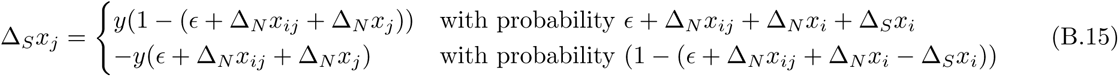

This can also be written as

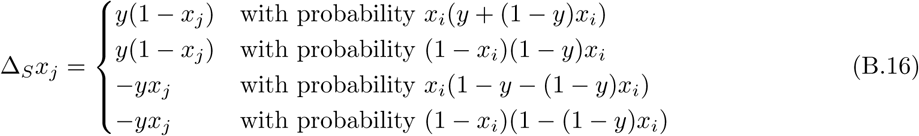

Here, the first case is that the beneficial allele arises on the same background as our neutral allele in population *i* and then is the haplotype that migrates into population *j*. The probability of the haplotype migrating is equal to its frequency in the population. The third case also includes the beneficial allele arising on the same background as our neutral allele, but the other haplotype migrates. The second and fourth cases are when the beneficial mutation arises on the other background as our neutral allele. In the second case, the haplotype containing our neutral allele migrates after the sweep and in the fourth, the other haplotype migrates.

The variance within population *i* and population *j* are the same as in the case of the beneficial allele migrating instantly. The only term changed by specifying that the pulse of migration happens after the sweep is 𝔼[Δ*_S_x_i_* Δ*_S_x_j_*] which is now ϵ (1 -ϵ)*y*^3^(1 − *f_jj_*). So,

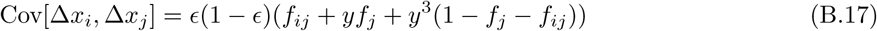

## S3 mssel input for simulations

**Independent sweep model.** mssel input for all independent sweep model is of the following form with different trajectory files for each *s*,

~~~
./mssel 40 1000 20 20 ind_sel0.1_stochastic.traj 0 -t 2000 -r 2000 10000
-I 4 10 0 0 10 0 10 10 0 -ej 0.05 3 4 -ej 0.05 2 1 -ej 0.07 4 1
~~~

**Standing variation model.**

~~~
./mssel 40 1000 20 20 sv_sel0.01_g0.001_t50_stochastic.traj 0 -t 200 -r 120 10000
-I 4 10 0 0 10 0 10 10 0 -ej 0.0346 2 1 -ej 0.0346 3 4 -ej 0.03575 4 1
./mssel 40 100 20 20 sv_sel0.01_g0.001_t250_stochastic.traj 0 -t 200 -r 200 10000
-I 10 0 0 10 0 10 10 0 -ej 0.039 3 4 -ej 0.039 2 1 -ej 0.0408 4 1
./mssel 40 1000 20 20 sv_sel0.01_g0.001_t500_stochastic.traj 0 -t 200 -r 200 10000
-I 10 0 0 10 0 10 10 0 -ej 0.04 2 1 -ej 0.04 3 4 -ej 0.047 4 1
./mssel 40 100 20 20 sv_sel0.01_g0.001_t1000_stochastic.traj 0 -t 200 -r 200 10000
-I 10 0 0 10 0 10 10 0 -ej 0.04 3 4 -ej 0.04 2 1 -ej 0.0595 4 1
./mssel 40 1000 20 20 sv_sel0.01_g0.001_t5000_stochastic.traj 0 -t 200 -r 200 10000
-I 10 0 0 10 0 10 10 0 -ej 0.135 2 1 -ej 0.135 3 4 -ej 0.1595 4 1
~~~

We also simulated under two additional selection coefficients, *s* = [0.001, 0.05], keeping *t* = 500 and *g* = 0.001.

~~~
./mssel 40 100 20 20 sv_sel0.001_g0.001_t500_stochastic.traj 0 -t 200 -r 200 10000
-I 10 0 0 10 0 10 10 0 -ej 0.3455 3 4 -ej 0.3455 2 1 -ej 0.3578 4 1
./mssel 40 100 20 20 sv_sel0.05_g0.001_t500_Ne10000_stochastic.traj 0 -t 200 -r 200 10000
-I 10 0 0 10 0 10 10 0 -ej 0.00695 3 4 -ej 0.00695 2 1 -ej 0.01935 4 1
~~~

**Migration model.**

~~~
./mssel 40 1000 20 20 mig_sel0.01_mig1e-04_stochastic.traj 0 -t 200 -r 200 10000
-I 10 0 0 10 0 10 10 0 -ej 0.07 2 1 -ej 0.07 3 4 -ej 0.1 4 1
-em 0.059 3 2 0 -em 0 3 2 4
./mssel 40 1000 20 20 mig_sel0.01_mig0.001_stochastic.traj 0 -t 200 -r 200 10000
-I 10 0 0 10 0 10 10 0 -ej 0.07 2 1 -ej 0.07 3 4 -ej 0.1 4 1
-em 0.059 3 2 0 -em 0 3 2 40
./mssel 40 1000 20 20 mig_sel0.01_mig0.01_stochastic.traj 0 -t 200 -r 200 10000
-I 10 0 0 10 0 10 10 0 -ej 0.07 2 1 -ej 0.07 3 4 -ej 0.1 4 1
-em 0.059 3 2 0 -em 0 3 2 400
./mssel 40 1000 20 20 mig_sel0.01_mig0.1_stochastic.traj 0 -t 200 -r 200 10000
-I 10 0 0 10 0 10 10 0 -ej 0.07 2 1 -ej 0.07 3 4 -ej 0.1 4 1
-em 0.059 3 2 0 -em 0 3 2 4000
~~~

We also simulated under two additional selection coefficients, *s* = [0.005, 0.05], keeping *m* = 0.001.

~~~
./mssel 40 100 20 20 mig_sel0.05_mig0.001_stochastic.traj 0 -t 200 -r 200 10000
-I 10 0 0 10 0 10 10 0 -ej 0.021 2 1 -ej 0.021 3 4 -ej 0.03 4 1
-em 0.014 3 2 0 -em 0 3 2 40
./mssel 40 100 20 20 mig_sel0.005_mig0.001_stochastic.traj 0 -t 200 -r 200 10000
-I 10 0 0 10 0 10 10 0 -ej 0.12 2 1 -ej 0.12 3 4 -ej 0.17 4 1
-em 0.11 3 2 0 -em 0 3 2 40
~~~

## S4 Parametric-bootstrap simulation details

### S4.1 Copper tolerance in *Mimulus guttatus* specifics

Below are the input for the simulation runs to generate parametric bootstraps for the *Mimulus guttatus* analysis. We simulate with *N*_*e*_ = 7500, except for in the migration model where *N*_*e*_ = 30000 (to allow for smaller *ŝ*).

**Neutral model.**

~~~
./ms 194 100 -s 5723 -r 239.7203 169294 -I 4 62 42 40 50 -ej 0.057 4 1 -ej 0.056 2 1
-ej 0.085 3 1
~~~

**Independent mutations model**. 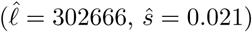

~~~
./mssel 194 100 102 92 mim_indMLE_comp.traj 87565.86 -s 5723 -r 239.7203 169294
-I 4 0 62 42 0 0 40 50 0 -ej 0.057 4 1 -ej 0.056 2 1 -ej 0.085 3 1
~~~

**Migration model.** 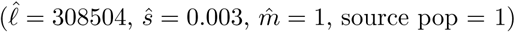

~~~
./mssel 194 100 102 92 mim_migMLE_comp_Ne30000.traj 93403.6 -s 5723 -r 958.8812 169294
-I 4 0 62 42 0 0 40 50 0 -ej 0.057 4 1 -ej 0.056 2 1 -ej 0.085 3 1
-em 0.04975 3 1 0 -em 0.0496 3 1 120000
~~~

**Standing variant with source model**. 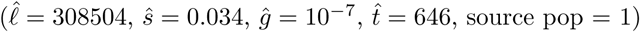

~~~
./mssel 194 100 102 92 mim_svSourceMLE_comp.traj 93403.6 -s 5723 -r 239.7203 169294
-I 4 0 62 42 0 0 40 50 0 -ej 0.057 4 1 -ej 0.056 2 1 -ej 0.085 3 1
-em 0.043 3 1 0.001 -em 0.045 3 1 0
~~~

### S4.2 Industrial pollutant tolerance in *Fundulus heteroclitus* specifics

Below are the input for the simulation runs to generate parametric bootstraps for the *Fundulus heteroclitus* analysis. We simulate with *N*_*e*_ = 1000 for all models.

**Neutral model.**

~~~
./ms 768 100 -s 66593 -r 214.4814 2470984 -I 8 96 96 98 100 100 86 94 98
-ej 0.0274276738490838 4 3 -ej 0.0344793500868448 3 1 -ej 0.0473737546397982 2 1
-ej 0.0529009970762367 6 1 -ej 0.060223521932099 5 1 -ej 0.0281723542369385 8 7
-ej 0.131042855088188 7 1
~~~

**Independent mutations model.** 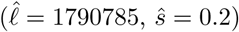

~~~
./mssel 768 100 380 388 indMLE_killi_Ne1000.traj 1789333 -s 66593 -r 214.4814 2470984
-I 8 96 0 0 96 98 0 0 100 100 0 0 86 94 0 0 98 -ej 0.0274276738490838 4 3
-ej 0.0344793500868448 3 1 -ej 0.0473737546397982 2 1 -ej 0.0529009970762367 6 1
-ej 0.060223521932099 5 1 -ej 0.0281723542369385 8 7 -ej 0.131042855088188 7 1
~~~

**Migration model.** 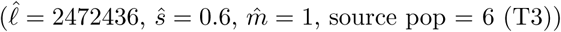

~~~
./mssel 768 100 380 388 mig_mle_Ne1000_killi.traj 2470984 -s 66593 -r 214.4814 2470984
-I 8 96 0 0 96 98 0 0 100 100 0 0 86 94 0 0 98 -ej 0.0274276738490838 4 3
-ej 0.0344793500868448 3 1 -ej 0.0473737546397982 2 1 -ej 0.0529009970762367 6 1
-ej 0.060223521932099 5 1 -ej 0.0281723542369385 8 7 -ej 0.131042855088188 7 1
-em 0.00614 8 6 0 -em 0.006 8 6 4000 -em 0.00614 2 6 0 -em 0.006 2 6 4000
-em 0.00614 4 6 0 -em 0.006 4 6 4000
~~~

**Standing variant with source model.** 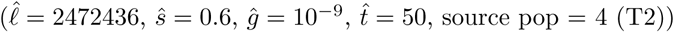

~~~
./mssel 768 100 380 388 sv_killiMLE_Ne1000.traj 2470984 -s 66593 -r 214.4814 2470984
-I 8 96 0 0 96 98 0 0 100 100 0 0 86 94 0 0 98 -ej 0.0274276738490838 4 3
-ej 0.0344793500868448 3 1 -ej 0.0473737546397982 2 1 -ej 0.0529009970762367 6 1
-ej 0.060223521932099 5 1 -ej 0.0281723542369385 8 7 -ej 0.131042855088188 7 1
-em 0.0243 8 4 0 -em 0.0240 8 4 0.0001 -em 0.0243 2 4 0 -em 0.0240 2 4 0.0001
-em 0.0243 6 4 0 -em 0.0240 6 4 0.0001
~~~

**Migration in North and independent mutation in South model.** (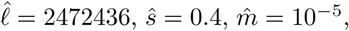

~~~
./mssel 768 100 380 388 migInd_mle_killi_Ne1000.traj 2470984 -s 66593 -r 214.4814 2470984
-I 8 96 0 0 96 98 0 0 100 100 0 0 86 94 0 0 98 -ej 0.0274276738490838 4 3
-ej 0.0344793500868448 3 1 -ej 0.0473737546397982 2 1 -ej 0.0529009970762367 6 1
-ej 0.060223521932099 5 1 -ej 0.0281723542369385 8 7 -ej 0.131042855088188 7 1
-em 0.01237 2 6 0 -em 0.0089 2 6 0.04 -em 0.01237 4 6 0 -em 0.0089 4 6 0.04
~~~

**Standing variation with source in North and independent mutation in South model.** 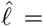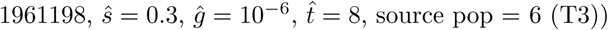

~~~
./mssel 768 100 380 388 svInd_killi_Ne1000.traj 1959746 -s 66593 -r 214.4814 2470984
-I 8 96 0 0 96 98 0 0 100 100 0 0 86 94 0 0 98 -ej 0.0274276738490838 4 3
-ej 0.0344793500868448 3 1 -ej 0.0473737546397982 2 1 -ej 0.0529009970762367 6 1
-ej 0.060223521932099 5 1 -ej 0.0281723542369385 8 7 -ej 0.131042855088188 7 1
-em 0.0195 2 6 0 -em 0.01925 2 6 0.0001 -em 0.0195 4 6 0 -em 0.01925 4 6 0.0001
~~~

## S5 Supplemental tables and figures

**Figure S1:**
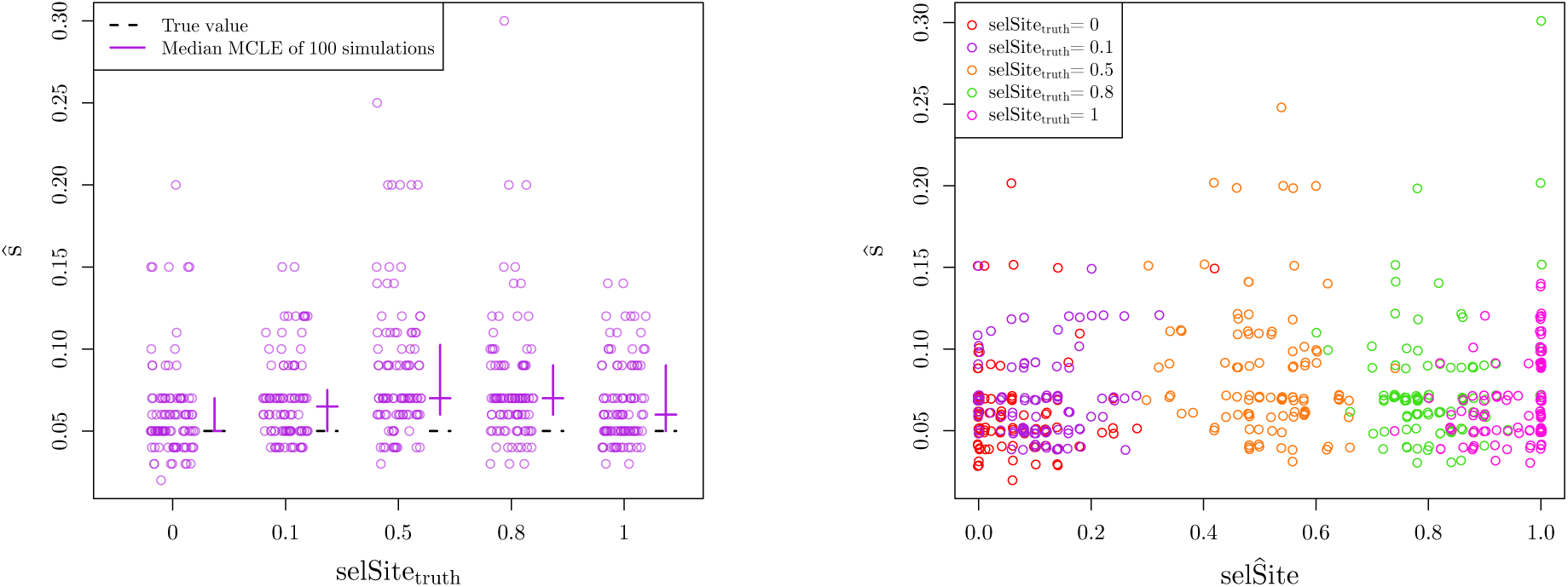
MCLE of **parameters** for **independent mutation** simulations **allowing selected site to vary**. (a) MCLE of **selection coefficients** as function of true location of selected site. Each location of selected site has 100 simulations under **independent mutation model** (10 chromosomes per population, Ne =100,000, s = 0.05). Crossbars indicate first and third quartiles with second quartiles (medians) as the horizontal line. The true values of the parameters are marked with dashed, black lines. (b) MCLE of **selection coeffcients** versus MCLE of **location of selected site**. True location of selected site is marked by color. Each location of selected site has 100 simulations under **independent mutation model** (10 chromosomes per population, Ne = 100,000, s =0.05)

**Figure S2:**
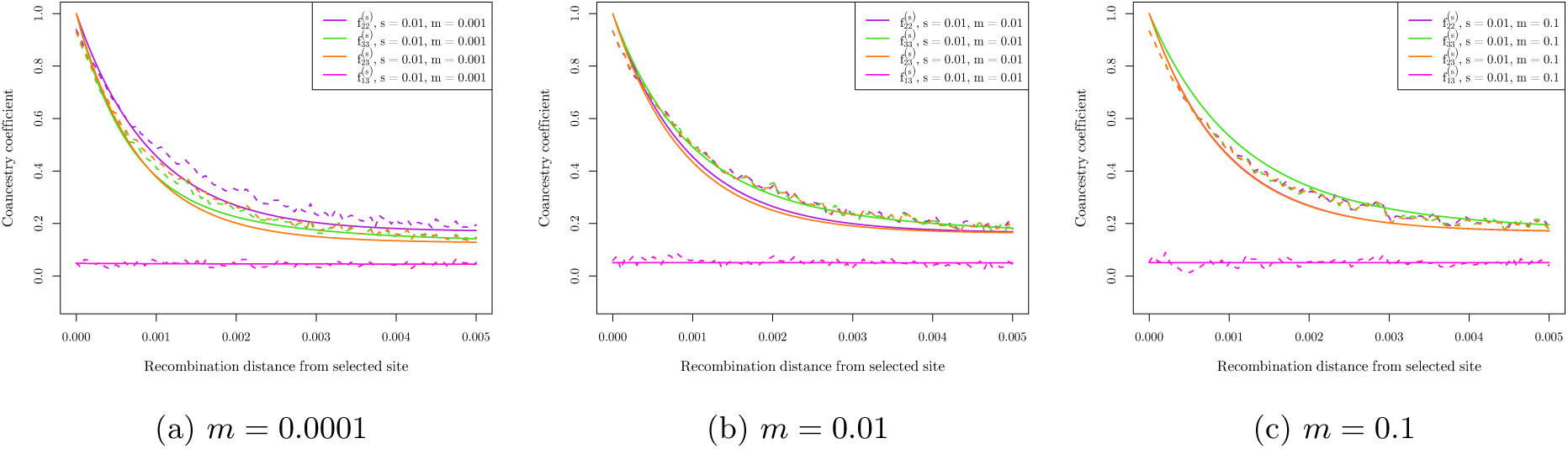
Average coancestry coefficient values for migration simulations with various *m*, across 100 runs of simulations for each of 100 bins of distance away from the selected site, showing the migration rate parameter does not have a large effect on both expectations (solid lines) and simulation results (dashed lines). For all simulations, *s* = 0.01, *N*_*e*_ = 10, 000, and the source of the beneficial allele is population 2.

**Figure S3:**
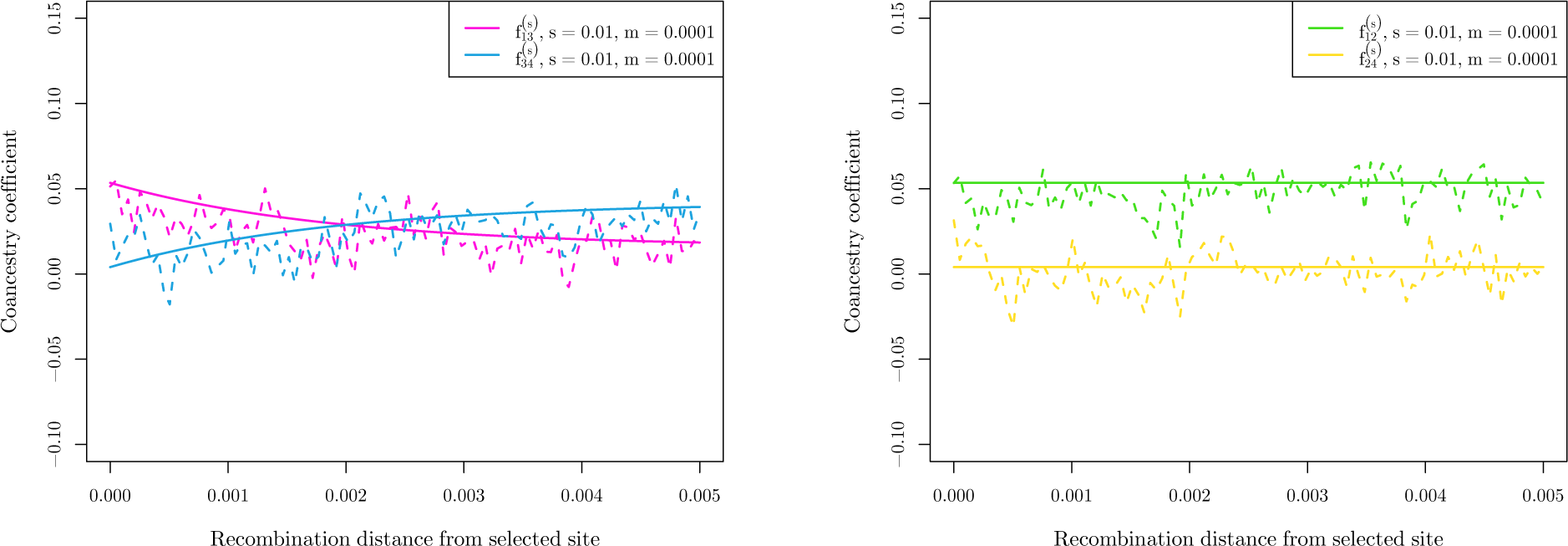
Average coancestry coefficient values for migration simulations across 100 runs of simulations for each of 100 bins of distance away from the selected site, between source and recipient populations and non-selected populations (*s* = 0.01, *m* = 0.001, *N*_*e*_ = 10,000). (a) Average coancestry coe_cient values for migration simulations across 100 runs of simulations for each of 100 bins of distance away from the selected site, between recipient population (3) and non-selected populations (1 and 4). (b) Average coancestry coe_cient values for migration simulations across 100 runs of simulations for each of 100 bins of distance away from the selected site, between source population (2) and non-selected populations (1 and 4).

**Figure S4:**
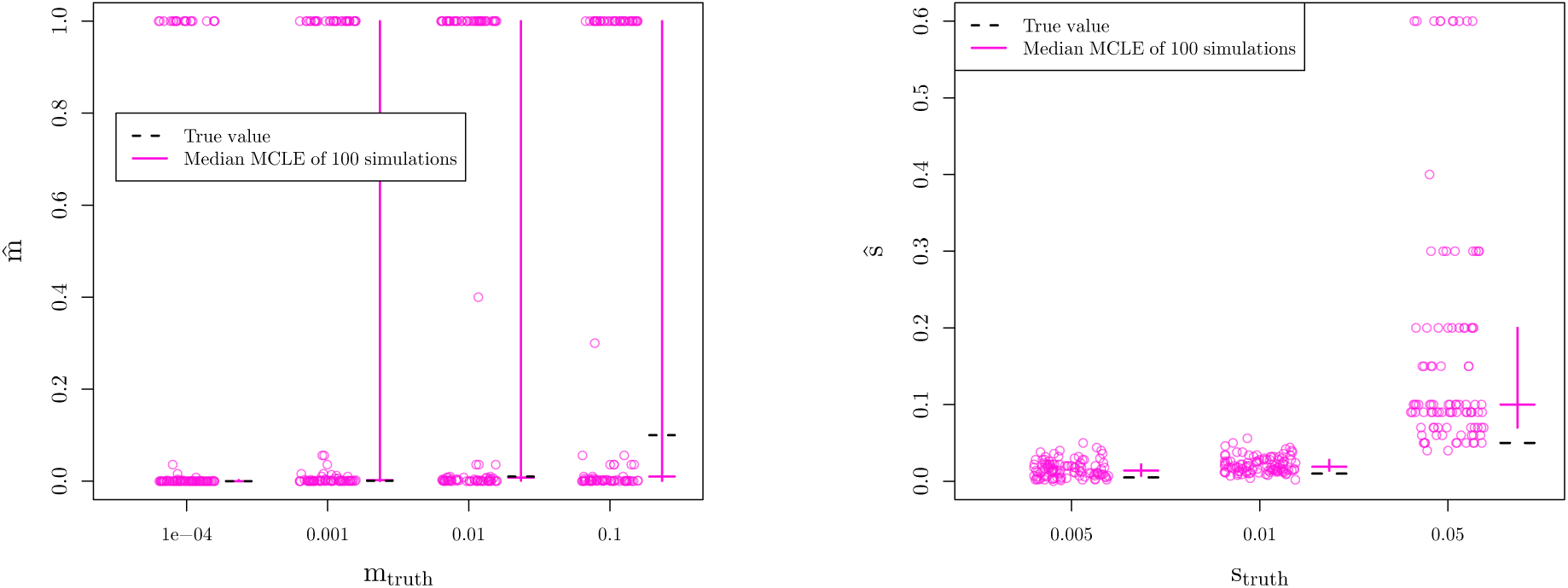
MCLE of **parameters** for **migration model** simulations. We vary the true value of the parameter used for simulations along the x-axis and show the MCLE for each of 100 simulations (points). Crossbars indicate first and third quartiles with second quartiles (medians) as the horizontal line. The true values of the parameters are marked with dashed, black lines. (a) MCLE of **migration rates** for 100 simulations under **migration model** (10 chromosomes per population, Ne =10,000, s = 0.01) (b) MCLE of **selection coeffcients** for 100 simulations under **migration model** (10 chromosomes per population, Ne = 10,000, m = 0.001)

**Figure S5:**
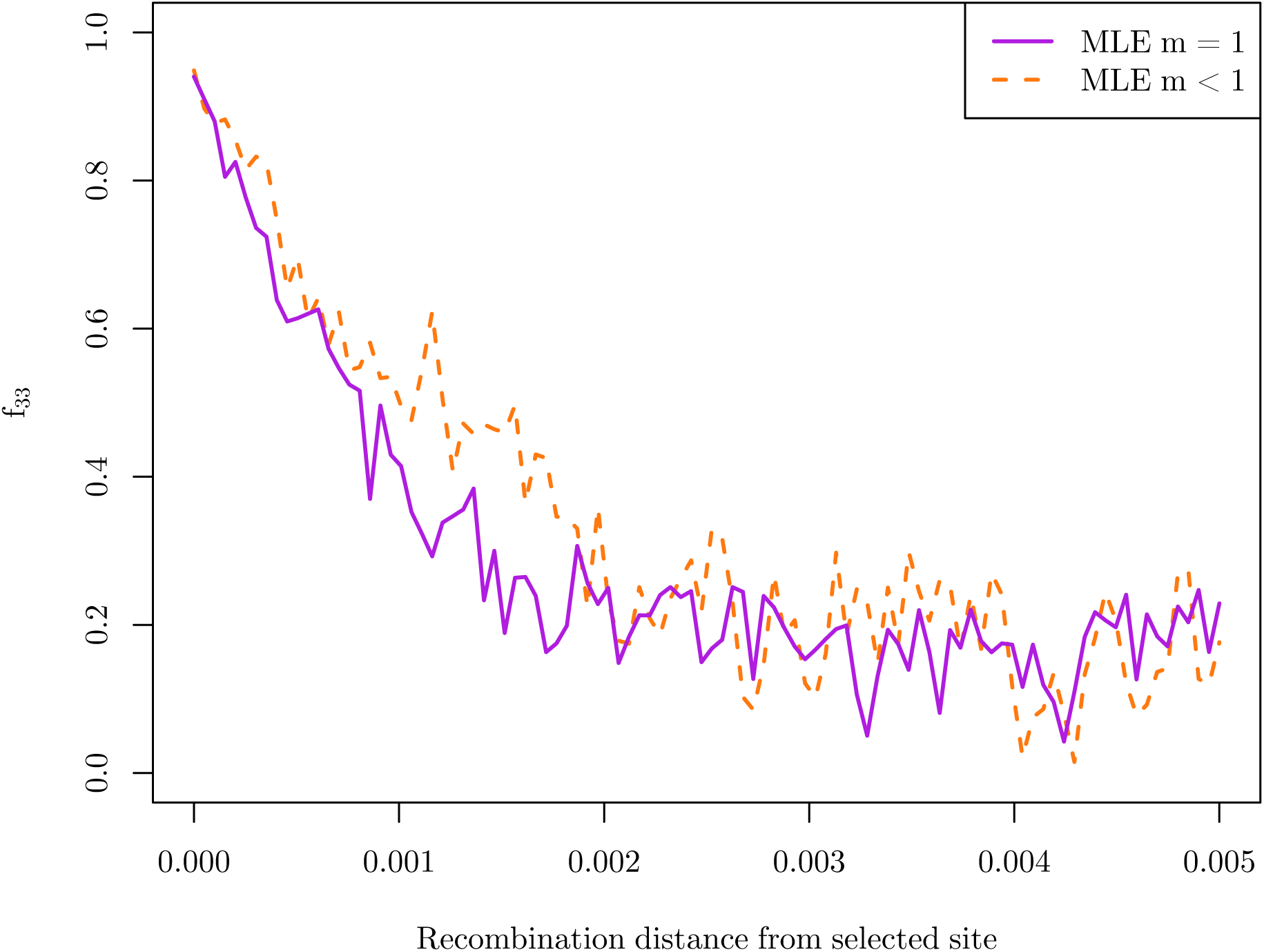
Coancestry coefficient for the recipient population as a function of recombination distance from the selected site, partitioned into simulations with MCLE for *m* = 1 and *m* < 1 (*s* = 0.01, *m* = 0.001, *N*_*e*_= 10,000).

**Figure S6:**
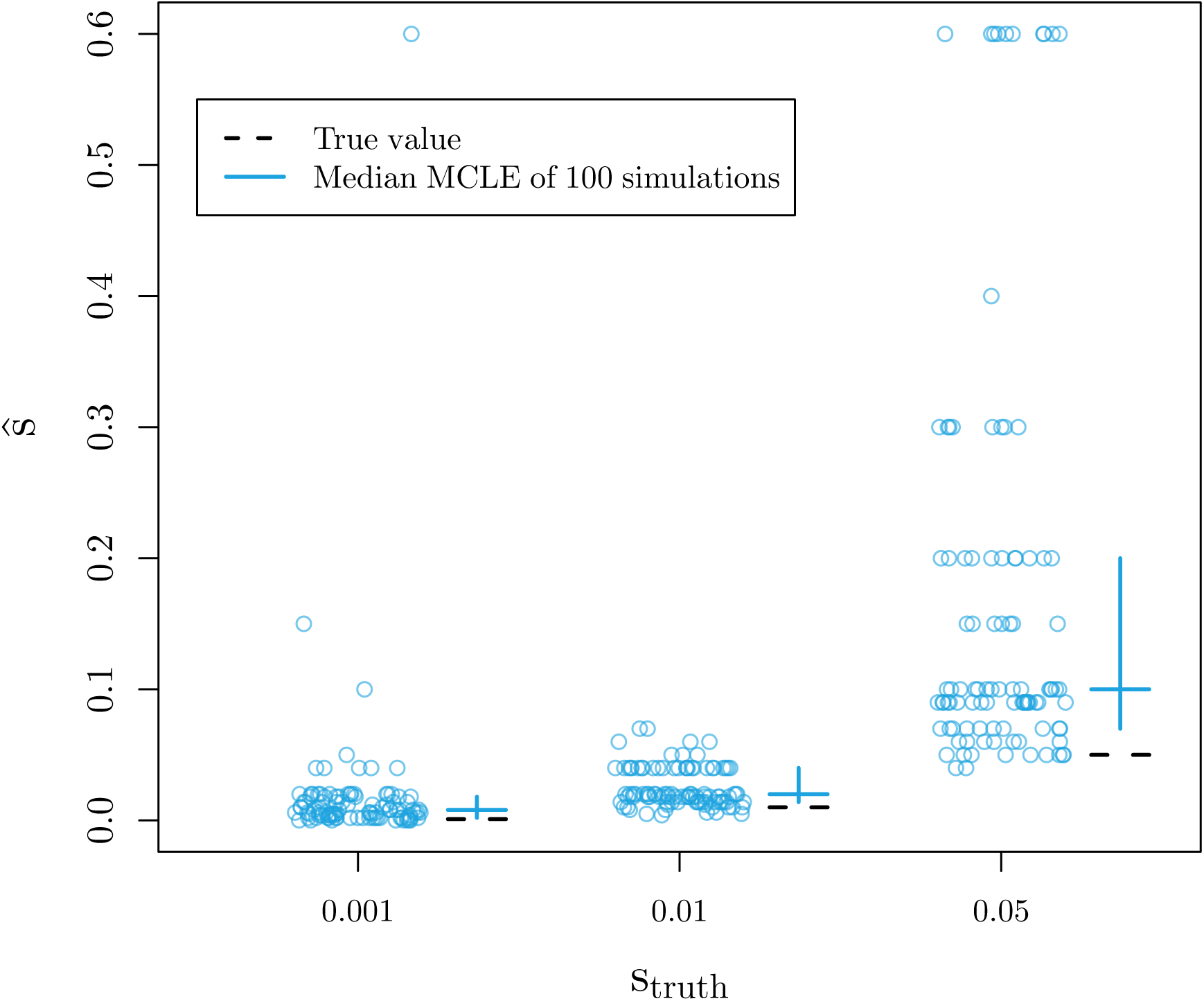
MCLE of **selection coefficients** for 100 simulations under**standing variant model** (10 chromosomes per population, *N*_*e*_ = 10,000, *t* = 500, *g* = 0.001). We vary the true value of the parameter used for simulations along the x-axis and show the MCLE for each of 100 simulations (points). Crossbars indicate first and third quartiles with second quartiles (medians) as the horizontal line. The true values of the parameters are marked with dashed, black lines.

**Figure S7:**
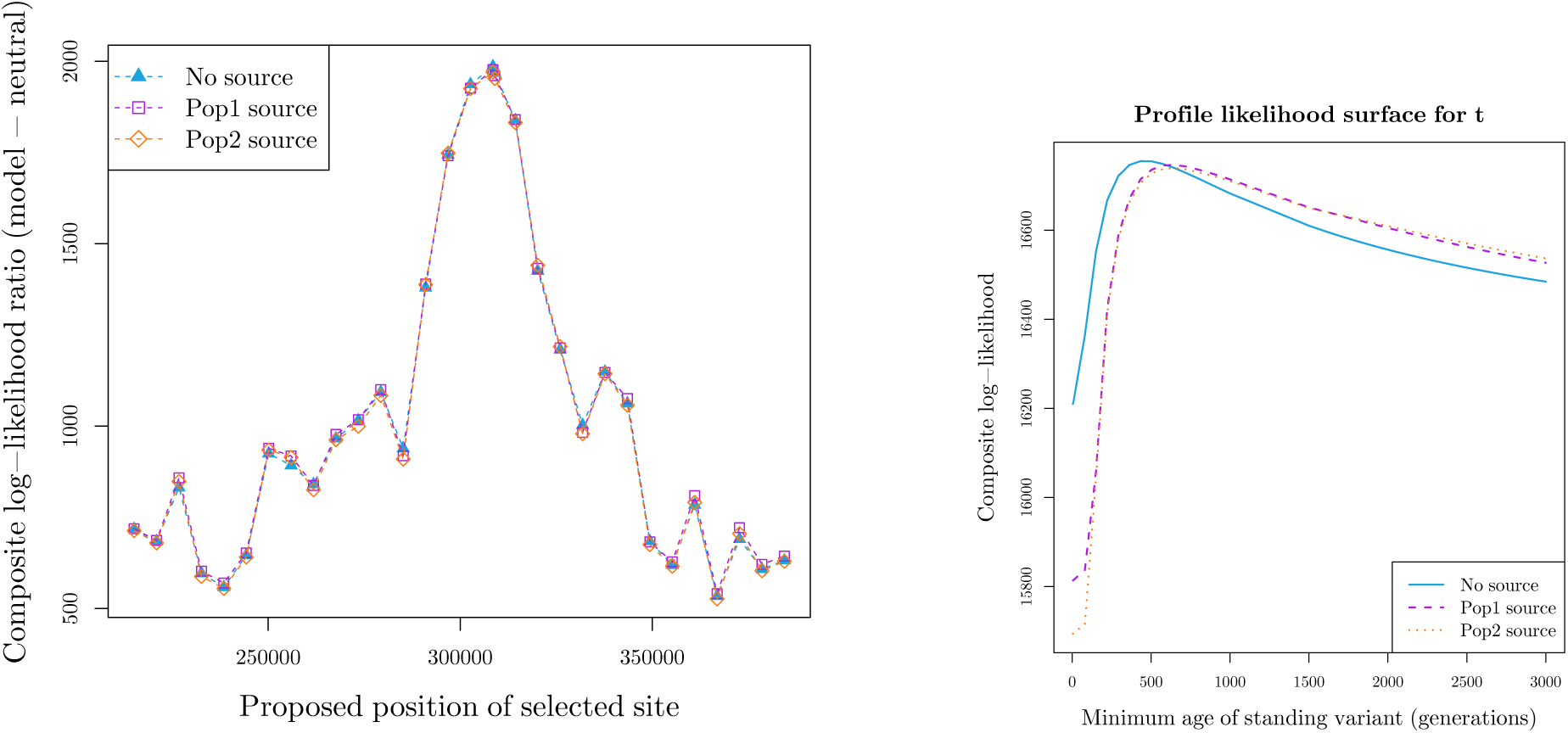
Inference results for standing variant model applied to *Mimulus* data using both original standing variant model and more complex model where a source population is specified. In this case, the composite log-likelihoods do not change, but the parameter estimates do. We obtain higher MCLE for *t* when a source is specified (646 generations) compared to the original no source model (434 generations). This fits our expectation as *t* has slightly different interpretations under the two models. (a) Composite log-likelihood for standing variation model with no source speci_ed and both selected populations as potential sources, as a function of the proposed selected site. (b) Profile composite log-likelihood of the minimum age of the standing variant for standing variant model with no source speci_ed and both selected populations as potential sources.

**Figure S8:**
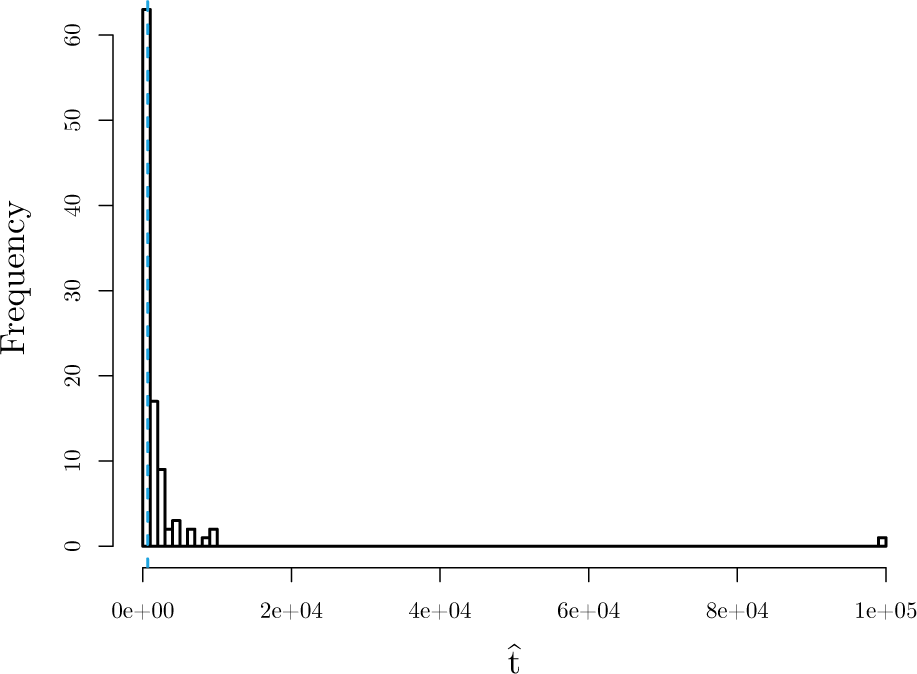
Histogram of MCLE for mimimum age of the standing variant (*t*) for 100 simulations under MCLE of standing variation with source model for *Mimulus guttatus*. MCLE from actual data is shown with dashed, blue line.

**Figure S9:**
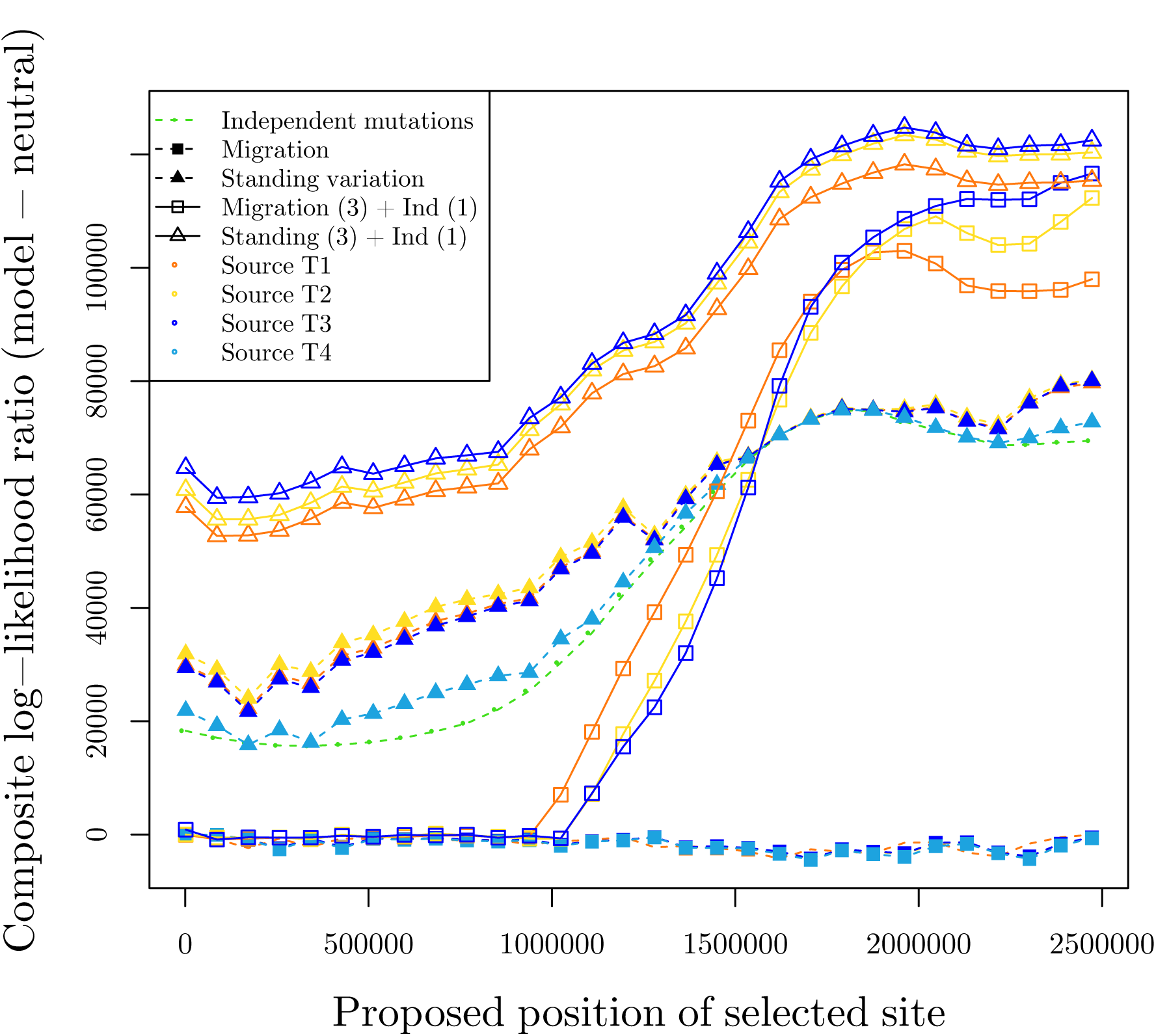
Composite log-likelihood for *Fundulus heteroclitus* pollutant tolerance adaptation on Scaffold9893, showing all possible sources for models with migration and standing variant model, as a function of the proposed selected site.

**Figure S10:**
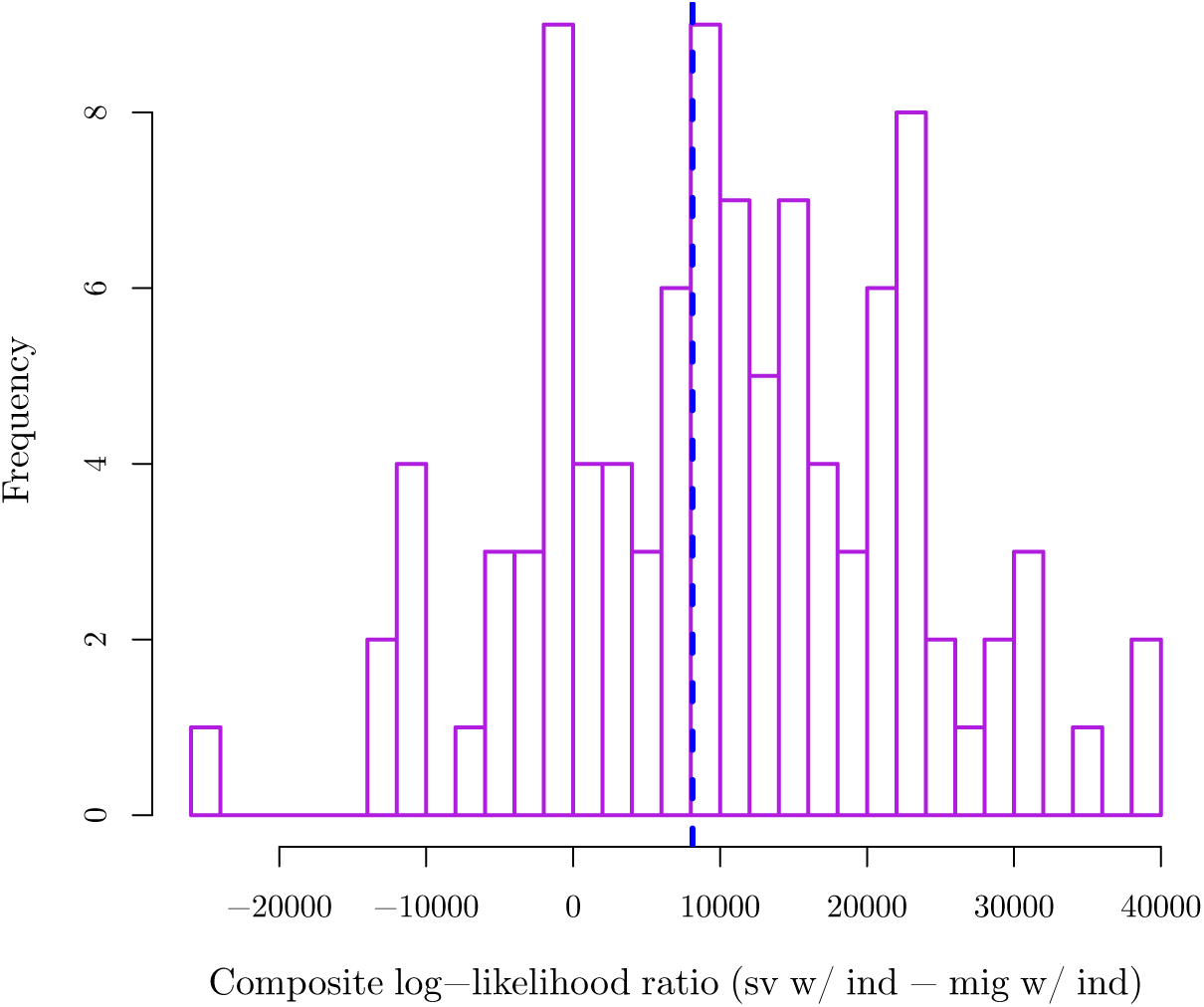
Histogram of composite log-likelihood ratio for 100 simulations under MCLE of migration in Northern tolerant populations and independent mutation in Southern tolerant populations for *Fundulus heteroclitus* (standing variation with T3 as source in Northern tolerant populations and independent mutation in Southern tolerant populations migration in Northern tolerant populations and independent mutation in Southern tolerant populations). Observed value from actual data is shown with dashed, blue line.

**Figure S11:**
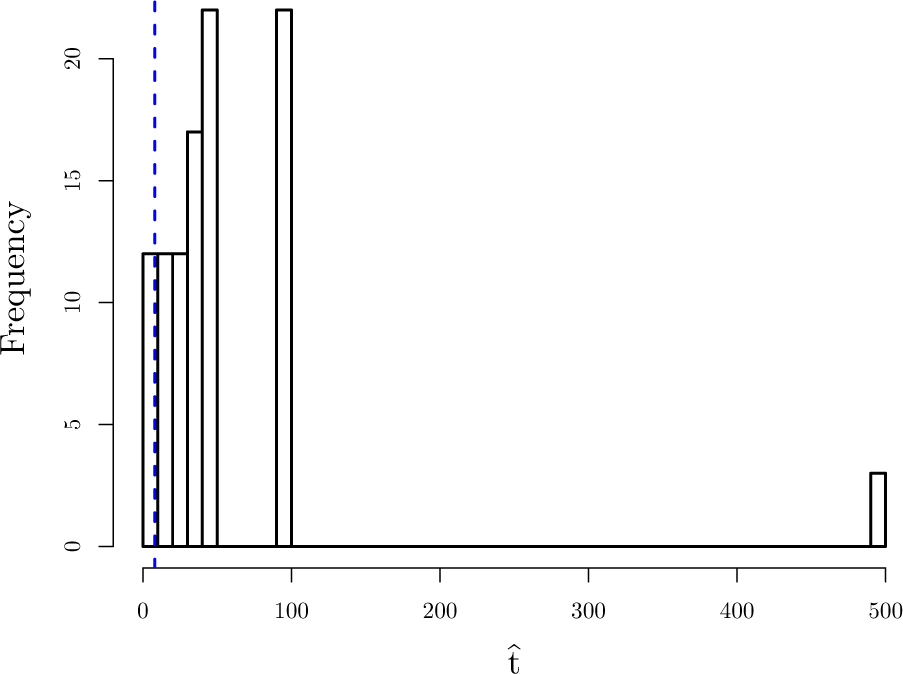
Histogram of MCLE for mimimum age of the standing variant (*t*) for 100 simulations under MCLE of standing variation with T3 as source in Northern tolerant populations and independent mutation in Southern tolerant populations for *Fundulus heteroclitus*. MCLE from actual data is shown with dashed, blue line.

**Figure S12:**
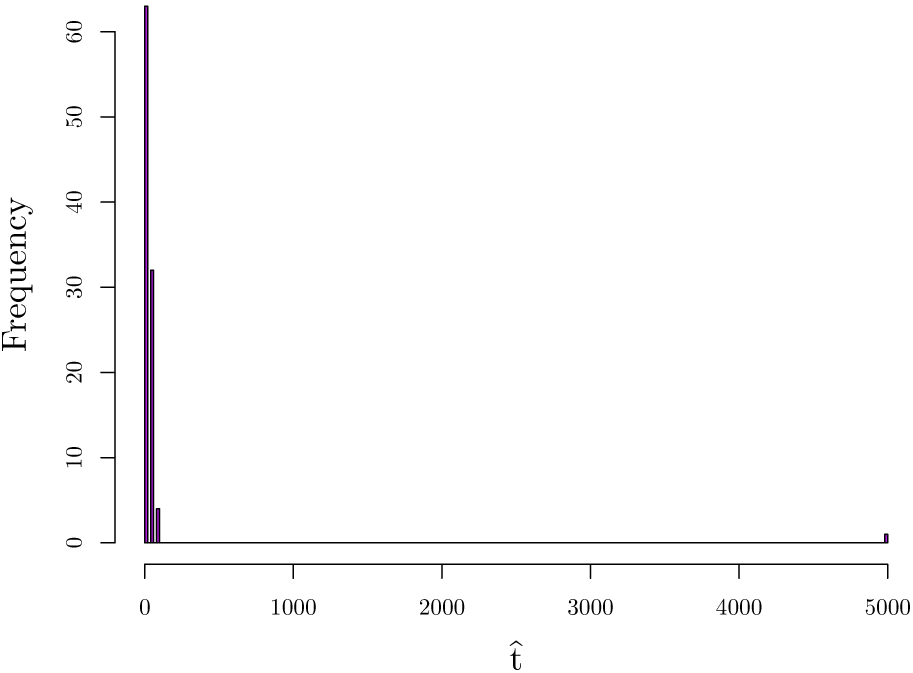
Histogram of MCLE for mimimum age of the standing variant 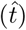 for 100 simulations under MCLE of migration with T3 as source in Northern tolerant populations and independent mutation in Southern tolerant populations for *Fundulus heteroclitus*.

**Table S1:** Parameter spaces for composite-likelihood calculations for simulated datasets

|  |  |
| --- | --- |
| Position of selected site | 0 |
| $s$ | $10^{-4}$, $5 \times 10^{-4}$, $10^{-3}$, $2 \times 10^{-3}$, $4 \times 10^{-3}$, $5 \times 10^{-3}$, $6 \times 10^{-3}$, $8 \times 10^{-3}$, 0.01, 0.012, 0.014, 0.018, 0.02, 0.03, 0.04, 0.05, 0.06, 0.07, 0.09, 0.1, 0.11, 0.12, 0.14, 0.15, 0.2, 0.25, 0.3, 0.35, 0.4, 0.5, 0.6 |
| $t$ | 0, 5, 15, 25, 40, 50, 60, 75, 100, 150, 200, 250, 300, 350, 400, 450, 500, 550, 600, 650, 700, 750, 800, 900, 1000, 1200, 1500, 1800, 2000, 2500, 3000, 3500, 4000, 4500, 5000, 5500, 6000, 6500, 7000, 7500, 8000, 9000, $10^4$, $1.5 \times 10^5$, $2 \times 10^5$, $3 \times 10^5$, $5 \times 10^5$, $7 \times 10^5$, $9 \times 10^5$, $10^5$, $10^6$ |
| $g$ | $10^{-3}$ |
| $m$ | $10^{-5}$, $10^{-4}$, $5 \times 10^{-4}$, $10^{-3}$, $5 \times 10^{-3}$, 0.01, 0.2, 0.5, 0.9, 1 |
| Migration source population | 2 |

**Table S2:**
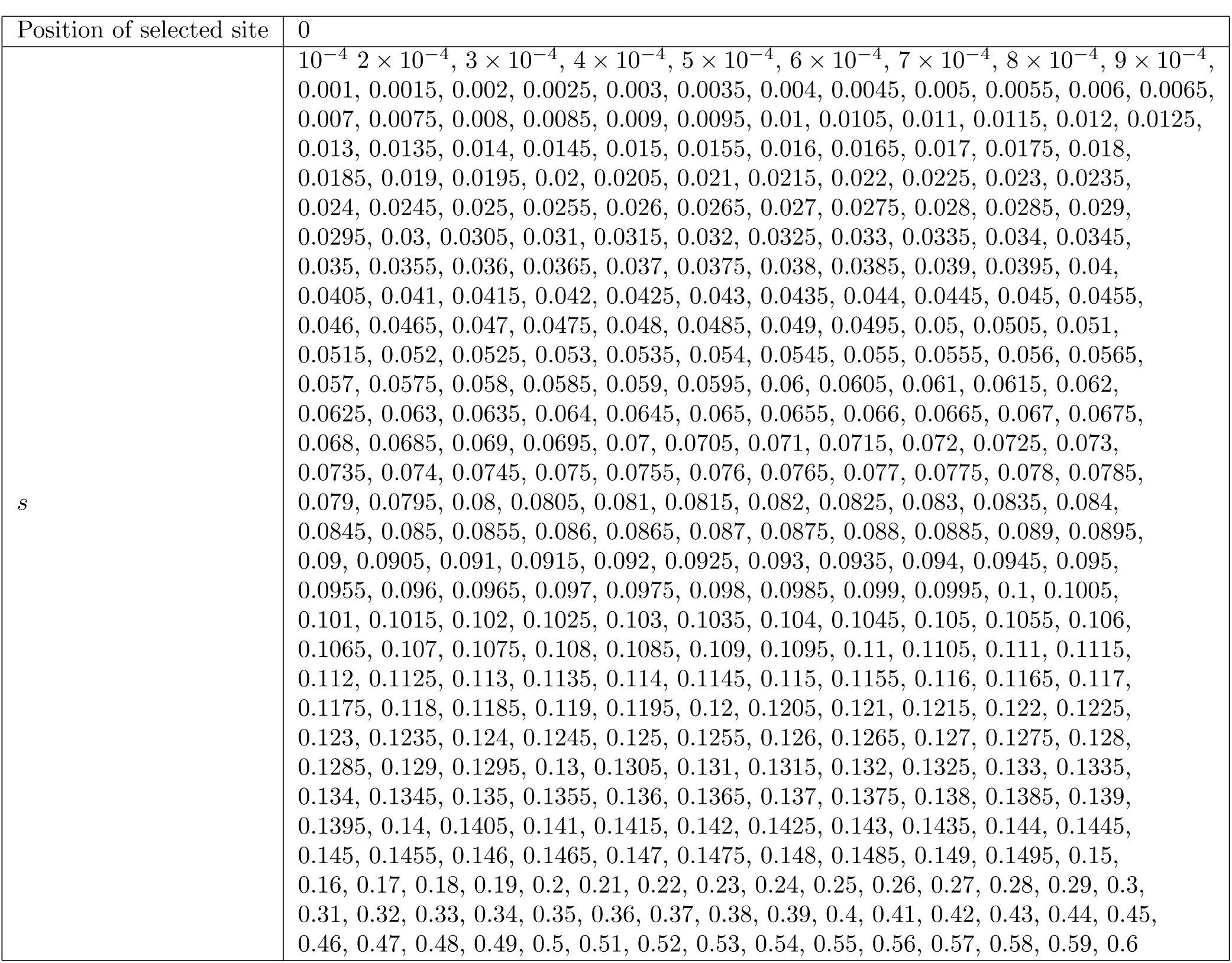
Parameter spaces for composite-likelihood calculations for independent sweep model simulations

|  |  |
| --- | --- |
| Position of selected site | 0 |
| $s$ | $10^{-4}, 2 \times 10^{-4}, 3 \times 10^{-4}, 4 \times 10^{-4}, 5 \times 10^{-4}, 6 \times 10^{-4}, 7 \times 10^{-4}, 8 \times 10^{-4}, 9 \times 10^{-4},$<br>0.001, 0.0015, 0.002, 0.0025, 0.003, 0.0035, 0.004, 0.0045, 0.005, 0.0055, 0.006, 0.0065,<br>0.007, 0.0075, 0.008, 0.0085, 0.009, 0.0095, 0.01, 0.0105, 0.011, 0.0115, 0.012, 0.0125,<br>0.013, 0.0135, 0.014, 0.0145, 0.015, 0.0155, 0.016, 0.0165, 0.017, 0.0175, 0.018,<br>0.0185, 0.019, 0.0195, 0.02, 0.0205, 0.021, 0.0215, 0.022, 0.0225, 0.023, 0.0235,<br>0.024, 0.0245, 0.025, 0.0255, 0.026, 0.0265, 0.027, 0.0275, 0.028, 0.0285, 0.029,<br>0.0295, 0.03, 0.0305, 0.031, 0.0315, 0.032, 0.0325, 0.033, 0.0335, 0.034, 0.0345,<br>0.035, 0.0355, 0.036, 0.0365, 0.037, 0.0375, 0.038, 0.0385, 0.039, 0.0395, 0.04,<br>0.0405, 0.041, 0.0415, 0.042, 0.0425, 0.043, 0.0435, 0.044, 0.0445, 0.045, 0.0455,<br>0.046, 0.0465, 0.047, 0.0475, 0.048, 0.0485, 0.049, 0.0495, 0.05, 0.0505, 0.051,<br>0.0515, 0.052, 0.0525, 0.053, 0.0535, 0.054, 0.0545, 0.055, 0.0555, 0.056, 0.0565,<br>0.057, 0.0575, 0.058, 0.0585, 0.059, 0.0595, 0.06, 0.0605, 0.061, 0.0615, 0.062,<br>0.0625, 0.063, 0.0635, 0.064, 0.0645, 0.065, 0.0655, 0.066, 0.0665, 0.067, 0.0675,<br>0.068, 0.0685, 0.069, 0.0695, 0.07, 0.0705, 0.071, 0.0715, 0.072, 0.0725, 0.073,<br>0.0735, 0.074, 0.0745, 0.075, 0.0755, 0.076, 0.0765, 0.077, 0.0775, 0.078, 0.0785,<br>0.079, 0.0795, 0.08, 0.0805, 0.081, 0.0815, 0.082, 0.0825, 0.083, 0.0835, 0.084,<br>0.0845, 0.085, 0.0855, 0.086, 0.0865, 0.087, 0.0875, 0.088, 0.0885, 0.089, 0.0895,<br>0.09, 0.0905, 0.091, 0.0915, 0.092, 0.0925, 0.093, 0.0935, 0.094, 0.0945, 0.095,<br>0.0955, 0.096, 0.0965, 0.097, 0.0975, 0.098, 0.0985, 0.099, 0.0995, 0.1, 0.1005,<br>0.101, 0.1015, 0.102, 0.1025, 0.103, 0.1035, 0.104, 0.1045, 0.105, 0.1055, 0.106,<br>0.1065, 0.107, 0.1075, 0.108, 0.1085, 0.109, 0.1095, 0.11, 0.1105, 0.111, 0.1115,<br>0.112, 0.1125, 0.113, 0.1135, 0.114, 0.1145, 0.115, 0.1155, 0.116, 0.1165, 0.117,<br>0.1175, 0.118, 0.1185, 0.119, 0.1195, 0.12, 0.1205, 0.121, 0.1215, 0.122, 0.1225,<br>0.123, 0.1235, 0.124, 0.1245, 0.125, 0.1255, 0.126, 0.1265, 0.127, 0.1275, 0.128,<br>0.1285, 0.129, 0.1295, 0.13, 0.1305, 0.131, 0.1315, 0.132, 0.1325, 0.133, 0.1335,<br>0.134, 0.1345, 0.135, 0.1355, 0.136, 0.1365, 0.137, 0.1375, 0.138, 0.1385, 0.139,<br>0.1395, 0.14, 0.1405, 0.141, 0.1415, 0.142, 0.1425, 0.143, 0.1435, 0.144, 0.1445,<br>0.145, 0.1455, 0.146, 0.1465, 0.147, 0.1475, 0.148, 0.1485, 0.149, 0.1495, 0.15,<br>0.16, 0.17, 0.18, 0.19, 0.2, 0.21, 0.22, 0.23, 0.24, 0.25, 0.26, 0.27, 0.28, 0.29, 0.3,<br>0.31, 0.32, 0.33, 0.34, 0.35, 0.36, 0.37, 0.38, 0.39, 0.4, 0.41, 0.42, 0.43, 0.44, 0.45,<br>0.46, 0.47, 0.48, 0.49, 0.5, 0.51, 0.52, 0.53, 0.54, 0.55, 0.56, 0.57, 0.58, 0.59, 0.6 |

**Table S3:** Parameter spaces for composite-likelihood calculations for independent sweep model simulations when position of selected site varies

|  |  |
| --- | --- |
| Position of selected site | 0, 0.01, 0.02, 0.04, 0.06, 0.08, 0.1, 0.12, 0.14, 0.16, 0.18, 0.2, 0.22, 0.24, 0.26,<br>0.28, 0.3, 0.32, 0.34, 0.36, 0.38, 0.4, 0.42, 0.44, 0.46, 0.48, 0.5, 0.52, 0.54, 0.56, 0.58,<br>0.6, 0.62, 0.64, 0.66, 0.68, 0.7, 0.72, 0.74, 0.76, 0.78, 0.8, 0.82, 0.84, 0.86, 0.88, 0.9,<br>0.92, 0.94, 0.96, 0.98, 1 |
| $s$ | $10^{-4}, 5 \times 10^{-4}, 0.001, 0.002, 0.004, 0.005, 0.006, 0.008, 0.01, 0.012, 0.014,$<br>0.018, 0.02, 0.03, 0.04, 0.05, 0.06, 0.07, 0.09, 0.1, 0.11, 0.12, 0.14, 0.15, 0.2, 0.25,<br>0.3, 0.35, 0.4, 0.5, 0.6 |

**Table S4:** Parameter spaces for composite-likelihood calculations for migration model simulations

|  |  |
| --- | --- |
| Position of selected site | 0 |
| $s$ | $10^{-4}$, 0.001, 0.002, 0.003, 0.004, 0.005, 0.006, 0.007, 0.008, 0.009, 0.01, 0.011, 0.012, 0.013, 0.014, 0.015, 0.016, 0.018, 0.02, 0.022, 0.024, 0.026, 0.028, 0.03, 0.032, 0.034, 0.036, 0.038, 0.04, 0.042, 0.044, 0.046, 0.048, 0.05, 0.052, 0.054, 0.056, 0.058, 0.06, 0.062, 0.064, 0.066, 0.068, 0.07, 0.08, 0.09, 0.1, 0.11, 0.12, 0.13, 0.14, 0.15, 0.2, 0.3, 0.4, 0.5, 0.6 |
| $m$ | $1^{-5}$, $8 \times 10^{-5}$, $0^{-4}$, $1.2 \times 10^{-4}$, $1.4 \times 10^{-4}$, $1.6 \times 10^{-4}$, $1.8 \times 10^{-4}$, $2 \times 10^{-4}$, $2.2 \times 10^{-4}$, $2.4 \times 10^{-4}$, $2.6 \times 10^{-4}$, $2.8 \times 10^{-4}$, $3 \times 10^{-4}$, $3.2 \times 10^{-4}$, $3.4 \times 10^{-4}$, $3.6 \times 10^{-4}$, $3.8 \times 10^{-4}$, $4 \times 10^{-4}$, $8 \times 10^{-4}$, 0.001, 0.0012, 0.0014, 0.0016, 0.0018, 0.002, 0.0022, 0.0024, 0.0026, 0.0028, 0.003, 0.0032, 0.0034, 0.0036, 0.0038, 0.004, 0.006, 0.008, 0.01, 0.012, 0.014, 0.016, 0.036, 0.056, 0.076, 0.096, 0.116, 0.136, 0.156, 0.176, 0.196, 0.3, 0.4, 0.5, 0.6, 0.7, 0.8, 0.9, 1 |
| Migration source population | 2 |

**Table S5:** Parameter spaces for composite-likelihood calculations for standing variation model simulations

|  |  |
| --- | --- |
| Position of selected site | 0 |
| $s$ | $10^{-4}$, 0.0020, 0.0040, 0.0050, 0.0060, 0.0080, 0.0100, 0.0120, 0.0140, 0.0180, 0.0200, 0.0400, 0.0500, 0.0600, 0.0700, 0.0900, 0.1000, 0.1500, 0.2000, 0.3000, 0.4000, 0.5000, 0.6000 |
| $t$ | 5, 5, 25, 40, 50, 60, 75, 100, 150, 200, 250, 300, 350, 400, 450, 500, 550, 600, 650, 700, 750, 800, 900, 1000, 1500, 2000, 2500, 3000, 3500, 4000, 4500, 5000, 5500, 6000, 6500, 7000, 7500, 8000, 9000, 10000, 15000, 20000, 30000, 50000, 70000, 9000, $10^5$ |
| $g$ | $10^{-6}$, $10^{-5}$, $10^{-4}$, $10^{-3}$, $10^{-2}$ |

**Table S6:** Neutral **F** matrix from 12 scaffolds with no strong signatures of selection in *Mimulus guttatus* populations (Scaffold7 and regions adjacent to scaffolds 1, 4, 8, 47, 80, 84, 106, 115, 129, 148, 198). Populations 1 and 3 are copper tolerant.

|  | Pop1 | Pop2 | Pop3 | Pop4 |
| --- | --- | --- | --- | --- |
| Pop1 | 0.1571 | 0.0266 | 0.0153 | 0.0356 |
| Pop2 | 0.0266 | 0.1008 | 0.0000 | 0.0204 |
| Pop3 | 0.0153 | 0.0000 | 0.1807 | 0.0179 |
| Pop4 | 0.0356 | 0.0204 | 0.0179 | 0.1232 |

**Table S7:**
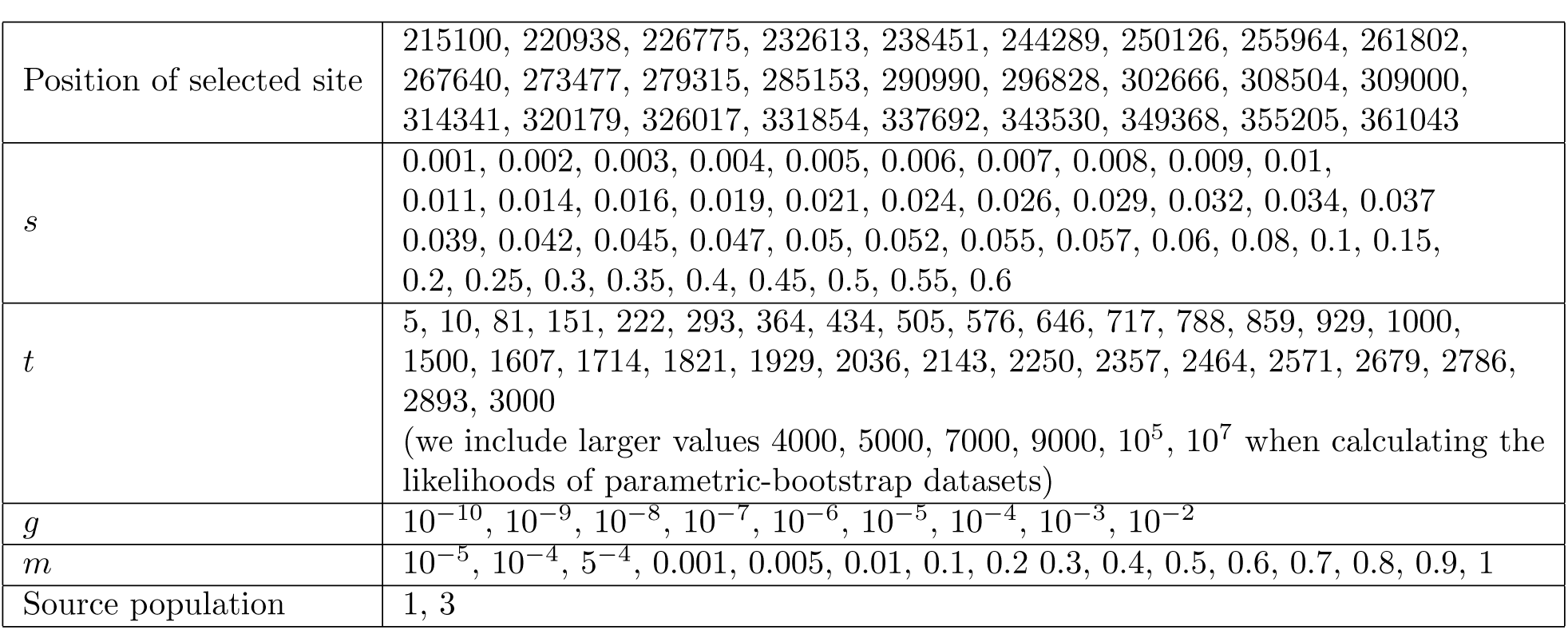
Parameter spaces for composite-likelihood calculations for *Mimulus*

**Table S8:** Parametric-bootstrap results for *Mimulus* analysis

| Model | Range of CLR from 100 simulations<br>(standing source - simulation model) | Observed CLR |
| --- | --- | --- |
| Neutral | [-30.42, 145.04] | 1985.87 |
| Independent mutations | [-0.05, 88.02] | 436.21 |
| Migration | [4.12, 749.45] | 945.95 |

**Table S9:**
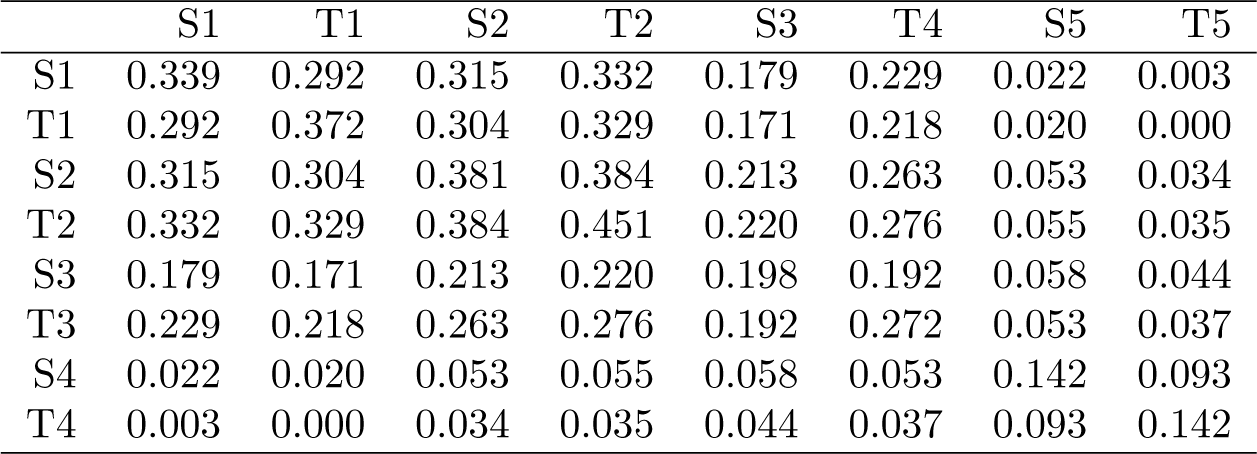
Neutral **F** matrix from four scaffolds with no strong signatures of selection in *Fundulus heteroclitus* populations (Scaffold0, Scaffold1, Scaffold2, Scaffold3)

|  | S1 | T1 | S2 | T2 | S3 | T4 | S5 | T5 |
| --- | --- | --- | --- | --- | --- | --- | --- | --- |
| S1 | 0.339 | 0.292 | 0.315 | 0.332 | 0.179 | 0.229 | 0.022 | 0.003 |
| T1 | 0.292 | 0.372 | 0.304 | 0.329 | 0.171 | 0.218 | 0.020 | 0.000 |
| S2 | 0.315 | 0.304 | 0.381 | 0.384 | 0.213 | 0.263 | 0.053 | 0.034 |
| T2 | 0.332 | 0.329 | 0.384 | 0.451 | 0.220 | 0.276 | 0.055 | 0.035 |
| S3 | 0.179 | 0.171 | 0.213 | 0.220 | 0.198 | 0.192 | 0.058 | 0.044 |
| T3 | 0.229 | 0.218 | 0.263 | 0.276 | 0.192 | 0.272 | 0.053 | 0.037 |
| S4 | 0.022 | 0.020 | 0.053 | 0.055 | 0.058 | 0.053 | 0.142 | 0.093 |
| T4 | 0.003 | 0.000 | 0.034 | 0.035 | 0.044 | 0.037 | 0.093 | 0.142 |

**Table S10:**
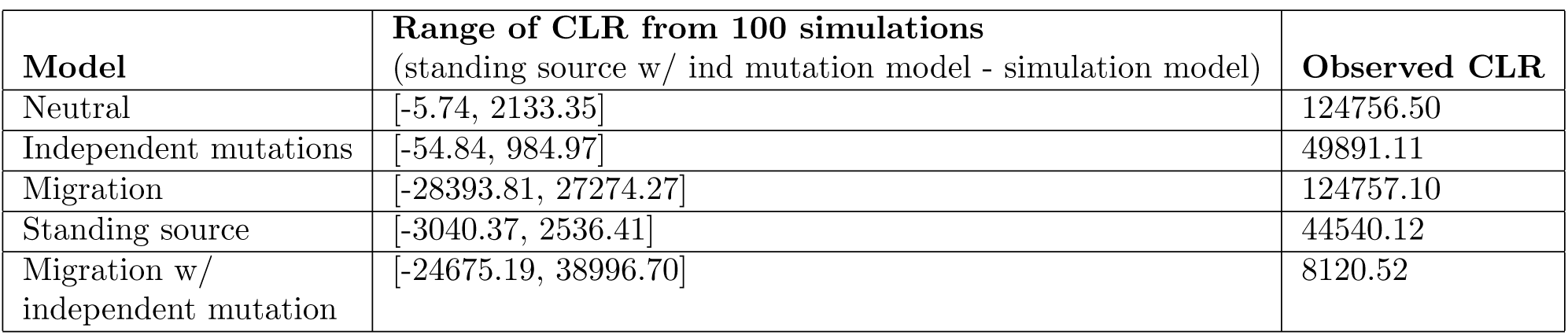
Parameter spaces for composite-likelihood calculations for *Fundulus*

**Table S11:**
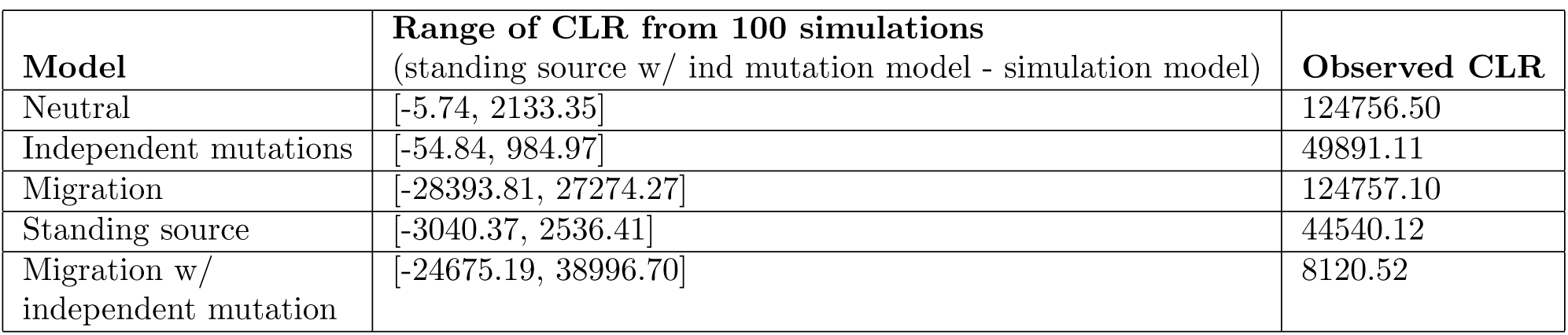
Parametric-bootstrap results for *Fundulus* analysis

## References

Arendt, J. and D. Reznick (2008). Convergence and parallelism reconsidered: what have we learned about the genetics of adaptation? Trends in Ecology and Evolution 23(1), 26–32.

Aubury, L. E. (1902). The copper resources of California (No. 23). Superintendent State Printing.

Barrett, R. D. and D. Schluter (2008). Adaptation from standing genetic variation. Trends in ecology & evolution 23(1), 38–44.

Barton, N. (1998). The effect of hitch-hiking on neutral genealogies. Genet. Res. 72, 123—133.

Barton, N. and B. O. Bengtsson (1986, Dec). The barrier to genetic exchange between hybridising populations. Heredity 57(3), 357–376.

Berg, J. J. and G. Coop (2015). A coalescent model for a sweep of a unique standing variant. Genetics 201(2), 707–725.

Bierne, N. (2010). The distinctive footprints of local hitchhiking in a varied environment and global hitchhiking in a subdivided population. Evolution 64(11), 3254–3272.

Bierne, N., P.-A. Gagnaire, and P. David (2013). The geography of introgression in a patchy environment and the thorn in the side of ecological speciation. Current Zoology 59(1), 72–86.

Chan, Y. F., M. E. Marks, F. C. Jones, G. Villarreal, M. D. Shapiro, S. D. Brady, A. M. Southwick, D. M. Absher, J. Grimwood, J. Schmutz, et al. (2010). Adaptive evolution of pelvic reduction in sticklebacks by recurrent deletion of a pitx1 enhancer. science 327(5963), 302–305.

Charlesworth, B., M. Nordborg, and D. Charlesworth (1997, Oct). The effects of local selection, balanced polymorphism and background selection on equilibrium patterns of genetic diversity in subdivided populations. GenetRes 70(2), 155–174.

Chen, H., N. Patterson, and D. Reich (2010). Population differentiation as a test for selective sweeps. Genome research 20(3), 393–402.

Colosimo, P. F., K. E. Hosemann, S. Balabhadra, G. Villarreal, M. Dickson, J. Grimwood, J. Schmutz, R. M. Myers, D. Schluter, and D. M. Kingsley (2005). Widespread parallel evolution in sticklebacks by repeated fixation of ectodysplasin alleles. science 307(5717), 1928–1933.

Coop, G., D. Witonsky, A. Di Rienzo, and J. K. Pritchard (2010). Using environmental correlations to identify loci underlying local adaptation. Genetics 185(4), 1411–1423.

DeGiorgio, M., K. E. Lohmueller, and R. Nielsen (2014). A model-based approach for identifying signatures of ancient balancing selection in genetic data. PLoS Genet 10(8), e1004561.

Durrett, R. and J. Schweinsberg (2004, 10). Approximating selective sweeps. 66, 129–38.

Duvernell, D. D., J. B. Lindmeier, K. E. Faust, and A. Whitehead (2008). Relative influences of historical and contemporary forces shaping the distribution of genetic variation in the atlantic killifish, fundulus heteroclitus. Molecular Ecology 17(5), 1344–1360.

Ewens, W. (2004). *Mathematical Population Genetics 1: Theoretical Introduction*. Interdisciplinary Applied Mathematics. Springer New York.

Gillespie, J. H. (2000). Genetic drift in an infinite population. The pseudohitchhiking model. Genetics 155, 909–919.

Guerrero, R. F., F. Rousset, and M. Kirkpatrick (2012). Coalescent patterns for chromosomal inversions in divergent populations. Phil. Trans. R. Soc. B 367(1587), 430–438.

Harvey, P. H. and M. D. Pagel (1991). The comparative method in evolutionary biology, Volume 239. Oxford university press Oxford.

Hedrick, P. W. (2013). Adaptive introgression in animals: examples and comparison to new mutation and standing variation as sources of adaptive variation. Molecular ecology 22(18), 4606–4618.

Heliconius Genome Consortium (2012). Butterfly genome reveals promiscuous exchange of mimicry adaptations among species. Nature 487(7405), 94–98.

Hudson, R. R. (2001). Two-locus sampling distributions and their application. Genetics 159(4), 1805–1817.

Hudson, R. R. (2002). Generating samples under a Wright–Fisher neutral model of genetic variation. Bioinformatics 18, 337–338.

Hudson, R. R. and N. L. Kaplan (1988). The coalescent process in models with selection and recombination. Genetics 120(3), 831–840.

Jones, F. C., M. G. Grabherr, Y. F. Chan, P. Russell, E. Mauceli, J. Johnson, R. Swofford, M. Pirun,M. C. Zody, S. White, E. Birney, S. Searle, J. Schmutz, J. Grimwood, M. C. Dickson, R. M. Myers, C. T. Miller, B. R. Summers, A. K. Knecht, S. D. Brady, H. Zhang, A. A. Pollen, T. Howes, C. Amemiya, J. Baldwin, T. Bloom, D. B. Jaffe, R. Nicol, J. Wilkinson, E. S. Lander, F. Di Palma, K. Lindblad-Toh, and D. M. Kingsley (2012, Apr). The genomic basis of adaptive evolution in threespine sticklebacks. Nature 484(7392), 55–61.

Kaplan, N., R. R. Hudson, and M. Iizuka (1991). The coalescent process in models with selection, recombination and geographic subdivision. Genetical research 57(01), 83–91.

Kaplan, N. L., R. R. Hudson, and C. H. Langley (1989, December). The “hitchhiking effect” revisited. Genetics 123, 887–899.

Kelly, J. K. and M. J. Wade (2000). Molecular evolution near a two-locus balanced polymorphism. Journal of Theoretical Biology 204(1), 83–101.

Kim, Y. and T. Maruki (2011). Hitchhiking effect of a beneficial mutation spreading in a subdivided population. Genetics 189(1), 213–226.

Kim, Y. and R. Nielsen (2004). Linkage disequilibrium as a signature of selective sweeps. Genetics 167(3), 1513–1524.

Kim, Y. and W. Stephan (2002). Detecting a local signature of genetic hitchhiking along a recombining chromosome. Genetics 160, 765–777.

Kirkpatrick, M. (2010). How and why chromosome inversions evolve. PLoS Biol 8(9), e1000501.

Larribe, F. and P. Fearnhead (2011). On composite likelihoods in statistical genetics. Statistica Sinica, 43–69.

Lee, Y. W. (2009). Genetics Analysis of Standing Variation for Floral Morphology and Fitness Components. Ph. D. thesis, Duke University.

Lipson, M., P.-R. Loh, A. Levin, D. Reich, N. Patterson, and B. Berger (2013, May). Efficient moment-based inference of admixture parameters and sources of gene flow. Mol Biol Evol 30(8), 1788–1802.

Losos, J. B. (2011). Convergence, adaptation, and constraint. Evolution 65(7), 1827–1840.

MacNair, M. R., S. E. Smith, and Q. J. Cumbes (1993, 11). Heritability and distribution of variation in degree of copper tolerance in mimulus guttatus at copperopolis, california. Heredity 71(5), 445–455.

Martin, A. and V. Orgogozo (2013). The loci of repeated evolution: a catalog of genetic hotspots of phenotypic variation. Evolution 67(5), 1235–1250.

Maynard Smith, J. (1971). What use is sex? Journal of Theoretical Biology 30(2), 319–335.

Maynard Smith, J. and J. Haigh (1974, February). The hitch-hiking effect of a favourable gene. Genet Res 23(1), 23–35.

Nacci, D., L. Coiro, D. Champlin, S. Jayaraman, R. McKinney, T. R. Gleason, W. R. Munns Jr., J. L. Specker, and K. R. Cooper (1999). Adaptations of wild populations of the estuarine fish fundulus heteroclitus to persistent environmental contaminants. Marine Biology 134(1), 9–17.

Nacci, D. E., D. Champlin, and S. Jayaraman (2010). Adaptation of the estuarine fish fundulus heteroclitus (atlantic killifish) to polychlorinated biphenyls (pcbs). Estuaries and Coasts 33(4), 853–864.

Nicholson, G., A. V. Smith, F. JÓnsson, Ó. Gústafsson, K. Stefánsson, and P. Donnelly (2002). Assessing population differentiation and isolation from single-nucleotide polymorphism data. Journal of the Royal Statistical Society: Series B (Statistical Methodology) 64(4), 695–715.

Nielsen, R., S. Williamson, Y. Kim, M. Hubisz, A. Clark, and C. Bustamante (2005). Genomic scans for selective sweeps using SNP data. Genome Res. 15, 1566–1575.

Orr, H. A. (2005, January). The probability of parallel evolution. Evolution 59(1), 216–220.

Pearce, R. J., H. Pota, M.-S. B. Evehe, E.-H. Bâ, G. Mombo-Ngoma, A. L. Malisa, R. Ord, W. Inojosa,A. Matondo, D. A. Diallo, W. Mbacham, I. V. van den Broek, T. D. Swarthout, A. Getachew, S. Dejene,M. P. Grobusch, F. Njie, S. Dunyo, M. Kweku, S. Owusu-Agyei, D. Chandramohan, M. Bonnet, J.-P. Guthmann, S. Clarke, K. I. Barnes, E. Streat, S. T. Katokele, P. Uusiku, C. O. Agboghoroma, O. Y. Elegba, B. Cissé, I. E. A-Elbasit, H. A. Giha, S. P. Kachur, C. Lynch, J. B. Rwakimari, P. Chanda, M. Hawela, B. Sharp, I. Naidoo, and C. Roper (2009, 04). Multiple origins and regional dispersal of resistant dhpsin African Plasmodium falciparummalaria. PLoS Med 6(4), e1000055.

Pease, J. B., D. C. Haak, M. W. Hahn, and L. C. Moyle (2016). Phylogenomics reveals three sources of adaptive variation during a rapid radiation. PLoS Biol 14(2), e1002379.

Pennings, P. S. and J. Hermisson (2006, May). Soft sweeps ii—molecular population genetics of adaptation from recurrent mutation or migration. Mol Biol Evol 23(5), 1076–1084.

Przeworski, M., G. Coop, and J. D. Wall (2005). The signature of positive selection on standing genetic variation. Evolution 59(11), 2312–2323.

Racimo, F. (2016). Testing for ancient selection using cross-population allele frequency differentiation. Genetics 202(2), 733–750.

Racimo, F., S. Sankararaman, R. Nielsen, and E. Huerta-Sánchez (2015). Evidence for archaic adaptive introgression in humans. Nature Reviews Genetics 16(6), 359–371.

Reid, N. M., D. A. Proestou, B. W. Clark, W. C. Warren, J. K. Colbourne, J. R. Shaw, S. I. Karchner, M. E. Hahn, D. Nacci, M. F. Oleksiak, D. L. Crawford, and A. Whitehead (2016). The genomic landscape of rapid repeated evolutionary adaptation to toxic pollution in wild fish. Science 354(6317), 1305–1308.

Roesti, M., S. Gavrilets, A. P. Hendry, W. Salzburger, and D. Berner (2014). The genomic signature of parallel adaptation from shared genetic variation. Molecular ecology 23(16), 3944–3956.

Rosenzweig, B. K., J. B. Pease, N. J. Besansky, and M. W. Hahn (2016). Powerful methods for detecting introgressed regions from population genomic data. Molecular Ecology 25(11), 2387–2397.

Samanta, S., Y.-J. Li, and B. S. Weir (2009). Drawing inferences about the coancestry coefficient. Theoretical Population Biology 75(4), 312–9.

Santiago, E. and A. Caballero (2005). Variation after a selective sweep in a subdivided population. Genetics 169, 475–483.

Schluter, D. and G. L. Conte (2009, Jun). Genetics and ecological speciation. Proc Natl Acad Sci U S A 106 Suppl 1, 9955–9962.

Slatkin, M. and T. Wiehe (1998). Genetic hitch-hiking in a subdivided population. Genetical research 71(02), 155–160.

Song, Y., S. Endepols, N. Klemann, D. Richter, F.-R. Matuschka, C.-H. Shih, M. W. Nachman, and M. H. Kohn (2011). Adaptive introgression of anticoagulant rodent poison resistance by hybridization between old world mice. Current Biology 21(15), 1296–1301.

Stern, D. L. (2013, 11). The genetic causes of convergent evolution. Nat Rev Genet 14(11), 751–764.

Thompson, E. A. (2013). Identity by descent: Variation in meiosis, across genomes, and in populations. Genetics 194(2), 301–326.

Tishkoff, S. A., F. A. Reed, A. Ranciaro, B. F. Voight, C. C. Babbitt, J. S. Silverman, K. Powell, H. M. Mortensen, J. B. Hirbo, M. Osman, M. Ibrahim, S. A. Omar, G. Lema, T. B. Nyambo, J. Ghori, S. Bump- stead, J. K. Pritchard, G. A. Wray, and P. Deloukas (2007, Jan). Convergent adaptation of human lactase persistence in Africa and Europe. Nat Genet 39(1), 31–40.

Turner, T., E. Bourne, E. V. Wettberg, T. Hu, and S. Nuzhdin (2010, January). Population resequencing reveals local adaptation of *Arabidopsis lyrata*to serpentine soils. Nat. Genet. 42(3), 260–3.

Varin, C., N. Reid, and D. Firth (2011). An overview of composite likelihood methods. Statistica Sinica, 5–42.

Weir, B. S. and W. G. Hill (2002). Estimating F-statistics. Annual Review of Genetics 36, 721–50.

Welch, J. J. and C. D. Jiggins (2014). Standing and flowing: the complex origins of adaptive variation. Molecular Ecology 23(16), 3935–3937.

Wiuf, C. (2006). Consistency of estimators of population scaled parameters using composite likelihood. Journal of mathematical biology 53(5), 821–841.

Wood, T. E., J. M. Burke, and L. H. Rieseberg (2005, 02). Parallel genotypic adaptation: when evolution repeats itself. Genetica 123(1-2), 157–170.

Wright, K. M., U. Hellsten, C. Xu, A. L. Jeong, A. Sreedasyam, J. A. Chapman, J. Schmutz, G. Coop, D. S. Rokhsar, and J. H. Willis (2015). Adaptation to heavy-metal contaminated environments proceeds via selection on pre-existing genetic variation. bioRxiv.

Wright, S. (1943). Isolation by distance. Genetics 28(2), 114.

Wright, S. (1951, March). The genetical structure of populations. Annals of Eugenics 15(4), 323–354.

